# Integration of Patient-Derived Organoids and Organ-on-Chip Systems: Investigating Colorectal Cancer Invasion within the Mechanical and GABAergic Tumor Microenvironment

**DOI:** 10.1101/2023.09.14.557797

**Authors:** Carly Strelez, Rachel Perez, John S. Chlystek, Christopher Cherry, Ah Young Yoon, Bethany Haliday, Curran Shah, Kimya Ghaffarian, Ren X. Sun, Hannah Jiang, Roy Lau, Aaron Schatz, Heinz-Josef Lenz, Jonathan E. Katz, Shannon M. Mumenthaler

## Abstract

Three-dimensional (3D) in vitro models are essential in cancer research, but they often neglect physical forces. In our study, we combined patient-derived tumor organoids with a microfluidic organ-on-chip system to investigate colorectal cancer (CRC) invasion in the tumor microenvironment (TME). This allowed us to create patient-specific tumor models and assess the impact of physical forces on cancer biology. Our findings showed that the organoid-on-chip models more closely resembled patient tumors at the transcriptional level, surpassing organoids alone. Using ’omics’ methods and live-cell imaging, we observed heightened responsiveness of KRAS mutant tumors to TME mechanical forces. These tumors also utilized the γ-aminobutyric acid (GABA) neurotransmitter as an energy source, increasing their invasiveness. This bioengineered model holds promise for advancing our understanding of cancer progression and improving CRC treatments.

**Highlights:**

- Microfluidic organ-on-chip system integrated with patient-derived CRC organoids
- Physical forces influence invasion, particularly in KRAS mutant tumor cells
- GABAergic signaling contributes to increased invasion within a dynamic TME
- This model explores patient heterogeneity, TME interactions, and cancer progression

**GRAPHICAL ABSTRACT:** 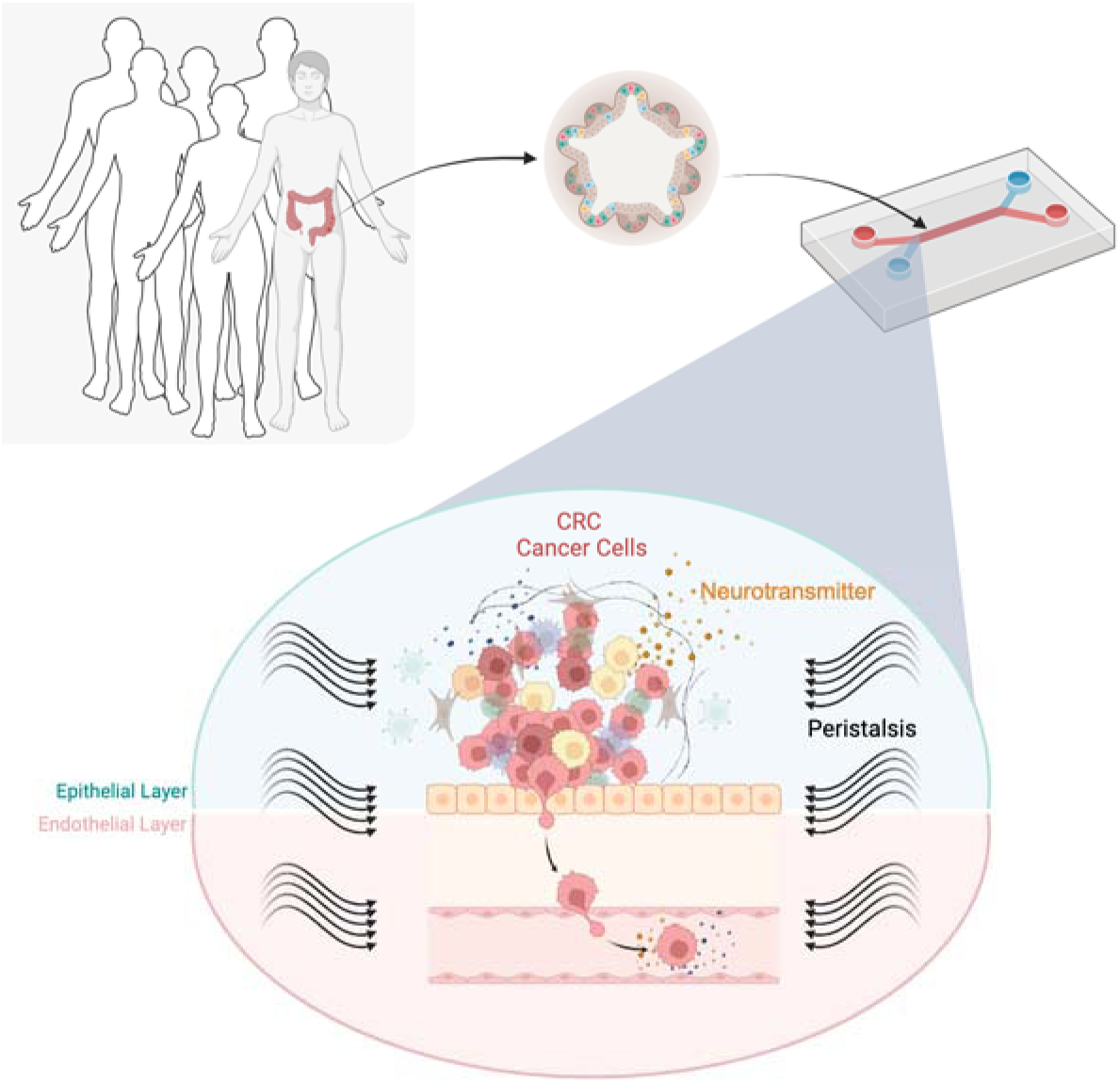

## INTRODUCTION

Cancer, a complex and multifaceted disease, continues to challenge our understanding and treatment efforts. While remarkable strides have been made in elucidating the genetic underpinnings of various malignancies^1, 2^, there remains a critical need to comprehensively explore the tumor microenvironment (TME) and its intricate interplay with cancer progression. In this manuscript, we explore a promising convergence of two advanced technologies: organoids and organ-on-chip (OOC) systems. This integration represents a notable advancement in our ongoing efforts to better understand the complex mechanisms underlying cancer metastasis.

The emergence of organoid-on-chip platforms represents a paradigm shift in our ability to recapitulate the intricacies of the TME *in vitro*^3^. This approach builds upon 3D cell culture techniques, going beyond simplistic monolayer cultures to create a more sophisticated three-dimensional environment^4, 5^. Patient-derived organoids have been instrumental in precision medicine, contributing significantly to various aspects of cancer research, from mechanistic studies to drug discovery applications^6, 7^. One key feature of this innovation is its patient-specific nature, allowing us to replicate individual tumor characteristics and heterogeneity accurately^8^. This aspect is crucial for bridging the gap between laboratory experiments and clinical relevance. However, despite the substantial advancements in organoid technology, limitations persist, particularly in the context of studying cancer metastasis. These limitations stem primarily from the absence of crucial components such as the vasculature and biophysical properties. Microfluidic organ-on-chip technology emerges as a promising solution, offering a more comprehensive platform for investigating normal organ development and disease pathologies, including cancer, by integrating vascular networks^9–11^ and simulating biochemical and biophysical conditions^12, 13^. Specifically, tumor-on-chip models have been invaluable for exploring the complexities of the TME and its interactions, driving advancements in tumor modeling, drug screening, personalized medicine, and immunotherapy approaches^14–18^.

Traditionally, our understanding of cancer metastasis has predominantly focused on genetic and biochemical factors. While these aspects remain crucial, the recognition of biomechanical elements, such as how mechanical forces affect cancer cell behavior, has grown in recent years. Biophysical forces encompass a range of mechanical factors, including shear forces such as fluid flow, compression, and tension, among others. In the context of tumor biology, these forces play a pivotal role in shaping cancer progression^19–21^ by influencing the ability of cancer cells to proliferate, invade surrounding tissues, intravasate into blood or lymphatic vessels, and extravasate at distant sites^22, 23^. Moreover, biophysical cues from the TME can impact cell signaling, gene expression, and even therapeutic responses^12, 24^. Recent studies suggest that alterations in neurotransmitter signaling within the gut may contribute to the initiation and progression of colorectal cancer (CRC)^25, 26^. Dysregulation of serotonin, for instance, has been linked to changes in gut motility and inflammation, which can create an environment conducive to CRC development^27, 28^. By integrating biophysical forces into our models, we can now gain deeper insights into how these forces influence tumor cell invasion and dissemination.

In this manuscript, we present a comprehensive exploration of how engineered cancer models, which combine the strengths of organoids and organ-on-chip technology, advance our understanding of cancer metastasis, biophysical forces, and neurotransmitter signaling in the context of CRC. We not only provide a valuable bioengineered model as a resource for the cancer research community but also demonstrate the effectiveness of our approach through a proof-of-concept study.

## RESULTS

### Development of tumor organoid-on-chip to model colorectal cancer

To create a patient-derived CRC organoid-on-chip, we implemented the following workflow: (1) procured fresh CRC tumor tissues from both primary and metastatic locations, (2) cultivated and biobanked tumor organoids, and (3) optimized the process for seeding and culturing organoids within the microfluidic OOC device with cyclic strain to mimic peristalsis, as illustrated in **Figure 1A**. We adapted Emulate’s OOC technology, previously employed to simulate the functional aspects of normal organs such as the intestine^29^. Tumor organoids were introduced into the top channel, while human intestinal microvascular endothelial cells (HIMECs) were seeded into the bottom channel of a microfluidic chip, separated by a porous membrane. To prepare the channels, their surfaces were first activated and coated with extracellular matrix (ECM). The top channel was lined with Matrigel, while collagen IV and fibronectin coated the bottom channel. Approximately one day after seeding the tumor organoids and HIMECs, we initiated flow at a specific rate of 30 µL hr^-1^. To mimic physiological peristalsis, we introduced mechanical stretching at 10% deformation and 0.2 Hz approximately 48 hr later. The values for peristalsis were selected based on clinical measurements of the intestine and previous work performed in normal intestine-chips^29, 30^. These experiments continued for an additional 6 days, as depicted in the experimental timeline in **Figure S1**.

**Figure 1.**
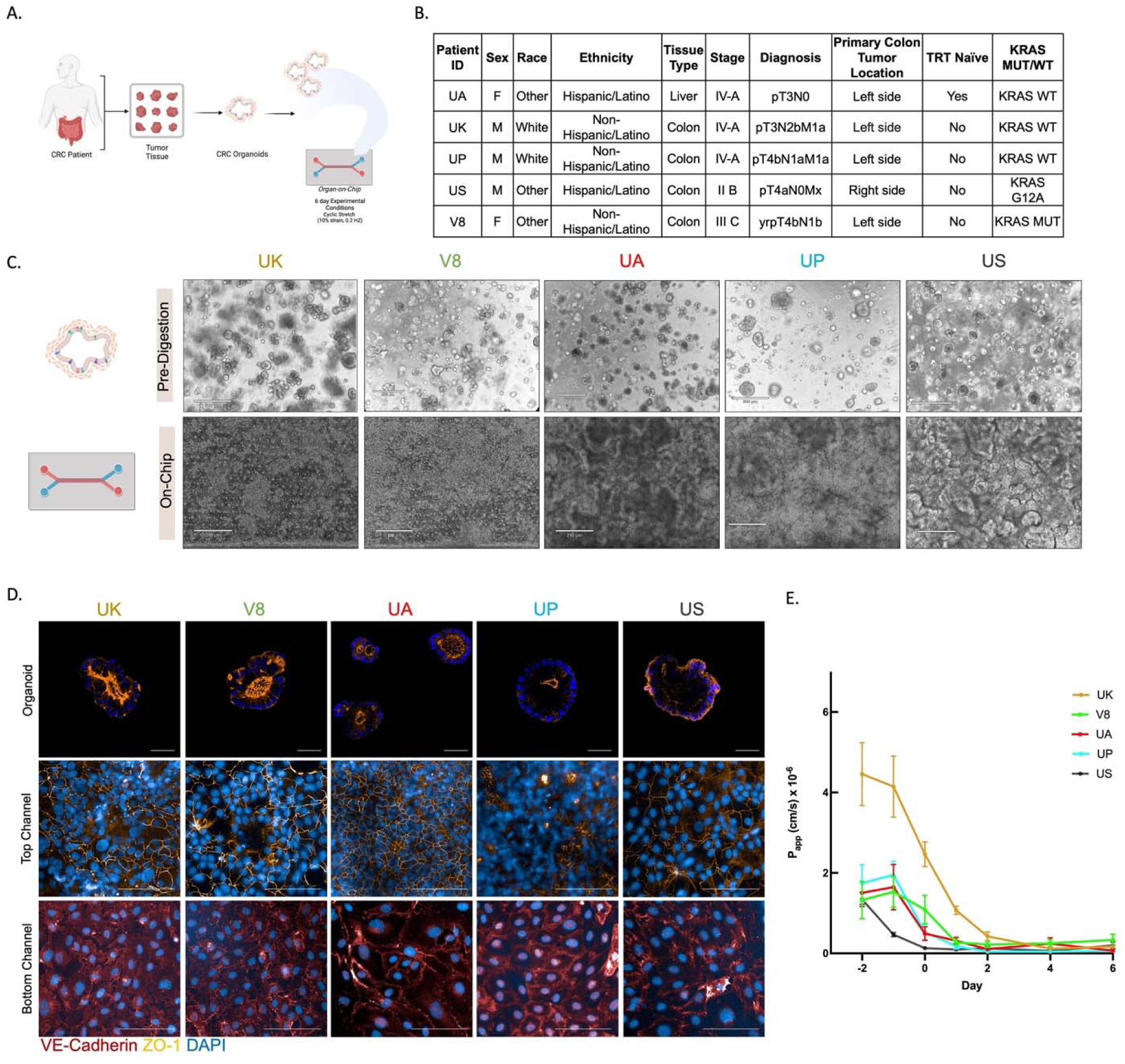
Patient-derived organoid-on-chip to model colorectal cancer. **(A.)**Schematic of organoid-on-chip development. Colorectal cancer (CRC) organoids were derived from patient tumor tissues. Organoids were then seeded into the top channel of the OOC and cultured with fluid flow and rhythmic mechanical strain to mimic peristalsis for 6 days. Schematic was made using BioRender. **(B.)** Summary of clinical details for each de-identified patient sample, including gender and ethnicity, tumor staging, location, and mutational status. **(C.)** Representative brightfield images of patient-derived CRC organoids in culture prior to digestion (top panel) and after 6 days on-chip (bottom panel) showing the morphology differences between patients. Scale bars for the pre-digested organoids represent 500 μm and scale bars for the organoids on-chip represent 210 μm. **(D.)** Representative confocal images of organoid-only cultures (top panel), epithelial top channel (middle panel), and endothelial bottom channel (bottom panel) of organoids-on-chips immunostained for ZO-1 (gold) and VE-Cadherin (red) on day 6. DAPI (blue) labels the cell nuclei. Scale bars in the organoid and organoid-on-chip images represent 50 and 100μm, respectively. **(E.)** The apparent permeability (P_app_) of the CRC organoid-on-chips cultured in the presence of flow (30 μL/hr) and stretch (10% strain, 0.2 Hz) over the course of the experiment. P_app_ values were calculated from the concentration of 3kDa Dextran that diffused from the epithelial channel to the endothelial channel. N=6-7 chips per patient. Data are represented as mean ± SEM.

We selected a diverse range of tumor organoids from our biorepository, ensuring representation across various factors such as sex, race/ethnicity, cancer stage, and mutational status (as shown in**Figure 1B**). The selected tumors encompassed stages IIB through IV-A and included two with KRAS mutations and three with wild-type KRAS, reflecting the prevailing rates of KRAS mutations in CRC patient populations^31^. These tumor organoids exhibited varying morphologies across different patients, as depicted in **Figure 1C** (top panel), and this diversity was also observed in the on-chip culture setting (**Figure 1C**, bottom panel). To assess the integrity of permeability barriers, we employed immunofluorescence staining of zonula occludens-1 (ZO-1) in both organoid-only cultures and organoids-on-chip (**Figure 1D** and **Figure S2**). Notably, each organoid displayed robust ZO-1 expression, although the specific patterns were unique to each patient. Additionally, we visualized endothelial adherent junctions through immunofluorescence staining of VE-cadherin. Across all donors, HIMECs consistently exhibited strong VE-cadherin expression, as depicted in **Figure 1D** (bottom panel) and **Figure S2**. Moreover, the culture conditions on the microfluidic chip, incorporating both flow and stretching, led to the formation of a robust intestinal barrier by Day 0, which was consistently maintained for over one week, as indicated in **Figure 1E**. While there was notable variability among donors (particularly prior to Day 0), it is reasonable to attribute this variation to differences in the growth rate behaviors of individual organoids, as illustrated in **Figure 1C**.

### Characterizing heterogeneity within and across model systems

We aimed to assess the degree of similarity between CRC organoid-on-chips and patient-matched tissue in comparison to independently cultured organoids. We achieved this by conducting thorough ’omics’ and functional evaluations, as illustrated in **Figure 2A**. Our approach involved conducting bulk RNA-seq analysis on three distinct patient-matched sample groups: (1) tumor tissue, (2) organoids cultured using conventional cell culture methods, and (3) organoids harvested from the epithelial channel of the CRC organoid-on-chip. We then assessed the number of differentially expressed genes (p<0.05) in three key comparisons: organoid alone vs tumor tissue, organoid-on-chip vs tumor tissue, and organoid-on-chip vs organoid alone, involving a cohort of 5 patients. Our analysis revealed 5716 genes with significant expression differences between organoids and tumor tissue, while 4168 genes exhibited significant distinctions between the organoid-on-chip and tumor tissue (as illustrated in **Figure 2B**). The reduced number of differentially expressed genes in the organoid-on-chip vs tumor comparison suggests that the organoid-on-chip model demonstrates a higher degree of transcriptomic similarity to the tumor tissue compared to the organoid model. Furthermore, to gain insights into variations in Hallmark pathways between these model systems, we performed a gene set enrichment analysis (GSEA). Notably, the organoid vs tumor comparison demonstrated more significant enrichment in several pathways compared to the organoid-on-chip vs tumor comparison, further supporting the notion that the organoid-on-chip more closely mirrors the tumor in comparison to organoids alone (see **Table S1** for details).

**Figure 2.**
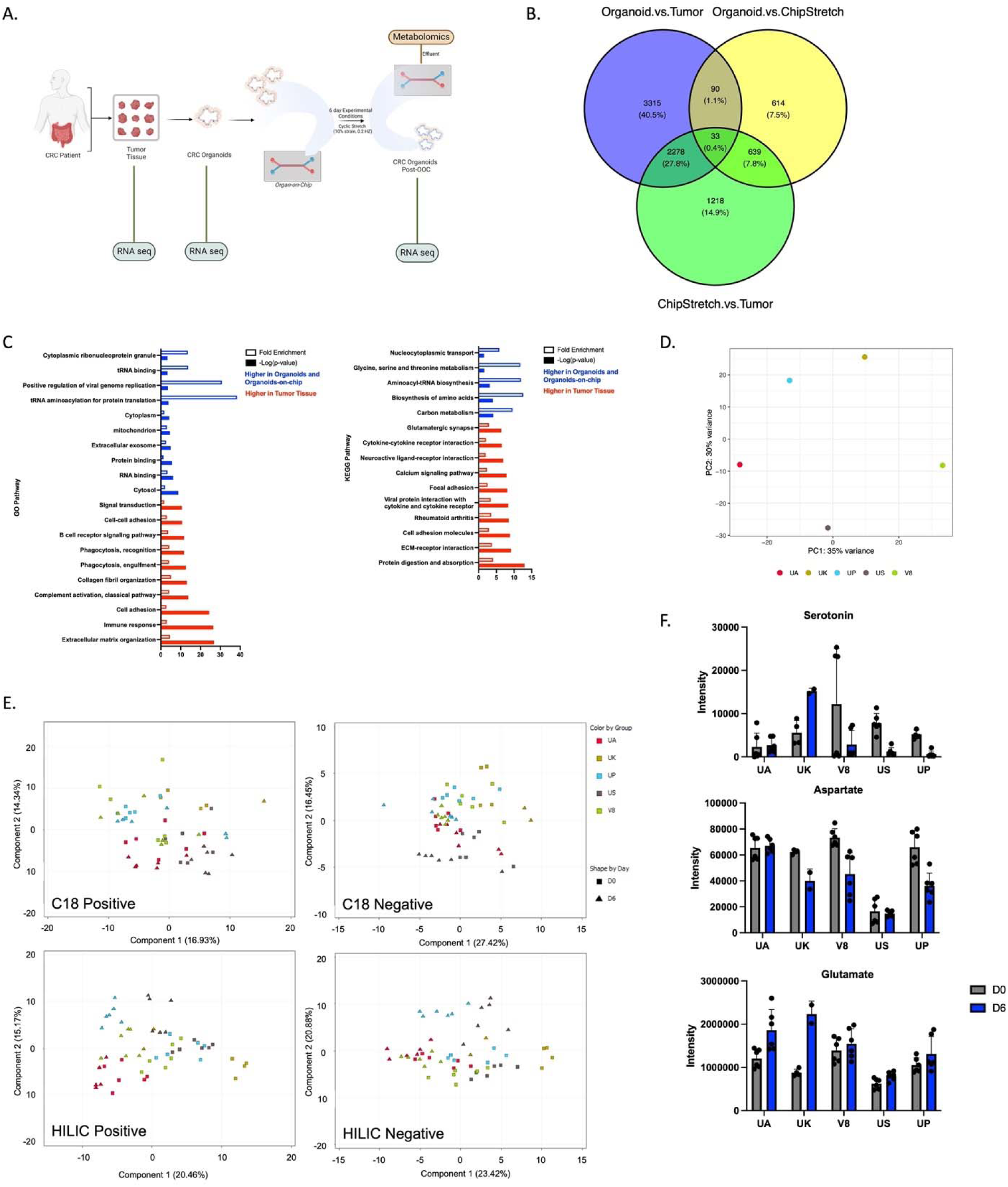
OOC reproduces patient heterogeneity and the neurotransmitter influenced TME. **(A.)** To characterize the OOC model system, effluent was collected at various time points during the on-chip experiment and mass spectrometry-based metabolomics was performed. In addition, RNA-seq analysis was performed on the tumor tissue, organoids prior to OOC seeding, and at the end of the on-chip experiment to compare across the model systems. Schematic was made using BioRender. **(B.)** Differential gene expression analysis was carried out to identify genes that are up or down-regulated (p<0.05) in the CRC organoid-on-chip compared to the organoids alone (yellow), CRC organoid-on-chip with fluid flow and rhythmic stretching compared to tumor (green), or the organoid compared to the tumor (blue). The numbers of unique and overlapping genes were identified and represented in a Venn Diagram. N = 5 independent donors and organoid, organoid-on-chip, and tumor tissue were patient-matched. **(C.)** Genes that were upregulated in organoids and organoid-on-chips as compared to the tumor tissue (blue) and genes that were upregulated in the tumor tissue as compared to organoids and organoid-on-chips (red) were subjected to over-enrichment analysis using DAVID software. **(D.)** Principal component analysis (PCA) of the RNA-seq of the CRC organoid-on-chips demonstrates the patient heterogeneity represented in the genes. Each dot represents one replicate of stretched, CRC organoid-on-chips and each dot is colored by donor.**(E.)** Epithelial effluent was collected from the stretched, CRC organoid-on-chips on days 0 and 6 of the experiment and mass spectrometry-based metabolomics was performed. The PCA on the differential metabolites demonstrates the clustering of samples corresponding to the different patients and the different time points. Results are reported for the mode chromatography/ionization mode that was found to be the most robust for the reported analyte. N=6 chips per timepoint per patient; N=4 on D0 and 2 on D6 for UK. **(F.)** To investigate the role of neurotransmitters in CRC, epithelial effluent was analyzed using a neurotransmitter-specific library. The intensities of several neurotransmitters are shown over time for each patient. N=6 chips per timepoint per patient; n=4 on D0 and 2 on D6 for UK. Individual data points are shown and mean ± SEM are represented.

To identify the specific attributes of the tumor tissue that were not effectively replicated in either the organoids or the organoid-on-chips, we conducted an over-enrichment analysis of the up-or down-regulated genes within both the organoid vs tumor and organoid-on-chip vs tumor comparisons, as depicted in **Figure 2C**. The genes exhibiting higher expression levels in both organoids and organoid-on-chips (indicated in blue) demonstrated enrichment in processes related to protein synthesis and metabolism. These enrichments were observed in both Gene Ontology (GO) biological process (BP) pathways and the Kyoto Encyclopedia of Genes and Genomes (KEGG) pathways. Conversely, genes with elevated expression levels in the tumor tissues (highlighted in red) displayed enrichment in ECM components and organization, cell adhesion, and various aspects of the immune system.

Considering the diverse selection of CRC patient samples used in developing our model and the morphological differences observed within our organoids-on-chip cohort, we conducted a thorough investigation into the apparent inter-patient heterogeneity within the organoid-on-chips. Our approach began with a comprehensive examination of transcriptome-wide expression changes across the five donor-derived OOC models, utilizing principal component analysis (PCA). This analysis revealed significant variance among patients, as illustrated in **Figure 2D**. Additionally, we conducted an in-depth exploration of metabolite changes within the organoid-on-chips by employing mass spectrometry-based metabolomics and examining effluent samples at various time points. The PCA analysis of metabolite intensities revealed distinct separation patterns based on both patient identity (indicated by symbol color) and the experimental day (indicated by symbol shape), as depicted in **Figure 2E** (**Tables S2** and **S3**).

Given the significant strengths of our model in replicating key aspects of the physical TME, such as its microfluidic capabilities and the influence of peristaltic forces, coupled with the growing acknowledgement of the gut-brain axis in cancer research^32, 33^, we were motivated to explore the intricate relationship between neurotransmitter signaling and colon cancer. We examined neurotransmitter levels in the effluent of our chips using a neurotransmitter-specific library. Notably, we identified alterations in several neurotransmitters over varying time periods and across different patients, suggesting that the role of neurotransmitters requires further study and that our model may be a good system to interrogate these changes (**Figure 2F**).

### Peristalsis-like physical forces drive invasive capabilities in KRAS mutant tumor cells

Mechanical forces, such as shear stress from fluid flow or cyclic strain induced by peristalsis, are known to play a crucial role in directing normal intestinal function^34, 35^. However, their influence within the context of CRC biology remains less understood. To address this, we investigated gene expression differences between organoid-on-chips subjected to peristalsis-like motions (10% deformation, 0.2 Hz) and those maintained under static conditions (with no deformation), as represented schematically in **Figure 3A**. While we did not observe any genes with significantly different expression levels below an adjusted p-value threshold of 0.05, GSEA revealed significant differences in key Hallmark gene sets between the stretched and not stretched conditions. Specifically, we found that the gene sets associated with epithelial-mesenchymal transition (EMT), oxidative phosphorylation (oxphos), and fatty acid metabolism exhibited substantial distinctions (adjusted p-value = 0.00367, normalized enrichment score (NES) = 1.65; adjusted p-value = 5.1E-06, NES = -1.92; adjusted p-value = 0.039, NES = -1.48, respectively). These findings indicate that stretching promotes tumor cell gene expression toward EMT while reducing oxphos and fatty acid metabolism. Interestingly, the KRAS signaling Hallmark pathway indicated a trend towards significance (adjusted p-value = 0.062, NES = 1.2) (**Table S4**).

**Figure 3.**
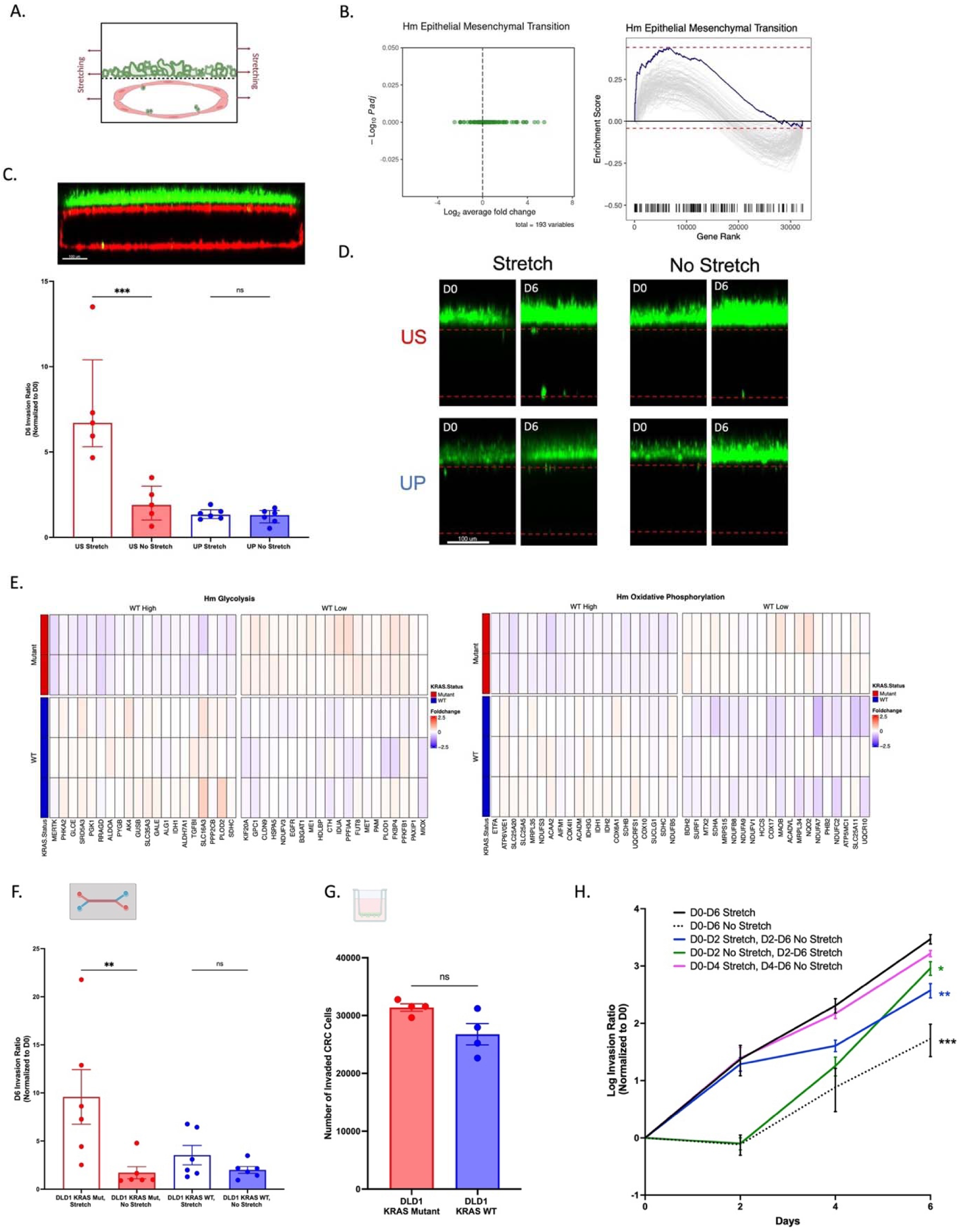
Peristalsis-like physical forces drive invasive capabilities in KRAS mutant tumor cells. **(A.)**Schematic showing the investigation of peristalsis-like mechanical forces on tumor cell behavior. Schematic was made using BioRender.**(B.)** Volcano plot (left) and ridgeplot (right) showing GSEA results of the Epithelial Mesenchymal Transition Hallmark gene set based on differential gene expression analysis of stretched vs not stretched conditions in the KRAS mutant setting. N=2 independent donors. **(C.)** Regions of the chips were imaged via confocal microscopy and a 3-D reconstruction was produced for quantifying the number of GFP+ tumor cells in the top channel and the number of GFP+ tumor cells that had invaded into the bottom, endothelial compartment, demarcated by RFP HIMEC cells (top image; scale bar represents 100μm). Invasion of US and UP in stretched and not stretched conditions was measured on day 0 and day 6 of the experiment. An invasion ratio was calculated based on the number of GFP+ cells in the bottom channel compared to the top channel and normalized by the day 0 counts. N=5-6 chips. Individual data are shown, with mean ± SEM represented and analyzed using a one-way ANOVA; ***p<0.001(.**D.)** Representative images show different invasion behavior for US and UP organoids, as quantified in C(.**E.)** Glycolysis and oxidative phosphorylation Hallmark gene sets were visualized using fold changes in patients with samples from stretched and not stretched conditions. N=5 independent doners; 2 KRAS mutant, 3 KRAS wildtype. After calculating foldchanges by patient, genes were ranked by p-value derived from a t-test between patients from different mutational status. For a given gene set, the 30 most significant genes from the gene set were selected for inclusion in the heatmap. Patients were ordered first by group and subsequently by average foldchange of the top 30 genes within each group. Foldchanges are thresholded to +/- 2.5 for visualization. **(F.)** Invasion of DLD1 KRAS Mutant and Wildtype tumor cells in stretched and not stretched on-chip conditions was measured on day 6 of the experiment. An invasion ratio was calculated based on the number of GFP+ cells in the bottom channel compared to the top channel and normalized by the day 0 counts. N=6 chips. Individual data are shown, with mean ± SEM represented and analyzed using a one-way ANOVA; **p<0.01. Schematic from BioRender.**(G.)** Invasive behavior of DLD1 KRAS Mutant and Wildtype tumor cells was measured 48 hr after cell seeding using traditional transwell invasion assays. N=4. Individual data are shown, with mean ± SEM represented and analyzed using a t-test. Schematic from BioRender(.**H.)** Stretching was introduced or removed from chips at various timepoints throughout the experiment. Invasion via fluorescence microscopy was measured every 2 days throughout the experiment and the number of GFP+ tumor cells was invaded and normalized by day 0 counts. Data is shown as mean ± SEM. N=4 chips. Invasion data on day 6 was compared between groups via one-way ANOVA; *p<0.05, **p<0.01, ***p<0.001.

Given that the patient-derived organoids used in this study spanned different KRAS statuses (N=3 KRAS wildtype (WT) and N=2 KRAS mutant (MUT)), we repeated the analysis of differentially expressed genes (DEG) and GSEA for stretched vs. not stretched conditions separately for KRAS MUT and KRAS WT organoid-on-chips. The EMT pathway was highly significant for the stretched vs. not stretched KRAS MUT organoid-on-chips (adjusted p-value = 9E-07, NES = 2.0), as shown in **Figure 3B**, yet it did not reach significance for the KRAS WT organoid-on-chips (adjusted p-value = 0.354, NES = 1.3).

Recognizing the strong association of EMT with metastasis^36^ and building upon our previous work employing an OOC model to study intravasation events in CRC^37^, we initiated an investigation into whether mechanical forces within our organoid-on-chips influenced tumor cell invasion. To facilitate this investigation, we fluorescently labeled the US (KRAS MUT) and UP (KRAS WT) organoids using Histone 2B-GFP (H2B-GFP) lentivirus prior to their seeding on the chips. Subsequently, we monitored GFP-tagged tumor cells via confocal imaging, and quantified invasion by assessing the number of GFP-positive cells that had penetrated from the upper epithelial channel into the lower endothelial channel, effectively mimicking intravasation (3D reconstruction seen in top portion of **Figure 3C**).

Under the stretched condition, US demonstrated an increase in tumor cell invasion, while UP did not (as depicted in **Figure 3C & 3D**). Importantly, the peristalsis-like mechanical strain did not disrupt the barrier function, as evidenced by a comparison of P_app_ values between stretched and static conditions (provided in **Table S5**), and further confirmed by immunofluorescence staining of ZO-1 and VE-Cadherin (**Figure S3**). Furthermore, no substantial variations in tumor cell growth due to peristalsis were observed between the stretched and unstretched conditions, as evidenced by the cell counts of US and UP organoids in the upper channel (**Figure S4**).

To better understand the connection between KRAS status and the metabolism shifts noted above, we analyzed stretched vs not stretched gene expression fold changes in KRAS MUT compared to KRAS WT organoid-on-chips. GSEA indicated that several metabolism pathways, such as oxphos, fatty acid metabolism, and glycolysis (adjusted p-value = 6.85E-6, NES = 1.99; adjusted p-value = 0.00017, NES = 1.86; adjusted p-value = 0.005, NES = 1.6, respectively), were enriched in the KRAS WT organoid-on-chips in the stretched condition (**Table S6**). To highlight the contrasting metabolic response to peristalsis, we plotted the fold changes of genes associated with oxphos and glycolysis in both KRAS WT and MUT organoid-on-chips under stretched and non-stretched conditions. Notably, we observed divergent responses to stretching. For example, genes within the glycolysis and oxphos gene set that exhibited higher fold changes in the stretched KRAS WT organoid-on-chips (labeled as “WT High” in **Figure 3E)**, demonstrated lower fold changes in the stretched KRAS MUT organoid-on-chips. The decreased oxphos seen in stretched KRAS MUT organoid-on-chips, compared to stretched KRAS WT organoid-on-chips, supports the change in glycolytic flux (i.e., “Warburg Effect”) observed in KRAS mutant tumors^38, 39^.

To further investigate the impact of KRAS mutational status on peristalsis-mediated invasion, we employed an isogenic CRC cell line consisting of DLD1 KRAS WT and DLD1 KRAS MUT cells^40^. Consistently, we observed that the DLD1 KRAS MUT cells exhibited increased invasion under stretched conditions, whereas the DLD1 KRAS WT cells did not respond in a similar manner (**Figure 3F**). To validate that this observed differential phenotype was indeed in response to mechanical forces, we conducted measurements of tumor cell invasion in a static environment using a transwell system. In this controlled setting, we did not detect any significant differences in invasive capabilities between the DLD1 KRAS WT and MUT cells (**Figure 3G**). Furthermore, we extended our investigation to other non-CRC cancer cell lines and observed a parallel trend. Specifically, KRAS MUT pancreatic cancer cells, PANC1, displayed increased invasion in response to mechanical stretch on-chip, whereas KRAS WT breast cancer cells, MCF7, did not exhibit a similar response (**Figure S5**).

To gain a more comprehensive understanding of the dynamics of the peristalsis-mediated invasion phenotype, we monitored the invasion of KRAS MUT HCT116 tumor cells at two-day intervals from day 0 to day 6. Throughout the experiment, we applied cycles of initiating and discontinuing rhythmic stretching. As depicted in **Figure 3H**, the tumor cells exhibited dynamic responses to the presence of peristalsis-mimicking stretching, evident in the increase in invasion rate (illustrated by the green line in **Figure 3H**) upon initiating stretch on day 2. Conversely, when we ceased the stretching stimulus on day 2 of the experiment, we observed a reduction in invasion, as shown by the blue line in **Figure 3H**. Furthermore, a significant difference in invasion on day 6 was observed between the chips subjected to continuous stretching for the entire six days compared to those where stretching commenced on day 2 (green line; p<0.05) and chips where stretching was terminated on day 2 (blue line; p<0.01).

### Organ-on-chip with physiological forces produces a GABAergic TME

To further investigate the alterations occurring in KRAS mutant tumor cells in response to peristalsis, we harnessed the capability to collect and analyze three distinct components from the chips: (1) tumor cells growing in the upper channel of the OOC, (2) tumor cells that had invaded into and adhered to the lower endothelial compartment, or (3) tumor cells in circulation that had been carried out of the channel and remained viable in the endothelial effluent (as schematically depicted in **Figure 4A**). Initially, we retrieved invaded circulating tumor cells from the endothelial effluent of both stretched and non-stretched HCT116-chips on day 6. Subsequently, we conducted RNAseq analysis and revealed shifts in GO BP pathways associated with cell migration, integrin binding, and cell-cell adhesion, all reinforcing the heightened invasiveness of these cells. Notably, the analysis also exposed significant changes in GABAergic-related pathways, an unexpected finding depicted in **Figure 4B**.

**Figure 4.**
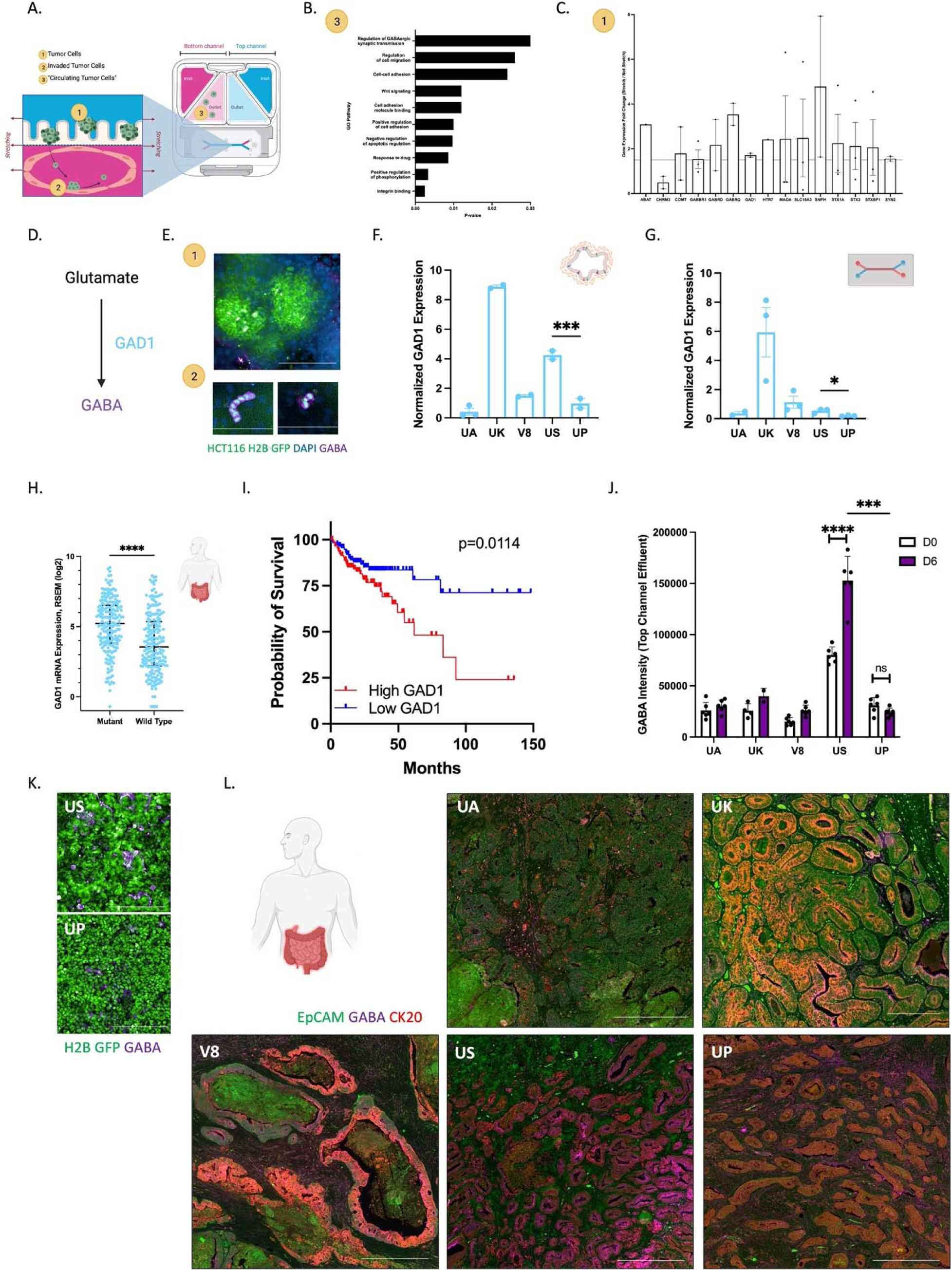
CRC-Chip with physiological forces produces a GABAergic TME. **(A.)** Schematic representing the initial events of the metastatic cascade that can be measured using the CRC-OOC. Tumor cells in the top channel (1) can be visualized and analyzed separately from tumor cells that have invaded and adhered in the endothelial compartment (2). Additionally, tumor cells that are found in the endothelial effluent (3; circulating tumor cells (“CTC-like” cells)) can also be collected and analyzed.**(B.)** Circulating tumor cells were collected from the endothelial effluent of stretched and not stretched HCT116 CRC-Chips and RNAseq was performed. GO Pathway analysis was performed on a subset of genes with either a 2-fold difference between stretched and not stretched CTCs or an FDR-adjusted p-value <0.1. N=2 biological replicates with 3 pooled chips in each replicate. **(C.)** Gene expression of neurotransmitter-related genes were measured by a neurotransmitter-specific PCR array. HCT116 tumor cells of stretched and not stretched chips were harvested on day 6 from the top epithelial channel and isolated via FACs. Data is displayed as the gene expression fold change of stretch versus not stretch conditions Expression was normalized to the average of 5 housekeeping genes. Genes that had a 1.5-fold increase or decrease in the stretched condition are displayed. N=3 biological replicates with 3 chips pooled per biological replicate. **(D.)** Schematic of the production of GABA from glutamate by the enzyme GAD1. In this figure, experiments related to GABA are indicated in purple and experiments related to GAD1 are indicated in light blue. **(E.)** Representative confocal immunofluorescent images of the epithelial (top; 1) or endothelial (bottom; 2) channel of the CRC-Chips stained for GABA (purple) on day 6. Invaded HCT116 H2B-GFP stain positive for GABA, while HCT116 H2B-GFP tumor cells that are in the top channel stain weakly for GABA. DAPI stains the nuclei of Caco2 C2BBe1 cells in the top channel and endothelial cells in the bottom channel. Scale bars represent 200 μm in the top channel image and 100 μm in the bottom channel images. Top channel images are maximum projections that span a 35 μm Z-height with a 5 μm step size. Bottom channel images are maximum projections that span a 10 μm Z-height with a 5 μm step size.**(F.)** RNAseq analysis was performed on CRC organoids and normalized GAD1 expression is shown. N=5 independent donors with 2-3 replicates each. Individual data points are shown and mean ± SEM is displayed. Analysis between US and UP data was performed using an unpaired t-test; ***p<0.001 (**G.)** CRC organoids were isolated from stretched chips and qPCR analysis of GAD1 gene expression was performed. N=5 independent doners with 3 replicates each. Individual data points are shown and mean ± SEM is displayed. Analysis between US and UP data was performed using an unpaired t-test p<0.05. (**H.)** GAD1 mRNA expression from TCGA in KRAS, NRAS, or BRAF mutant primary colon cancer tumors. N=196 patients with KRAS, NRAS, or BRAF mutant tumors; N=201 patients with KRAS, NRAS, or BRAF wildtype tumors. Individual data points are shown and median with interquartile range is represented. Data was analyzed with an unpaired t-test; ****p<0.0001. (**I.)** Kaplan-Meier curve with univariate analysis of the survival of patients with KRAS, NRAS, or BRAF mutated CRC tumors based on high versus low expression of GAD1 (defined as above or below the median GAD1 mRNA expression z-score of 0.3). Data was extracted from the TCGA. N=254 patients. Data was analyzed using a log-rank (Mantel-Cox test). (**J.)** Effluent from the epithelial channel of the patient-derived organoids was collected on day 0 (D0) and day 6 (D6). GABA intensity was analyzed from extracted metabolites N=6 chips per timepoint per patient; n=4 on D0 and 2 on D6 for UK. Data was analyzed using a two-way ANOVA; ***p<0.001; ****p<0.0001. (**K**.) US-H2B-GFP (top) and UP-H2B-GFP (bottom) stretched tumor-chips were stained for GABA (purple). Scale bars represent 200 μm. **L.** Representative 10x immunofluorescence images of the 5 tumors stained for EpCAM (green), CK20 (red), and GABA (purple). Scale bars represent 500 μm and 200 μm for UK. All schematics were made in or are from BioRender.

To better understand the changes associated with GABA signaling and increased invasion, we isolated GFP+ HCT116 tumor cells from the upper epithelial channel under both stretched and non-stretched conditions using flow cytometry. Then, we conducted a targeted neurotransmitter mRNA expression array to identify alterations in gene expression related to various neurotransmitter signaling pathways, including GABA. Our initial findings, as depicted in **Figure 4C**, revealed changes in genes including cholinergic (CHRM3, SLC18A3), catecholaminergic (COMPT, MAOA, HTR7), and general neurotransmitter vesicle release and uptake (SNPH, STX1A, STX3, STXBP1, SYN2). Furthermore, genes encoding subunits of the GABA receptors exhibited upregulation in response to mechanical forces. This is evident in the 1.5- fold increase in GABBR1 expression (a subunit of GABA R_B_) and the >2-fold increase in GABRD and GABRQ expression (two subunits of GABA R_A_). Additionally, GAD1, the enzyme responsible for converting glutamate into GABA (as schematically illustrated in**Figure 4D**), showed a >1.5-fold increase in expression in the stretched HCT116 tumor cells. Under stretched conditions, the invaded HCT116 tumor cells exhibited strong positive staining for GABA, as indicated in **Figure 4E**. These findings imply a potential link between GABA levels and the heightened invasive capabilities within the peristaltic CRC TME.

Building upon the presence of a neurotransmitter-rich TME within the organoid-on-chips (as shown in **Figure 2F**) and our observation of heightened GABAergic signaling in HCT116 tumor cells-on-chip when subjected to peristaltic forces, we investigated GAD1 expression within the organoid-on-chips. We observed variability in GAD1 gene expression among organoids alone, as illustrated in **Figure 4F**, as well as on-chip, as shown in **Figure 4G**. Interestingly, US and UP organoids exhibited distinct GAD1 expression patterns, with US organoids displaying significantly higher GAD1 expression compared to UP organoids, both in the organoid model (p<0.001) and on-chip (p<0.05). Given the previously described increased invasion in response to mechanical forces in the KRAS mutant setting, we sought to investigate a potential link between KRAS mutant tumors and GAD1. We conducted an analysis of GAD1 expression in CRC patient samples sourced from The Cancer Genome Atlas (TCGA) through the cBioPortal database. Through the examination of RNA-seq data, our findings reveal a noteworthy increase in GAD1 expression in primary colon tumors carrying KRAS, NRAS, or BRAF mutations, when compared to their KRAS wild-type counterparts (p<0.0001), shown in **Figure 4H**. Moreover, elevated GAD1 expression in KRAS, NRAS, or BRAF mutant colon tumors is indicative of a less favorable prognosis (p = 0.0114), as illustrated in **Figure 4I**. It is important to note that, to enhance the sample size of patient tumors for this study, we incorporated tumors irrespective of their treatment history.

To understand the connection between GAD1 levels and GABA secretion, we performed a targeted mass spectrometry-based metabolomics analysis of top channel effluent from stretched organoid-on-chips on day 0 and day 6. We observed variable, but measurable amounts of GABA in the effluent with organoids derived from different donors, as demonstrated in **Figure 4J**. Throughout the six-day experiment, we observed an increase in GABA levels in the US effluent (p<0.0001), while UP showed no significant change in GABA levels over time (n>0.05) (**Figure 4J**). Furthermore, we evaluated GABA levels in US and UP organoid-on-chips through immunostaining, confirming higher GABA levels in the US organoid-on-chips compared to the UP organoid-on-chips, consistent with the effluent data (**Figure 4K**). We expanded our analysis to the corresponding patient tumor tissues, staining for GABA, as well as CK20 and EpCAM to delineate CRC regions. As depicted in **Figures 4L** and **Figure S6**, the US tumor exhibited the highest levels of GABA within the cancer tissue, evident from the overlap (maroon) of CK20 (red) with GABA (purple). In contrast, UA, UK, V8, and UP tumor tissue all displayed lower levels of GABA, aligning with the organoid-on-chip measurements. These results indicated that the peristaltic organoid-on-chips recapitulated GABAergic aspects of patient tumors.

### Inhibiting GABA catabolism reduces peristalsis-mediated invasion in KRAS mutant tumor cells

To gain deeper insights into the influence of GABA on tumor cell invasion, we introduced exogenous GABA into the top epithelial channel of HCT116 tumor-chips during the 6-day experiment. The presence of exogenous GABA led to an increase in tumor cell invasion by HCT116 cells, as illustrated in **Figure 5A**. This observation suggests that these tumor cells are responsive to a GABAergic microenvironment. Our mass spectrometry analysis revealed low levels of endogenous GABA in the effluent from the epithelial channel of HCT116 tumor-chips. This characteristic positions HCT116 tumor-chips as a particularly suitable model for investigating the effects of exogenously introduced GABA. To examine the uptake of GABA by the tumor cells, we introduced ^13^C_4_-labeled GABA into the upper channel of the HCT116 tumor-chips. Subsequently, we collected tumor cells from this top epithelial channel and quantified the intracellular levels of ^13^C_4_-labeled GABA using mass spectrometry. The presence of intracellular^13^C_4_- labeled GABA, as depicted in **Figure 5B**, indicates that in the presence of a GABAergic environment, these tumor cells actively uptake GABA.

**Figure 5.**
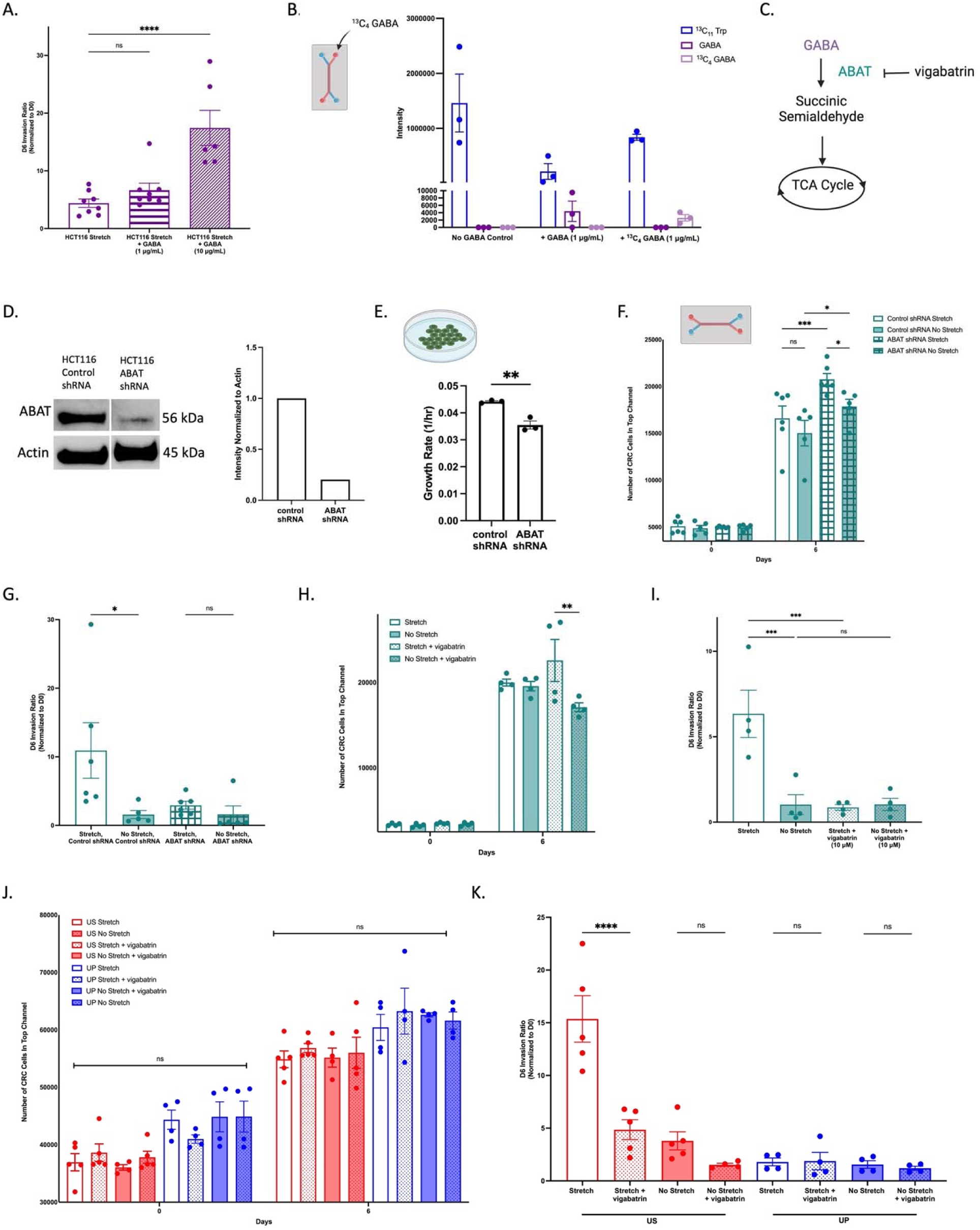
Inhibiting GABA catabolism reduces peristalsis-mediated invasion. **(A.)** Invasion of HCT116 tumor-chips in the presence or absence of exogenous GABA (flowed through the epithelial channel) was measured on day 6 (D6) of the experiment and normalized to day 0 (D0) invasion. N=6 chips. Individual data are shown, with mean ± SEM represented and analyzed using a one-way ANOVA; ****p<0.0001. **(B.)** Intracellular [^13^C_4_]GABA or unlabeled GABA was measured via mass spectrometry-based metabolomics in the HCT116 tumor-chips after the addition of exogenous GABA for six days. N=3 chips. **(C.)** Schematic of GABA catabolism by ABAT, subsequent entry into the TCA cycle, and inhibition of ABAT activity by vigabatrin. In this figure, experiments related to GABA are indicated as purple, and experiments related to ABAT are indicated as teal.**(D.)** Western blot analysis of ABAT in shRNA control or ABAT shRNA HCT116 tumor cells. Cropped western blot (left) and quantification (right) confirm knockdown of ABAT. **(E.)** Growth rate of ABAT-knockdown or control HCT116 tumor cells when grown in traditional cell culture methods. N=3. Individual data are shown and mean ± SEM are represented. Data was analyzed using a t-test; **p<0.01. **(F.)** Numbers of ABAT-knockdown or control HCT116 tumor cells in the top channel on-chip as measured via fluorescence microscopy and quantified on day 0 (D0) and day 6 (D6). N=5-6 chips. Individual data are shown and mean ± SEM are represented. Data was analyzed using a two-way ANOVA; *p<0.05;***p<0.001.**(G.)** Invasion of ABAT knockdown (KD) or control shRNA HCT116 tumor-chips in the presence or absence of stretching was measured on day 6 (D6) of the experiment and normalized to day 0 (D0) invasion. N=5-6 chips. Individual data points are shown and mean ± SEM are represented. Data was analyzed using a one-way ANOVA; *p<0.05. **(H.)** Numbers of HCT116 tumor cells in the top channel on-chip in the presence or absence of stretching, with or without vigabatrin was measured via fluorescence microscopy and quantified on day 0 (D0) and day 6 (Day 6). Individual data are shown and mean ± SEM are represented. N=4 chips. Data was analyzed using a two-way ANOVA; **p<0.01. **(I.)** Invasion of HCT116 tumor-chips in the presence or absence of stretching, with or without vigabatrin was measured on day 6 (D6) of the experiment and normalized to day 0 (D0) invasion. N=4 chips. Individual data points are shown and mean ± SEM are represented. Data was analyzed using a one-way ANOVA; ***p<0.001.**(J.)** Numbers of US-H2B-GFP (red) or UP-H2B-GFP (blue) tumor cells in the top channel on-chip in the presence or absence of stretching, with or without vigabatrin was measured via fluorescence microscopy and quantified on day 0 (D0) and day 6 (Day 6). N=4-5 chips. Individual data are shown and mean ± SEM are represented. Data was analyzed using a two-way ANOVA; ns=p>0.05.**(K.)** Invasion of US-H2B-GFP or UP-H2B-GFP organoid-tumor chips in the presence or absence of stretching, with or without vigabatrin was measured on day 6 (D6) of the experiment and normalized to day 0 (D0) invasion. N=4-5 chips. Individual data are shown and mean ± SEM are represented. Data was analyzed using a one-way ANOVA; **p<0.01. All schematics were made in or are from BioRender.

Given the heightened expression of 4-aminobutyrate aminotransferase (ABAT), the enzyme responsible for catabolism of GABA, observed in the stretched HCT116-tumor-chips (**Figure 4C**), the metabolic alterations induced by the peristaltic environment in the patient-derived organoid-on-chips (discussed previously and shown in **Figure 3**,**Table S4** and **S6**), and the known ability of tumor cells to utilize the GABA shunt as an energy source ^41, 42^, our hypothesis centers on the potential role of GABA metabolism in influencing the invasive phenotype of these tumor cells. Considering this, we employed genetic knockdown techniques to target ABAT in HCT116 cells (schematically shown in **Figure 5C**; **Figure 5D** and **Figure S7**). We subsequently conducted an examination of tumor cell invasion under stretching conditions, directly comparing the behavior of control cells with ABAT knockdown (ABAT KD) HCT116s. While ABAT KD HCT116 tumor cells exhibited slower growth in conventional cell culture methods compared to control HCT116 tumor cells (**Figure 5E**), a shift occurred on-chip, where more ABAT KD HCT116 tumor cells were observed than control HCT116 tumor cells by day 6 (**Figure 5F**). The ABAT KD HCT116 tumor cells exhibited no discernible increase in invasion when exposed to stretching, in contrast to the HCT116 shRNA control cells, as illustrated in **Figure 5G**.

Additionally, we applied a pharmacological approach to inhibit ABAT by introducing vigabatrin, a known ABAT inhibitor, into the upper epithelial channel throughout the duration of the chip experiments. The inhibition of ABAT resulted in a decrease in tumor cell numbers in the top channel of vigabatrin-treated, non-stretched chips when compared to the vigabatrin-treated, stretched chips (**Figure 5H**). Similar to the observations in ABAT KD cells, the pharmacological inhibition of ABAT led to a reduction in invasion in HCT116 tumor chips in response to peristalsis, as depicted in **Figure 5I**. In the organoid-on-chip models, there were no discernible differences in growth in response to ABAT inhibition in either the US or UP organoids, as shown in **Figure 5J**. However, inhibiting ABAT resulted in reduced invasion under stretched conditions in the US organoid-on-chip, while the invasion of UP organoids was unaffected by vigabatrin treatment, as demonstrated in **Figure 5K**. These findings support the notion of KRAS activating mutations playing a role in this GABAergic, mechano-sensitive invasion phenotype.

## DISCUSSION

Our main goal was to create a tumor organoid-on-chip model capable of maintaining patient diversity and replicating crucial mechanical properties pertinent to CRC, including fluid dynamics and peristalsis-like movements. This goal becomes especially significant in light of emerging evidence suggesting that mechanical forces may contribute to CRC progression^43, 44^. To achieve this, we seeded patient-derived tumor organoids and endothelial cells within an OOC system, to establish an epithelial – endothelial tissue-tissue interface and intact barrier. This integration allows for the study of metastasis in a personalized manner, which is a challenging aspect to replicate in a laboratory setting. Microfluidic OOC models, like the one we developed, serve as valuable tools for investigating various stages of the metastatic process, facilitating a better understanding of this complex phenomenon^45–47^.

OOC models have traditionally been employed in drug development, pharmacokinetics, and toxicology. More recently, efforts have been directed towards the convergence of organoids and OOC models to create organoids-on-chips, aiming to achieve higher physiological relevance^48^. While several organoid-on-chip models have been developed to mimic normal tissue structures and functions^29, 49, 50^, there is still a shortage of such models in cancer research^51^. In our study, we observed that the organoid-on-chip model exhibited a closer transcriptional resemblance to tumor tissue when compared to organoids cultured alone. This alignment can likely be attributed to the incorporation of physical forces and vasculature components within the OOC system. It’s important to acknowledge that achieving a complete replication of *in vivo* tumors when working with 3D *in vitro* models is extremely challenging. Instead, the primary objective is to capture specific essential features relevant to their intended context of use. Additionally, it is beneficial to identify potential gaps in a model. To achieve this, we conducted an over-enrichment analysis, shedding light on the elements that were noticeably absent in the organoid-on-chips when compared to the actual tumor tissue. As expected, we observed a lack of immune cell signature and certain ECM components. Nonetheless, it is worth noting that our model is adaptable, allowing for possible modifications such as the inclusion of immune cells or the incorporation of patient-specific tumor ECM in future research.

Recognizing the significance of the TME in cancer progression and treatment response, it’s clear that the impact of mechanical forces remains a largely understudied area. While *in vivo* animal studies do investigate these TME forces, there’s a gap in our understanding when it comes to their relevance in the human context and our ability to manipulate them. Other *in vitro* model systems like spheroids and organoids do capture significant aspects of the human TME, including factors like oxygen and nutrient gradients and the inclusion of stromal cell types. However, they may not fully account for the influence of physical forces and tissue-to-tissue interfaces, which also contribute to cancer development and metastasis. In particular, organoid models often overlook the role of luminal flow in the gut, potentially leading to the accumulation of metabolic waste within cellular environments. The combination of fluid flow and cyclic stretching has shown promise in extending cellular lifespan, supporting cell polarization, and promoting multi-lineage differentiation, particularly in models of the intestine^54, 55^. This emphasizes the critical need for developing microfluidic OOC models capable of accommodating these essential physical forces.

The influence of biophysical forces on cancer cell behavior has been a focus in research using OOC models, particularly in the context of breast and ovarian cancer. For instance, compressive stimuli applied to ovarian cancer cells via a 3D bioreactor has been associated with increased cell proliferation, invasion, and chemoresistance^56^. Similarly, mechanical compression in breast cancer has been linked to increased invasion, primarily by stimulating leader cells that coordinate migration^57^. Furthermore, breast cancer tumoroids cultured in a 3D environment under fluid flow and pressure have shown increased aggressiveness and reduced sensitivity to the drug doxorubicin^58^.

In our case, exploring the mechanical aspects of CRC cell behavior within the context of gut physiology is an area of significant interest. A handful of studies, including our previous work using OOC technology^37^, have delved into how peristalsis-like motions impact the invasive potential of CRC cell lines^59, 60^. This aspect is challenging to investigate using alternative models, and numerous questions remain unanswered. Of particular interest is whether these mechanical forces uniformly affect all cancer cells or if specific molecular alterations render some cells more vulnerable. In our current study, we observed significant transcriptomic alterations related to EMT and increased intravasation potential of KRAS MUT tumor cells within the organoid-on-chips subjected to peristalsis-like motion compared to WT. It is important to note that these differences between KRAS MUT and WT cells were not observed in static transwell conditions, emphasizing the role of mechanical forces. Previous research has suggested that Ras-transformed cells may display increased sensitivity to mechanical forces^61^. This phenomenon was demonstrated in a Drosophila midgut model, where Ras-transformed cells showed enhanced dissemination. This invasive behavior appeared to be regulated, in part, by the Piezo mechanosensitive ion channel^62^. Another study proposed that Ras-transformed epithelial cells displayed enhanced apical extrusion capabilities, which allowed them to avoid ferroptosis. This was attributed to the accumulation of CD44 and COL17A1 and metabolic alterations^63^. Similarly, we observed metabolic differences between KRAS MUT and WT cells in response to mechanical stimuli within the organoid-on-chips. It’s worth noting that KRAS mutant CRC tumors are known to exhibit metabolic dysregulation, with a higher reliance on glycolysis^40, 64^. Although we did not conduct a complete metabolic analysis in this study, our preliminary ‘omics’ studies suggest that mechanical stimuli may play a role in regulating the metabolic shift seen in KRAS mutant CRC tumors.

Furthermore, in pathological conditions such as CRC, there is often dysregulation of neurotransmitters, which can impact intestinal function and homeostasis^26^. Over the course of CRC progression, both cancer cells and neurons within the enteric nervous system of the intestine release neurotransmitters that have the potential to influence signaling pathways and promote the survival of cancer cells^65^. Additionally, migration of CRC cancer cells along enteric neurons has been observed^66^. Although our system does not incorporate neurons responsible for peristalsis in the gut *in vivo*^67^, we have the capacity to regulate peristalsis within the OOC system. In our organoid-on-chip microenvironment, we detected variations in neurotransmitter levels among different patients. Specifically, we identified GABA as a neurotransmitter taken up and synthesized by CRC tumor cells, and its enrichment was particularly notable in KRAS mutant tumors. Past studies have demonstrated that elevated GABA in clinical samples is associated with poor clinical outcomes and tumor cell-derived GABA contributes to cell proliferation and immunosuppression^68^. Additionally, upon analyzing TCGA samples, we observed a correlation between RAS mutational status and increased expression of GAD1, the enzyme responsible for GABA synthesis, with high GAD1 levels being associated with poorer survival.

In situations of metabolic stress, GABA can function as an energy source through a metabolic pathway known as the "GABA shunt." In a study focused on breast cancer metastases to the brain, GABAergic characteristics were observed^41^. The enzyme responsible for GABA catabolism, ABAT, was found to be necessary for medulloblastoma cells to disseminate and establish metastases^69^. Similarly, when we hindered CRC tumor cells’ capability to metabolize GABA by genetically and pharmacologically inhibiting ABAT, we observed a significant decrease in tumor cell invasion in the presence of mechanical forces. In another study investigating the relationship between a GABAergic TME and melanoma using a zebrafish model, it was discovered that GABAergic communication, which includes GABA R_A_ signaling and electrochemical crosstalk, between melanoma cells and keratinocytes, promoted melanoma growth and initiation^70^. Further investigation is needed to fully uncover the intricate mechanisms driving cancer metastasis and to understand the impact of neurotransmitter signaling within the TME.

In summary, our study has provided valuable insights into the influence of mechanical forces and neurotransmitter signaling on CRC cell behavior using an organoid-on-chip model. By closely mimicking the TME and incorporating patient-specific elements, we have revealed interesting patterns in KRAS mutant and wild-type cell responses to peristalsis-like motion. Additionally, our identification of GABAergic properties and the role of the GABA catabolism enzyme ABAT in cancer cell invasion suggest potential implications for CRC progression and targets for drug development^33^. These findings enhance our understanding of cancer biology and underscore the versatility of the organoid-on-chip system as a tool for exploring complex and heterogeneous interactions in cancer research, offering potential avenues for further study and personalized treatment approaches.

## LIMITATIONS OF THE STUDY

CRC is a complex and diverse disease. Although we have developed organoids-on-chips from samples of five different patients with varying tumor characteristics, a more comprehensive analysis is required before making confident assertions regarding subtype-specific discoveries. Furthermore, the peristalsis values selected for these studies were based on clinical measurements of normal, healthy intestine and colon tissue. It’s worth acknowledging that these values may not precisely replicate the *in vivo* conditions observed in colon cancer cases. Additionally, cancer and its treatments often lead to alterations in bowel functionality, a factor we are not replicating in this study.. Further research is essential to comprehend how tumor cells respond to a dynamically changing and potentially patient-specific physical TME.

## ACKNOWLEDGEMENTS

This research was supported by the NCI Tissue Engineering Consortium R01 CA241137 grant (SMM), a Stop Cancer Grant (SMM), and the USC Norris Comprehensive Cancer Center Core Grant P30CA014089 (HJL, SMM, USC Norris Cores). We would like to express our deepest gratitude to our philanthropic supporters, particularly to the Stephenson family, Emmet, Toni, and Tessa, for their donation of the Operetta/CLS HCS platforms and the funding support as part of the Stephenson Family Personalized Medicine Center. We are extremely grateful for the expertise, guidance, and support from Emulate, Inc. We would also like to thank: K. Kani for review and helpful comments on the manuscript; E. Spiller for discussions about organoid-on-chip experiments; S. Chilakala for developing and optimizing the mass spectrometry workflow; B. Larson for offering gastrointestinal pathology expertise; Y. Zhou for technical support with immunofluorescence; D. Hixon and B. Choi for western blot assistance; the University of Southern California Translational Pathology Core and the Molecular Genomics Core for providing tumor tissue slides and sequencing services; the Ellison Institute Bioanalytics Team (particularly N. Ung) for statistical support; the Ellison Institute Cell Line Team for assistance with organoid development and generation of fluorescently-labeled cell lines; J. Neman, Y. DeClerck, F. Battaglin, and S. Soni for meaningful discussions and guidance; and O. Castellanos and B. Tran for patient data assistance.

## AUTHOR CONTRIBUTIONS

C. Strelez designed, performed, analyzed, and interpreted the experiments and wrote the manuscript. RP performed, analyzed, and interpreted OOC experiments and helped write the manuscript. JSC performed, analyzed, and interpreted mass spectrometry experiments. CC analyzed RNAseq datasets and provided key interpretations of the data. AYY performed and analyzed mass spectrometry experiments and compiled data for figures in the manuscript. BH performed the transwell experiments, provided expertise in tissue staining, performed the imaging, and compiled images for the manuscript. C.Shah helped write the manuscript, performed OOC experiments, collected effluent, and performed flow cytometry sorting. KG performed optimization experiments for the organoid OOC and performed neurotransmitter array qPCR experiments, RXS performed CTC RNA seq analysis and key interpretations of the data. HJ aided in OOC experiments, data analysis, literature searches and writing of the manuscript. RL aided in performing cell culture experiments, particularly fluorescently labeling cell lines, and provided tissue staining assistance. AS performed qPCR experiments. HJL provided clinical expertise and patient samples. JEK provided conceptualization, experimental design support and interpretation of mass spectrometry data. SMM was responsible for study concept and design, interpretation of data, supervision of the study, and editing and revision of the manuscript. There was no outside writing assistance. All authors had access to the study data and reviewed, edited, and approved the final manuscript.

## DECLARATION OF INTEREST

The authors declare no competing interest.

### KEY RESOURCES TABLE

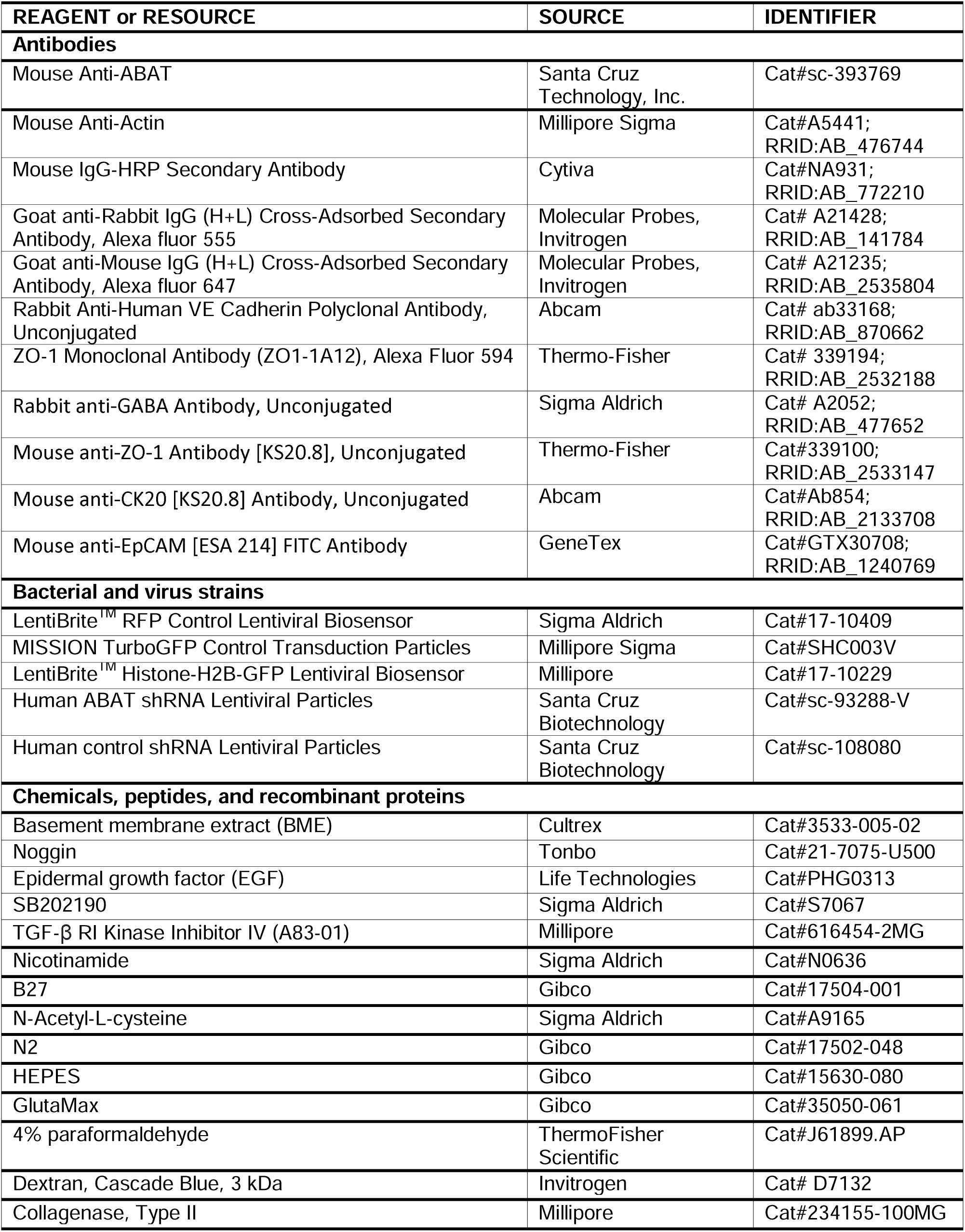
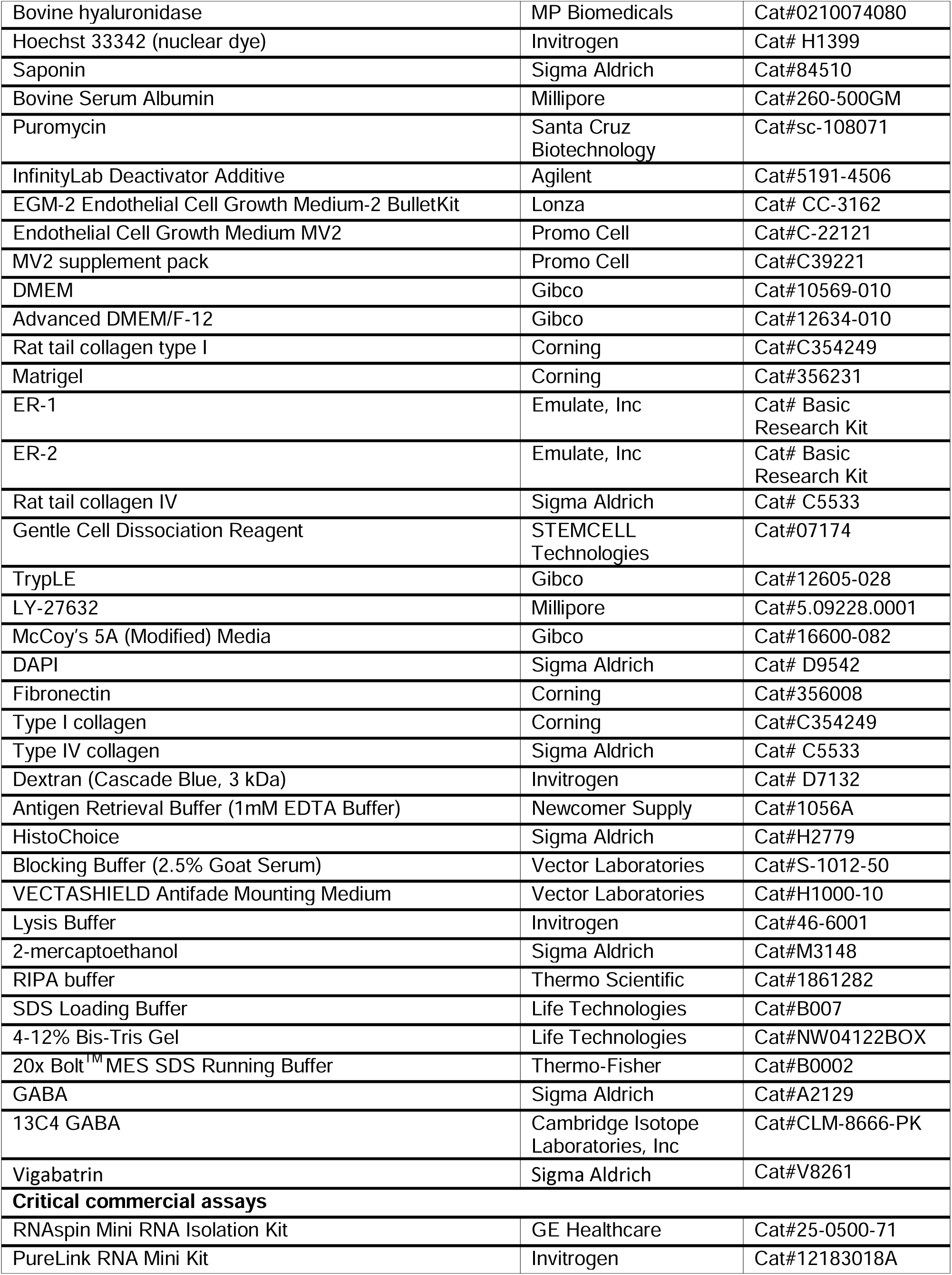
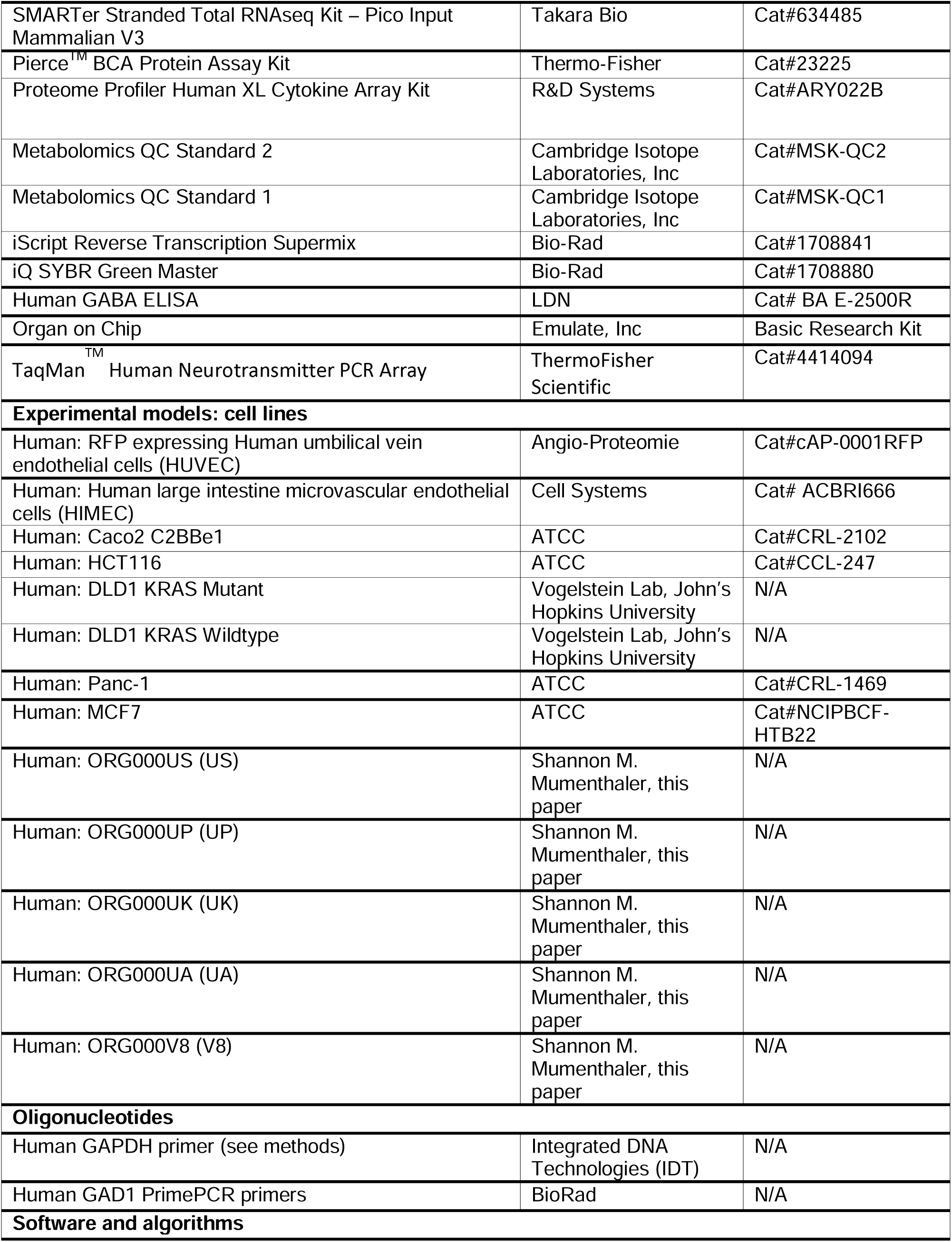
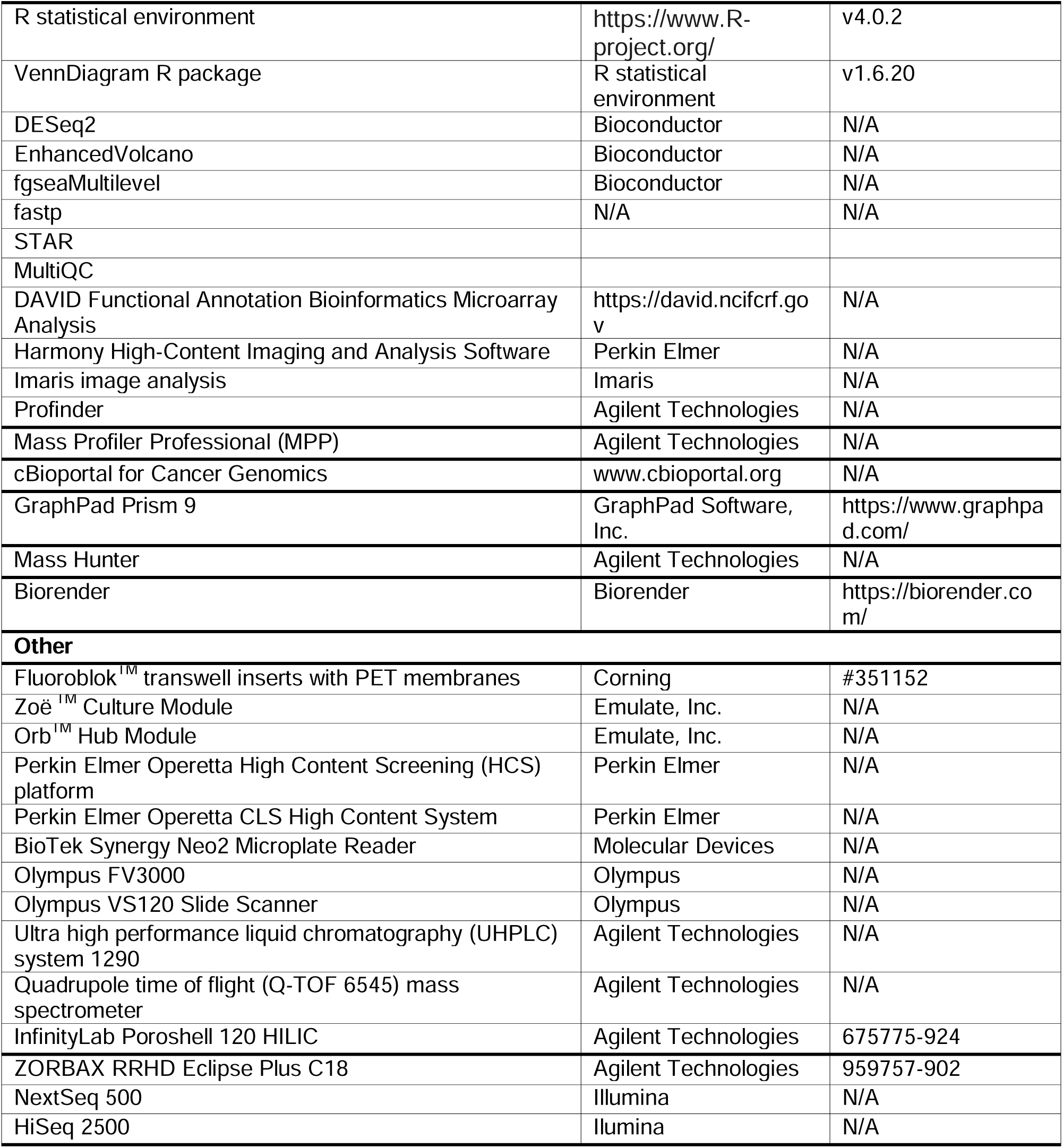

## METHODS

### Cell Lines

#### Commercially Available Cell Lines

Human umbilical vein endothelial cells (HUVEC) expressing Red Fluorescent Protein (RFP) (Angio-Proteomie, #cAP-0001RFP) were expanded in EBM-2 media with EGM-2 SingleQuots Supplements (2% FBS, 1% Penicillin-Streptomycin (Pen-Strep), Hydrocortisone, hFGF-B, VEGF, R3-IGF-1, Ascorbic Acid, hEGF, and heparin in proprietary concentrations) (Lonza #CC-3162; supplemented with 1% Pen-Strep in lieu of Gentamicin). Human large intestine microvascular endothelial cells (HIMEC) (Cell Systems, #ACBRI 666) were labeled with LentiBrite^TM^-RFP (Sigma, #17- 10409) and grown in Endothelial Cell Growth Medium MV2 (Promo Cell, #C-22121), with MV2 supplement pack (PromoCell, #C39221; supplemented with heat inactivated 5% FBS (Gemini, #100-500) in lieu of FCS) containing 5 ng/mL recombinant human epidermal growth factor (rhEGF), 10 ng/mL recombinant human basic fibroblast growth factor (rhbFGF), 20 ng/mL long R3 insulin-like growth factor (R3-IGF), 0.5 ng/mL recombinant human vascular endothelial growth factor 165 (rhVEGF), 1 μg/mL ascorbic acid, 0.2 μg/mL hydrocortisone and 50 μg mL-1 Primocin (InvivoGen #ant-pm-1). HIMEC cells used for chip conditions were between passage 6 and 8. Caco2 C2BBe1 cells (ATCC, #CRL-2102) were grown in DMEM (Gibco, #10569-010) with 10% fetal bovine serum (FBS) and 1% Pen-Strep. HCT116 (ATCC #CCL-247) were grown in McCoy’s 5A media (Gibco, #16600-082) with 10% FBS and 1% Pen-Strep, labeled with LentiBrite Histone-H2B-GFP Lentiviral Biosensor (Millipore, #17-10229), and sorted to achieve a pure fluorescent population. ABAT human shRNA lentiviral particles (Santa Cruz Biotechnology, Inc., #sc-93288-V) and control shRNA lentiviral particles (Santa Cruz Biotechnology, Inc., #sc-108080) were used to produce HCT116 H2B-GFP ABAT knock down cells at a multiplicity of infection (MOI) of 5 and 2 mg/mL of puromycin (Santa Cruz Biotechnology, #sc-108071) was used to select for knocked down cells. DLD1 cells were obtained from the Vogelstein Lab at John’s Hopkins University, labeled with MISSION TurboGFP Control Transduction particles (Millipore Sigma, #SHC003V), and 2 mg/mL of puromycin (Santa Cruz Biotechnology, #sc-108071) was used to select for knocked down cells. Cells were maintained in McCoy’s 5A media (Gibco, #16600-082) with 10% FBS and 1% Pen-Strep supplemented with 1 mg/mL of puromycin. Panc-1 cells (ATCC, #CRL-1469) were grown in DMEM (Gibco, #10569-010) with 10% FBS and 1% Pen-Strep, labeled with LentiBrite Histone-H2B-GFP Lentiviral Biosensor (Millipore, #17-10229), and sorted to achieve a pure fluorescent population. MCF7 cells (ATCC, #NCIPBCF-HTB22) were grown in DMEM (Gibco, #10569-010) with 10% FBS and 1% Pen-Strep, labeled with MISSION TurboGFP Control Transduction particles (Millipore Sigma, #SHC003V), GFP+ populations were selected by 2 mg/mL of puromycin and maintained in 1 mg/mL of puromycin. All cells were cultured under standard laboratory conditions (5% CO_2_, 37° C).

#### Patient-Derived Samples

Tissue resections were received from the USC Norris Comprehensive Cancer Center following Institutional Review Board (IRB) approval (Protocol HS-06-00678; approval date 08-02- 2019) and patient consent. Tumor profiles, including known tumor mutations, sex, and treatment information, are detailed in **Figure 1B**. Human organoids were derived from CRC tumors via a previously described method (Sato et al., 2011; Sato et al., 2009). Briefly, tumor pieces were minced and enzymatically digested using 1.5 mg/mL collagenase (Millipore, #234155), 10 μM LY27632 (Millipore, #5.09228.0001) and 20 μg/mL hyaluronidase (MP Biomedicals, #0210074080) for 30 minutes at 37°C. Established organoid cell lines were expanded by plating organoids with basement membrane extract (BME; Cultrex, #3533-005-02) cultured in colon media (Advance DMEM F12 (Gibco, #12634-010), supplemented with 10% FBS (Gemini, 100-500), 1% Pen-Strep (Gemini 400-109), 100 ng/mL Noggin (Tonbo, #21-7075-U500), 50 ng/mL epidermal growth factor (EGF) (Life Technologies, #PHG0313), 10 mM SB202190 (Sigma Aldrich, #S7067), 500 nM TGF-b RI Kinase Inhibitor IV (A83-01) (Millipore, #616454-2MG), 10 mM Nicotinamide (Sigma Aldrich, #N0636), 1 x B27 (Gibco, #17504-001), 1mM N-acetylcysteine (Sigma Aldrich, #A9165), 1 x N2 (Gibco, #17502-048), 1 x HEPES (Gibco, #15630-080), 1 x GlutaMax (Gibco, #35050-061) at 5% CO_2_, 37°C. Media was replaced every 2-3 days. CRC organoids used for chip conditions were between passage 12 and 23. After establishment and expansion of organoids, ORG000US and ORG000UP (US and UP) were subsequently labeled with H2B-GFP lentivirus as previously described (Kim et al. 2020). Briefly, organoids were dissociated with 50% TrypLE (Gibco; 12605-028) supplemented with 10 μM LY-27632 (Millipore, #5.09228.0001) and incubated at 37 °C for 5 min. After spinning, the pellet was resuspended in colon organoid media containing 5 µg mL-1 polybrene (Sigma; TR-1003-G) with LentiBrite Lentivirus H2B-GFP (MilliporeSigma; 1710229, 40 multiplicity of infection [MOI]) at 37 °C for 60 min. The pellet was then resuspended in BME and cultured and expanded. GFP-positive organoids were sorted with FACS using the ARIA IIu (BD Biosciences, San Diego, CA) to isolate only GFP positive-labeled cells.

### Colorectal Cancer Organ-on-Chip

Chips were acquired from Emulate, Inc and fabrication methods have been described previously by Emulate, Inc. In brief, the chips are made of transparent elastomeric polymer (polydimethylsiloxane, PDMS). They are divided into upper (1 mm high x 1 mm wide) and lower (0.2 mm high x 1 mm wide) microfluidic compartments separated by a thin, porous membrane (50 µm thick with 7 µm diameter pores; 17.1 mm^2^ co-culture region), with cancer cells in the upper compartment and endothelial cells in the lower compartment. Each compartment is coated with a tissue-specific ECM prior to cell seeding. The chips are attached to a Pod^TM^ portable module that encloses the chips to control sterility, holds inlet cell culture media and effluent, allows for monitoring via microscopy, and is designed to ensure no pressure differentials between channels. The chips and pod are then housed in an automated culture module instrument (Zoë^TM^ culture module and Orb^TM^ hub module, Emulate, Inc.) that controls the fluid flow and stretching forces while inside an incubator.

#### Caco2 + CRC Cell Lines

A CRC on-chip using commonly available CRC tumor cell lines has been previously described^37, 71^. Briefly, Emulate Chip-S1 Stretchable chips were activated by Emulate Reagent 1 and 2 (Emulate, Inc., ER-1 and ER-2) under UV light for 20 min at room temp. The epithelial and endothelial channels were coated with a mixture of 30 μg/mL type I collagen (Corning, #354249) and 100 μg/mL Matrigel (Corning, #356231) overnight at 4°C before washing with PBS. HUVEC cells (RFP-labeled or unlabeled) were seeded into the bottom channel (1-1.2×10^5^ cells in 20 μL; 5.8-7×10^5^ cells/cm^2^). The chips were inverted and incubated at 37°C for 2 hr. After HUVEC attachment, Caco2 C2BBe1 cells were seeded into the top channel (50,000-62,500 cells in 50 μL; 3-3.7×10^5^ cells/cm^2^), incubated for 2 hr at 37°C, and connected to flow. The chips were perfused with DMEM, 10% FBS, 1% Pen-Strep in the top channel and endothelial media (EBM-2 fully supplemented with 2% FBS, 1% Pen-Strep, Hydrocortisone, hFGF-B, VEGF, R3-IGF-1, Ascorbic Acid, hEGF, and heparin in proprietary concentrations) in the bottom channel at 30 μL/hr (0.02 dyne/cm^2^) starting the day after cell seeding. Cyclic, peristalsis-like membrane deformations (10% strain, 0.2 Hz) were also initiated the day after cell seeding using an electronic vacuum pump system (Emulate, Inc).

#### CRC-organoid-on-chip

Emulate Chip-S1 Stretchable Chips were activated by Emulate Reagents 1 and 2 (Emulate, Inc., ER-1 and ER-2) under UV light for 20 min at room temperature. The epithelial channel was coated with 250 μg/mL Matrigel (Corning, #356231) and the endothelial channel was coated with a mixture of 30 μg/mL Fibronectin (Corning #356008) and 200 μg/mL type IV collagen (Sigma #C5533) and incubated at 4°C overnight. The chips were then warmed at 37°C for 1 hr and each channel was rinsed with PBS and corresponding media. HIMEC (RFP-labeled or unlabeled) were resuspended at 6.0 x 10^6^ cells/mL and seeded on the bottom channel. The chips were incubated at 37°C, inverted for 2 hr, and returned to normal orientation to allow HIMEC cell adhesion throughout the channel membrane. CRC tumor organoids were collected from BME by incubating in Gentle Cell Dissociation Reagent (STEMCELL Technologies, #07174) at 4°C for 45-60 min, followed by dissociation with 50% TrypLE (Gibco; 12605-028) supplemented with 10 μM LY-27632 (Millipore, #5.09228.0001) and passed through a 40 μL cell strainer. Organoids were resuspended in colon media and seeded on the top channel at 11-14.3 x 10^6^ cells/mL density. All chips were incubated overnight in static conditions at 37°C, 5% CO_2_. The chips were then washed with fresh media and connected to Emulate portable pod modules filled with gas equilibrated medium and continuous flow (30μL/hr) was initiated through top and bottom channels for 48 hr. Cyclic conditions (10% strain, 0.2 Hz) were initiated after the formation of a monolayer, regarded as Day 0 and continued until end of experiment at Day 6.

### Barrier Function

To assess the barrier formation of each CRC organoid, 50 μg/mL Dextran (Cascade Blue, 3 kDa) (Invitrogen, #D7132) was added to the top channel, beginning on connection to flow (day -2). The Dextran was replenished at every media change and the effluent from both channels was collected throughout the experiment (Days -2, -1, 0, 1, 2, 4, 6). The fluorescence from the bottom channel was measured using a plate reader (Biotek) and apparent permeability (P_app_) was calculated using the following formula:

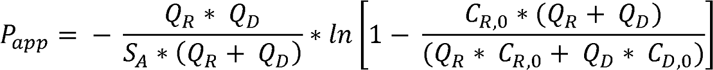

where S_A_ is the surface area of the culture overlapping channel (0.17cm^2^), Q_D_ & Q_R_ are the flow rates in the dosing channels (top epithelial) and receiving channel (bottom endothelial) respectively, in units of cm^3^/s, and C_R,0_ & C_D,0_ are the recovered concentrations in the receiving and dosing channels, respectively. For each donor, two independent experiments were conducted with duplicate or triplicate chips per condition.

### Immunofluorescence

#### Organoids

CRC organoids were seeded in 96 well plate with BME and cultured with colon media for 72 hr. Each well was rinsed with PBS and fixed with 4% Paraformaldehyde (PFA) (ThermoFisher Scientific, #J61899.AP) at 4°C on a rocker for 30 min. Each well was rinsed with 0.75% Glycine three times with 10 min incubation at room temperature followed by blocking buffer (5% bovine serum albumin (BSA), 0.2% Triton X-100, 0.04% Tween 20) for 2 hr at room temperature. Organoids were then stained overnight with primary antibody mouse anti-ZO-1 (1:250; Invitrogen #339100) in blocking buffer at 4°C. The next day, each well was rinsed with PBS three times with 20 min incubation at room temperature and stained with secondary antibody goat anti-mouse Alexa Fluor 647 (1:250; Invitrogen #A-21235) at room temperature for 90 min. After rinsing with PBS three times, organoids were stained with DAPI (1μg/mL; Sigma Aldrich, # D9542) for 10 min. Images were taken using an Olympus FV3000 confocal fluorescence microscope.

#### On-Chip

CRC Organ Chips were washed by flowing PBS through the endothelial and epithelial channels. The chips were fixed with 4% paraformaldehyde (ThermoFisher Scientific, #J61899.AP), incubated for 15 min, and permeabilized with 1% saponin. Blocking buffer of 2% bovine serum albumin (BSA) and primary antibodies were incubated overnight at 4°C before a 2-h incubation with secondary antibodies (1:500, Invitrogen, #A21428 and #A21235) diluted in blocking buffer. The primary antibodies used for the CRC-Chip studies were mouse anti-ZO-1 Alexa Fluor 594 (1:100; Invitrogen #339194), rabbit anti-GABA (1:400; Sigma-Aldrich #A2052), rabbit anti-VE-cadherin (1:25; Abcam, #ab33168), and DAPI (Sigma-Aldrich, #D9542) was used to label all nuclei. Chips were imaged using the Perkin Elmer Operetta CLS High Content System.

#### Tissue Slides

Formalin-fixed paraffin-embedded (FFPE) slides were incubated in a slide warmer for 1 hr at 60°C and further de-paraffinized by the immersion in HistoChoice (Sigma Aldrich, #H2779) for 3 rounds of 5 min each. To rehydrate, the slides were sequentially immersed, for 3 min each, 100% Ethanol, 100% Ethanol, 100% Ethanol, 95% Ethanol, 95% Ethanol, 70% Ethanol, 50% Ethanol, DI water, DI water. The slides were placed in a container with Antigen Retrieval Buffer (1mM EDTA Buffer (Newcomer Supply #1056A), 0.05% Tween 20), heated to boiling, and cooled for 30 min in buffer. Once cooled, the slides were incubated with 1M Glycine for 30 min in a humified chamber and rinsed three times with 1x PBS. The slides were incubated with blocking buffer (2.5% Goat Serum (Vector Laboratories #S-1012-50), 0.05% Tween-20) for 1 hr. Each slide was stained overnight with rabbit anti-GABA (1:400; Sigma-Aldrich A2052) and mouse anti-CK20 (1:100; Abcam, #Ab854 [KS20.8]) in blocking buffer at 4°C followed by three washes with 1x PBS. The slides were incubated with goat anti-rabbit Alexa Fluor 647 (1:100; Invitrogen #A-21235) and goat anti-mouse Alexa Fluor 555 (1:100; Invitrogen, #A21235) in blocking buffer for 2 hr, then rinsed with 1 X PBS three times. The slides were lastly stained with mouse anti-EpCAM FITC antibody (1:100; GeneTex, #GTX30708) in blocking buffer for 2 hr followed by rinsing with 1 X PBS three times. Each slide was mounted with VECTASHIELD Antifade Mounting Medium (Vector Labs, #H1000-10) and sealed with coverslip and clear nail polish. Images were on the Olympus VS120 Slide Scanner.

### RNAseq (Organoid, OOC, and Tumor Tissue)

#### RNA extraction from OOC

Endothelial HIMECs were removed and discarded via Trypsin from the bottom channel prior to collecting organoids. Organoids were lysed using Lysis Buffer (Invitrogen, #46-6001) with 1% 2-mercaptoethanol (Sigma, #M3148) and vigorous pipetting. RNA was isolated using PureLink RNA Mini Kit (Invitrogen, #12183018A) following manufacturer’s instructions.

#### RNA-seq

Sequencing libraries were generated by SMARTer Stranded Total RNAseq Kit – Pico Input Mammalian V3 (Takara Bio, #634485) according to manufacturer’s instructions. Organoid and tumor tissue cDNA synthesis, sample quality assessment, cDNA library preparation, and sample sequencing were performed by the University of Southern California (USC) Genomics Core Facility (organoids and tumor tissue). Samples were sequenced on NextSeq 500 (Illumina). Read length was 75 bp. Organoids-on-chip cDNA synthesis, sample quality assessment, cDNA library preparation and sample sequencing were handled by GENEWIZ from Azenta Life Sciences. Samples were sequenced on HiSeq 2500 (Illumina) rapid run flow cells. Read length was 150 bp.

#### Alignment and Quality Control

Quality of raw reads were evaluated using fastp. STAR was used to align reads against GENCODE v39 annotations (GRCh38). MultiQC was used to summarize statistics from both across all samples. Both duplication and alignment rate were used to select good quality samples. Samples were further inspected for outliers using principal component analysis (PCA) and correlation. Blinded variance stabilizing transformed (VST) values were used as inputs for both PCA and correlation.

Quality control metadata were visualized on the principal components and a heatmap of correlations by sample.

#### Differential Expression and Gene Set Enrichment Analysis

DESeq2 was used to perform differential expression using a negative binomial model with a Wald test to determine significance. Default DESeq2 metrics were used to filter genes based on expression levels and calculate normalization factors. All visualizations of gene expression use values derived from a blind variance stabilizing transformation of raw counts. Volcano plots of results were generated using EnhancedVolcano with an adjusted P value threshold of 0.05 to indicate statistical significance and a foldchange threshold of +/- 2. Results were subsequently processed by gene set enrichment analysis using fgseaMultilevel from the fgsea package with all Hallmark gene sets. Summary visualizations of gene set enrichment results were generated by unsupervised selection of the 10 most significant gene sets in the positive and negative direction. Over enrichment analysis was performed using DAVID functional analysis clustering (DAVID Functional Annotation Bioinformatics Microarray; https://david.ncifcrf.gov) using GO and KEGG gene sets. For this analysis, medium stringency was used (similarity term overlap = 3, similarity threshold = 0.5, group membership = 3, multiple linkage threshold = 0.5, and EASE = 1).

We further visualized gene sets using foldchanges in patients with samples from stretched and not stretched chips. After calculating foldchanges by patient, genes were ranked by p-value derived from a t-test between patients from differing clinical labels. For a given gene set, the 30 most significant genes from the gene set were selected for inclusion in the heatmap. Patients are ordered first by group and subsequently by average foldchange of the top 30 genes within each group. Foldchanges are thresholded to +/- 2.5 for visualization.

### Mass Spectrometry-Based Metabolomics of CRC-on-Chip

#### Metabolite Extraction

An 100 μL aliquot of each sample was extracted with 500 μL extraction solvent (80/20 Methanol/Water, -20 °C) supplemented with 10 μL “quench” standard mix (Cambridge Isotopes Inc. Metabolomics QC Kit, #MSK-QC2). The solution was briefly vortexed and sonicated for 1 min and then incubated for 1 h incubation at −20 °C. Samples were then centrifuged at 13,000 x g for 30 min at 4 °C. Without disturbing the pellet, 450 μL of supernatant were transferred to a new microcentrifuge tube and dried under vacuum centrifugation at room temperature (approximately 2 hr). The dried samples were resuspended in 100 μL of resuspension solvent (50/50/0.1 (v/v) Water/Acetonitrile/Formic Acid) supplemented with 10 μL “resuspension” standard mix (Cambridge Isotopes Inc. Metabolomics QC Kit, #MSK-QC1).

#### Liquid Chromatography with Tandem Mass Spectrometry (LC-MS/MS)

Extracted metabolites were analyzed using an Agilent ultra-high performance liquid chromatography (UHPLC) system (1290 Infinity II HPLC) interfaced with an Agilent quadrupole time of flight mass spectrometer (Q-TOF 6545) equipped with an orthogonal Jet Stream Technology Electrospray Ionization (DUAL AJS-ESI) interface. Samples were separated by 2 different chromatographic separation methods using two analytical columns and conditions. The hydrophilic interaction liquid chromatography (HILIC) with Agilent Poroshell HILIC-Z column (2.1 × 100 mm, 2.7μm) (Agilent InfinityLab Poroshell 120 HILIC-Z, #675775-924) used a two solvent gradient (Mobile Phase A: 10 mM Ammonium Acetate in 90/10 Water/Acetonitrile, pH 9.2, 5 μm Agilent InfinityLab Deactivator Additive; Mobile Phase B: 10 mM Ammonium Acetate in 90/10 Acetonitrile/Water, pH 9.2, 5 μm Agilent InfinityLab Deactivator Additive) at a flow rate of 0.5 mL/min. The column was equilibrated at 98% B for 3 min between runs. Subsequent to sample injection, the ratio was held for 3 min at 98% B after which the solvent ratio was then reduced from 98% B to 50% B over 7 minutes and then brought back up to 98% B over 5 min. The second chromatography was reverse-phase liquid chromatography (RP-HPLC C18) with an Agilent Zorbax RRHD Eclipse Plus C18 column (2.1 × 50 mm, 1.8μm) (Agilent Zorbax RRHD Eclipse Plus C18, #959757-902) which used a two solvent gradient (mobile phase A: water with 0.1% formic acid, B: methanol with 0.1% formic acid as mobile phases A and B, at a flowrate of 0.3 mL/min). The C18 column was equilibrated at 2% B and a sample was injected and held for 3 min at 2% B after which the solvent ratio was increased from 2% B to 98% B over 12 min, held at 98% B for 2 min and then reduced back to 2% B over 2 min. For both HILIC and RP-HPLC C18 chromatographic separation, data were collected in positive and negative ion mode. The injection volume was 2 μL and the column temperature was set at 30 °C. Data were acquired from 50 to 1250 m/z at 1 spectra/s. The jet stream technology electrospray ionization (AJS-ESI) parameters were: gas temperature 290°C (325°C for C18), drying gas 9 L/min, nebulizer 35 psi, fragmentor 125 V, sheath gas temperature 350°C, sheath gas flow 11 L/min at 1000V nozzle voltage.

#### Metabolite Data Analysis

The LC-MS/MS data were processed by Agilent Mass Hunter Workstation Data Acquisition (.d) and analyzed by Agilent Mass Hunter Profinder for batch recursive feature extraction. For both targeted and untargeted analysis, spectral peak extraction was performed with a minimum peak height of 500 counts and charge state of one. Retention time and mass alignment corrections were performed on the runs to remove non-reproducible signals. For untargeted analysis, the resulting features were then exported (.cef) to Agilent Technologies Mass Profiler Professional (MPP) software for multivariate analysis. Principal Component Analysis (PCA) was performed to check the quality of the samples and then the data containing filtered features were processed by unpaired t-test to find the difference between the groups among the filtered data. Analytes with p values < 0.05 and fold change > 2 were regarded as statistically significant. Additionally, reported p-values were adjusted per the Benjamini-Hochberg method of multiple test correction. Feature identification was performed by matching their m/z values and retention times (when available) with in-house libraries of measured values for the IROA technologies MSMLS compound library. Metabolites that did not match to our internal library were matched against METLIN library or were listed with a predicted molecular formula.

Additional targeted feature extraction was performed against a custom library of neurotransmitters using Profinder as described above. Features were exported to CSV for manual analysis by Excel.

### GABA Metabolite Measurements

For sample preparation, 1 mg/mL of GABA (Sigma Aldrich #A2129), or ^13^C_4_ GABA (Cambridge Isotope Laboratories, Inc. #CLM-8666-PK) was added to the top channel inlet media and flown through the channel over the course of the six-day experiment. Media was replaced on day 3 and cells were harvested from the top channel of the CRC tumor chips and processed via mass spectrometry-based metabolomics described in the previous section.

### Invasion Assay

Invasion was monitored and quantified as previously described^37, 71^. For GABA and ABAT inhibition studies, the following drugs were perfused through the top channel starting on day 0. GABA (1 μg/mL; Sigma Aldrich #A2129) and Vigabatrin (10 mM; Sigma #V8261). Inlet media with the GABA or ABAT inhibitor was refreshed on day 3 of the experiment.

### Tumor Cell Transwell Invasion Assay

FluoroblokTM transwell inserts with PET membranes (6.5 mm membrane diameter, 8 μm membrane pore size, 0.3 cm^2^ cell culture area, Corning, #351152) were coated with 30 μg mL-1 type I collagen (Corning, #C354249) and 100 μg mL-1 Matrigel (Corning, #356231) for 2 hr at 37°C before the ECM was gently aspirated from the insert. The inserts were placed in wells with 500 μL endothelial media in the bottom chamber, and CRC tumor cell lines (DLD1 KRAS WT GFP or DLD1 KRAS Mutant GFP) were added (2×10^5^ cells in 200 μL; 6.7×10^5^ cells/cm). Cells were incubated in the same cell culture media as organ-chip experiments in order to facilitate comparisons between experiments. CRC tumor cells in the top chamber were maintained in DMEM with 10% FBS and 1% Pen-Strep. Fully supplemented EBM-2 media (2% FBS, 1% Pen-Strep, Hydrocortisone, hFGF-B, VEGF, R3-IGF-1, Ascorbic Acid, hEGF, and heparin in proprietary concentrations) was placed in the bottom of the wells. 48 hr after tumor cell seeding, cells were stained with Hoechst 33342 fluorescent nuclear stain (1 mg/mL; Invitrogen #H3570) for 10 minutes. The transwells were imaged using the Perkin Elmer Operetta High Content Screening (HCS) platform. The number of cells in the bottom chamber were quantified using Perkin Elmer Harmony software.

### CTC Bulk RNAseq

Cells were collected from the endothelial effluent of stretched and not stretched CRC-Chips as previously described^37^. Briefly, invaded circulated tumor cells (CTCs) found in the endothelial effluent were collected after 48 hr of flow and cultured for down-stream analysis. Cells from 2 chips were pooled together to obtain enough cells for RNA sequencing. RNA extraction, cDNA synthesis, sample quality assessment, cDNA library preparation, and sample sequencing were performed by GENEWIZ from Azenta Life Sciences. Samples were sequenced on HiSeq 2500 (Illumina) rapid run flow cells. Read length was 150 bp. Reads were mapped on reference human genome GRch38. Quality control was performed using FastQC software. Gene expression quantification was performed using RSEM algorithms. Gene sets were identified from a subset of genes that either had a 2-fold difference between stretched and non-stretched or a false discovery rate (FDR)-adjusted p-value <0.1. Here, the alpha level was increased to 0.1 from 0.05 to enable discovery with small sample sizes.

### Neurotransmitter qPCR Array

Cellular RNA was extracted using RNAspin Mini RNA Isolation Kit (GE Healthcare; #25-0500-71) and cDNA was reverse transcribed using iScript Reverse Transcription Supermix (Bio-Rad, #1708841) following manufacturer’s instructions. TaqMan^TM^ Human Neurotransmitter PCR Array (ThermoFisher Scientific, #4414094) was used to detect the expression of neurotransmitter-and GABA-related genes in GFP+ HCT116 harvested from the top epithelial channel of stretched and not stretched chips and sorted via flow cytometry. The array includes 84 genes known to be involved in neurotransmitter synthesis, transport, and degradation, including GABAergic genes. Results were normalized to the average Cq values of 5 housekeeping genes. Chips from three independent chip experiments were used for this analysis.

### RT-qPCR

To interrogate GAD1 expression in the organoid-CRC-Chips, RT-qPCR analysis was performed on organoids collected from the chip effluent. Cellular RNA was extracted as outlined above (RNAseq) and cDNA was reverse transcribed using iScript Reverse Transcription Supermix (Bio-Rad, #1708841) following manufacturer’s instructions. The cDNA was then amplified using iScript SYBR Green Master Supermix or iScript SsoAdvanced^TM^ SYBER Green Master Supermix (Bio-Rad; #1708880 or #1725271). Human GAD1 PrimePCR primers were purchased from BioRad and the sequence for human GAPDH primers are as follows: F 5’TCTGGTAAAGTGGATATTGTTG3’; F 5’GATGGTGATGGGATTTCC3’. Results were normalized to GAPDH expression for all experiments.

### Western Blot

Total protein was extracted and lysed in RIPA buffer with 1x Halt^TM^ protease and phosphatase inhibitor cocktail (Thermo Scientific Cat# 1861282) and protein concentrations were determined using the Pierce^TM^ BCA Protein Assay Kit (Thermo-Fisher, #23225). 20 mg of protein lysates were boiled in SDS loading buffer (Life Technologies, #B007). Protein samples were loaded and fractionated on a 4-12% Bis-Tris gel (Life Technologies, #NW04122BOX) using 1xBolt^TM^ MES SDS Running Buffer (Thermo-Fisher, #B0002) and then transferred to a polyvinylidene difluoride membrane. The membrane was blocked in 5% bovine serum albumin (BSA) in Tris-buffered saline with 0.1% Tween-20 (TBS-T buffer) and primary antibodies against total protein were used. Secondary antibody (mouse IgG-HRP; 1:5000; Cytiva, #NA931) was visualized using an enhanced chemiluminescent horseradish peroxidase substrate. A protein ladder was used to determine molecular mass and ImageJ was used to measure intensity of the blot. The primary antibodies used for the HCT116 H2B-GFP ABAT knock down validation were mouse anti-ABAT (1:500; Santa Cruz Technology, Inc., #sc-393769) and mouse anti-Actin (1:10000; Millipore Sigma, #A5441).

### TCGA Data Analysis

RNA-seq data from The Cancer Genome Atlas (TCGA) colonic adenocarcinoma database (TCGA-COAD) was extracted from the cBioportal for Cancer Genomics (www.cbioportal.org). CRC cases were selected based on any KRAS, NRAS, or BRAF mutations and any treatment status was included. The expression of GAD1 in the KRAS mutant tumors versus KRAS WT tumors in the primary colon tumor was analyzed. All KRAS, NRAS, or BRAF mutant CRC tumors were stratified based on GAD1 expression, with “high” and “low” GAD1 expression designated as above or below the median expression for the cohort (specifically, median mRNA expression z-scores relative to all samples = 0.3). The prognosis of each group was examined using Kaplan-Meier survival estimators and significance was assessed using a log-rank (Mantel-Cox test).

### Quantification and Statistical Analysis

Unless otherwise notes, all experiments were performed with 2-3 chips per condition and repeated independently at least 2 times. Analysis of variance (ANOVA) and t-tests were performed using GraphPad Prism 9 software, with the p-value < 0.0001: ****; p-value < 0.001: ***; p-value < 0.01: **; p-value < 0.05: *, as noted in all figure legends. Unless otherwise noted, individual data points are reported and shown in all figures and the mean with standard error of the mean (SEM) is represented.

