## Supplemental Information for "Integration of Patient-Derived Organoids and Organ-on-Chip Systems: Investigating Colorectal Cancer Invasion within the Mechanical and GABAergic Tumor Microenvironment"

Supplementary Information

**Table S1**. GSEA of Hallmark gene sets on gene expression comparisons between model systems.

|  | Chip vs Tumor | Organoid vs Tumor | Organoid vs Chip |
| --- | --- | --- | --- |
|  | padj | padj | padj |
| Hm Adipogenesis | NA | NA | 0.264057161 |
| Hm Allograft Rejection | 2.41657E-09 | 1.49464E-09 | 0.64778217 |
| Hm Androgen Resp | 0.001375175 | 1 | 0.000715273 |
| Hm Angiogenesis | 0.000106823 | 1.5145E-08 | 0.31075419 |
| Hm Apical Junction | 3.18594E-06 | 8.96863E-09 | 0.172655051 |
| Hm Apical Surface | 0.30190678 | 0.375251762 | 0.78808522 |
| Hm Apoptosis | 0.183694084 | 2.19838E-06 | 0.000862262 |
| Hm Bile Acid Met | 0.802606449 | 0.555367709 | 0.172655051 |
| Hm Cholesterol Homeostasis | 1.53697E-06 | NA | 0.028933986 |
| Hm Coagulation | 7.48534E-08 | 6.44617E-14 | 0.031040601 |
| Hm Complement | 0.000302455 | 5.5395E-09 | 0.001091401 |
| Hm Dna Repair | 0.627273073 | NA | 0.066059894 |
| Hm E2f Targets | NA | NA | 0.788351191 |
| Hm Epithelial Mesenchymal Transition | 9.72441E-28 | 4.27429E-37 | 0.027690196 |
| Hm Estrogen Resp Early | NA | 1 | 3.47062E-05 |
| Hm Estrogen Resp Late | NA | 1 | 0.000515551 |
| Hm Fatty Acid Met | 5.5836E-07 | NA | 0.00190008 |
| Hm G2m Checkpoint | NA | NA | 0.272785623 |
| Hm Glycolysis | NA | NA | 0.305029109 |
| Hm Hedgehog Sig | 0.280181723 | 0.027814244 | 0.509762111 |
| Hm Heme Met | NA | 1 | 0.000933759 |
| Hm Hypoxia | 0.131282737 | 0.000313828 | 0.166672849 |
| Hm Il2 Stat5 Sig | 0.001375175 | 0.000313828 | 0.376299945 |
| Hm Il6 Jak Stat3 Sig | 0.000171913 | 1.15632E-06 | 0.248015873 |
| Hm Inflammatory Resp | 1.64415E-10 | 7.68929E-14 | 0.000704096 |
| Hm Interferon Alpha Resp | 0.042532166 | 0.000795422 | 0.12292527 |
| Hm Interferon Gamma Resp | 1.69036E-07 | 6.58716E-10 | 0.066059894 |
| Hm Kras Sig Dn | 0.000487285 | 2.88006E-06 | 0.226150725 |
| Hm Kras Sig Up | 2.68685E-07 | 9.53694E-15 | 0.028933986 |
| Hm Mitotic Spindle | NA | NA | 0.826044704 |
| Hm Mtorc1 Sig | NA | NA | 0.001646747 |
| Hm Myc Targets V1 | NA | NA | 0.127876589 |
| Hm Myc Targets V2 | 0.063018851 | NA | 0.361599546 |
| Hm Myogenesis | 2.68685E-07 | 6.54177E-11 | 0.165636593 |
| Hm Notch Sig | 0.618489583 | 0.603612815 | 0.826044704 |
| Hm Oxidative Phosphorylation | NA | NA | 0.028933986 |
| Hm P53 Path | 0.928857715 | 1 | 0.132903901 |
| Hm Pancreas Beta Cells | 0.562102851 | 0.191295547 | 0.165898618 |
| Hm Peroxisome | 0.000527023 | NA | 0.044957521 |
| Hm Pi3k Akt Mtor Sig | 0.005531386 | NA | 0.06058597 |
| Hm Protein Secretion | 5.70711E-11 | NA | 3.47368E-06 |
| Hm Reactive Oxygen Species Path | 0.095244923 | NA | 0.053888021 |
| Hm Spermatogenesis | 0.694141486 | 0.411428571 | 0.994818653 |
| Hm Tgf Beta Sig | 0.78754118 | 0.000313828 | 0.000821472 |
| Hm Tnfa Sig Via Nfkb | 5.47778E-06 | 3.90933E-12 | 3.47368E-06 |
| Hm Unfolded Protein Resp | 3.58882E-16 | NA | 7.12266E-05 |
| Hm Uv Resp Dn | 2.60084E-06 | 1.98788E-11 | 0.002054662 |
| Hm Uv Resp Up | 0.056137357 | 0.342570473 | 0.00413105 |
| Hm Wnt Beta Catenin Sig | 0.825114168 | 0.042727665 | 0.737311385 |
| Hm Xenobiotic Met | NA | 1 | 0.004219087 |

**Table S2**. Metabolites measured in epithelial effluent on D0 and D6, related to PCA in Figure 2E; C18 modes.

| **C18 Positive** | | | **C18 Negative** | | |
| --- | --- | --- | --- | --- | --- |
| **Compound (ID)** | **Mass** | **RT** | **Compound (ID)** | **Mass** | **RT** |
| (2R,4R)-tert-butyl 4-(hydroxymethyl)-2-phenylthiazolidine-3-carboxylate | 295.1257 | 5.7349987 | (+)-Pimaric acid | 302.2226 | 15.265993 |
| (3beta,8beta)-3-Hydroxy-7(11)-eremophilen-12,8-olide | 250.1566 | 12.942009 | (E,E)-3,7,11-Trimethyl-2,6,10-dodecatrienyl pentanoate | 306.2543 | 15.727019 |
| (4OH,8Z,t18:1) sphingosine | 315.2782 | 14.505988 | (Z,Z)-2-Methyl-5-(8,11,14-pentadecatrienyl)-1,3-benzenediol | 328.2394 | 15.50402 |
| (E,E)-3,7,11-Trimethyl-2,6,10-dodecatrienyl pentanoate | 306.251 | 16.130993 | 1734.711@9.51701 | 1734.711 | 9.51701 |
| (ent-16betaOH)-16,17-Dihydroxy-9(11)-kauren-19-oic acid | 334.2129 | 13.526985 | 2-Methyl-5-(8-pentadecenyl)-1,3-benzenediol | 332.2697 | 15.873016 |
| (R)-Apiumetin | 244.0704 | 11.2159815 | 2-Methyl-5-(8,11-pentadecadienyl)-1,3-benzenediol | 330.2541 | 15.646981 |
| (R)-Humulone | 362.209 | 10.706988 | 2-O-Caffeoyltartronic acid | 282.0336 | 0.77799994 |
| (S)-a-Amino-2,5-dihydro-5-oxo-4-isoxazolepropanoic acid N2-glucoside | 334.0989 | 10.34599 | 243.1828@11.649009 | 243.1828 | 11.649009 |
| (Z)-2-amino-1-hydroxyoctadec-4-en-3-one | 297.2677 | 15.56201 | 280.0449@1.182 | 280.0449 | 1.182 |
| (Z)-3,7-Dimethyl-2,6-octadienyl formate | 182.1282 | 10.442008 | 2E-Phytenoic acid | 310.2856 | 16.218012 |
| (Z)-Tamarindienal | 126.0309 | 4.513 | 3-(2-Furanyl)-2-propenal | 122.0363 | 9.184009 |
| (Z)-Tamarindienal Esi+3.0130002 | 126.0315 | 3.0130002 | 344.0942@9.130994 | 344.0942 | 9.130994 |
| (Z)-Tamarindienal Esi+4.196998 | 126.032 | 4.196998 | 372.2257@15.526 | 372.2257 | 15.526 |
| (Z)-Tamarindienal Esi+4.514 | 126.0312 | 4.514 | 411.0662@8.732988 | 411.0662 | 8.732988 |
| 1-(2,4,5-Trimethoxyphenyl)-1,2-propanedione | 238.0826 | 7.8120084 | 453.1@6.9569893 | 453.1 | 6.9569893 |
| 1-Methoxy-1H-indole-3-carboxaldehyde | 175.0634 | 10.354003 | 5-Dodecyldihydro-2(3H)-furanone | 254.2226 | 15.386988 |
| 1,3-Butadiene | 54.0467 | 7.752996 | 5,8,10,14-Eicosatetraenoic acid, 12-hydroxy-, [S-(E,Z,Z,Z)]-; 12(S)-5,8,10,14-Hydroxyeicosatetraenoic acid; 12(S)-HETE; 12-HETE; 12-Hydroxy-5,8,10,14-eicosatetraenoic acid; 12-Hydroxyeicosatetraenoic acid; 12-L-Hydroxy-5,8,10,14-eicosatetraenoic acid; 12S | 320.2339 | 14.458987 |
| 1,8-Naphthyridine-3-carboxylic acid, 1-ethyl-1,4-dihydro-7-hydroxy-4-oxo- glucuronide | 410.0971 | 9.196 | 514.1586@9.148991 | 514.1586 | 9.148991 |
| 1018.5271@10.366991 | 1018.5271 | 10.366991 | 525.3391@15.2590065 | 525.3391 | 15.2590065 |
| 1040.2435@10.359992 | 1040.2435 | 10.359992 | 527.3168@14.720979 | 527.3168 | 14.720979 |
| 1057.7478@16.322014 | 1057.7478 | 16.322014 | 541.3357@15.020997 | 541.3357 | 15.020997 |
| 1140.8506@16.306011 | 1140.8506 | 16.306011 | 574.1183@8.966996 | 574.1183 | 8.966996 |
| 1141.8907@10.333007 | 1141.8907 | 10.333007 | 6-Deoxyfagomine | 131.0945 | 1.3209993 |
| 1142.2245@10.333007 | 1142.2245 | 10.333007 | 6,8-Dihydroxypurine | 152.0332 | 1.0999991 |
| 1142.5586@10.330003 | 1142.5586 | 10.330003 | 609.3208@15.020997 | 609.3208 | 15.020997 |
| 1145.8016@16.306995 | 1145.8016 | 16.306995 | Acetylenic acids; 17-Octadecen-9-ynoic acid | 278.2227 | 15.277987 |
| 1189.8269@16.300022 | 1189.8269 | 16.300022 | Asp His Lys | 398.1917 | 11.485994 |
| 120.0213@9.192994 | 120.0213 | 9.192994 | Benzyl b-L-arabinopyranoside | 238.1191 | 11.270997 |
| 1233.8533@16.290018 | 1233.8533 | 16.290018 | C20:4n-2,6,9,12 | 304.2395 | 15.525016 |
| 1277.8853@16.284018 | 1277.8853 | 16.284018 | DAPT | 432.1886 | 13.078987 |
| 12R-HOME(13Z) | 298.2507 | 14.906005 | Diethyl decanedioate | 258.1818 | 13.588991 |
| 1321.907@16.272976 | 1321.907 | 16.272976 | Ferulic acid | 194.0566 | 9.185001 |
| 133.9952@0.7210005 | 133.9952 | 0.7210005 | Folic acid | 441.1376 | 6.908997 |
| 13E-Docosenamide | 337.335 | 15.312997 | Gamma-glutamyl-Alanine | 217.1052 | 0.6500006 |
| 14,15-HxA3 (11S) | 336.2281 | 14.197014 | His Ala Asn | 340.1502 | 11.643995 |
| 1409.96@16.256992 | 1409.96 | 16.256992 | Integerrine | 593.3015 | 15.020013 |
| 144.0093@9.013991 | 144.0093 | 9.013991 | Isoferuloyl C1-glucuronide | 370.0881 | 8.244005 |
| 148.016@8.161004 | 148.016 | 8.161004 | Melleolide D | 482.1647 | 1.1660002 |
| 148.0163@9.192994 | 148.0163 | 9.192994 | Meta-Tyrosine | 181.0737 | 1.1620013 |
| 159.1622@5.948995 | 159.1622 | 5.948995 | methoxyacetic acid | 90.0318 | 0.83499926 |
| 1605.6794@9.604003 | 1605.6794 | 9.604003 | Methyl 3-(2,3-dihydroxy-3-methylbutyl)-4-hydroxybenzoate | 254.114 | 9.265011 |
| 1627.6619@9.603016 | 1627.6619 | 9.603016 | Nicarbazin | 302.0624 | 4.0599985 |
| 1734.7205@9.503005 | 1734.7205 | 9.503005 | PA(15:0/0:0) | 396.2251 | 15.505004 |
| 1756.7021@9.501999 | 1756.7021 | 9.501999 | Pseudouridine | 244.0681 | 1.1760007 |
| 185.9622@9.187992 | 185.9622 | 9.187992 |  |  |  |
| 188.0619@10.347004 | 188.0619 | 10.347004 |  |  |  |
| 1952.8643@15.220986 | 1952.8643 | 15.220986 |  |  |  |
| 2-(8-[3]-ladderane-octanyl)-sn-glycero-3-phosphoethanolamine | 487.3048 | 14.938989 |  |  |  |
| 2-Acetylfuran | 110.0369 | 13.604986 |  |  |  |
| 2-Amino-5-phosphopentanoic acid | 197.0455 | 10.350994 |  |  |  |
| 2-C-Methyl-D-erythritol 4-phosphate | 216.0405 | 9.194994 |  |  |  |
| 2-Ethylthiazole | 113.029 | 10.349986 |  |  |  |
| 2-Oxo-4-methylthiobutanoic acid | 148.0164 | 11.2159815 |  |  |  |
| 2,2'-(3-methylcyclohexane-1,1-diyl)diacetic acid | 214.1187 | 7.580004 |  |  |  |
| 2,2'-(3-methylcyclohexane-1,1-diyl)diacetic acid Esi+7.262009 | 214.1187 | 7.262009 |  |  |  |
| 2,3-Dihydro-3-hydroxy-6-methoxy-2,2-dimethyl-4H-1-benzopyran-4-one | 222.0895 | 11.2159815 |  |  |  |
| 2,3-Dihydro-5-(3-hydroxypropanoyl)-1H-pyrrolizine | 179.0945 | 6.5800037 |  |  |  |
| 2,3-Dihydroabscisic alcohol | 252.1703 | 12.000985 |  |  |  |
| 2,3,4,5,2',3',4',6'-Octamethoxychalcone | 448.1715 | 11.686999 |  |  |  |
| 2,4-Dihydroxyacetophenone 5-sulfate | 232.0051 | 9.189992 |  |  |  |
| 2037.0765@10.675008 | 2037.0765 | 10.675008 |  |  |  |
| 2037.0768@10.67899 | 2037.0768 | 10.67899 |  |  |  |
| 2037.5782@10.682006 | 2037.5782 | 10.682006 |  |  |  |
| 2037.5792@10.682006 | 2037.5792 | 10.682006 |  |  |  |
| 218.1805@12.578988 | 218.1805 | 12.578988 |  |  |  |
| 2189.9873@9.470996 | 2189.9873 | 9.470996 |  |  |  |
| 225.1732@11.6689825 | 225.1732 | 11.6689825 |  |  |  |
| 2291.9253@9.47301 | 2291.9253 | 9.47301 |  |  |  |
| 238.0215@9.187007 | 238.0215 | 9.187007 |  |  |  |
| 239.2284@12.583007 | 239.2284 | 12.583007 |  |  |  |
| 243.1842@11.670013 | 243.1842 | 11.670013 |  |  |  |
| 245.1461@12.301016 | 245.1461 | 12.301016 |  |  |  |
| 25-Acetyl-6,7-didehydrofevicordin F 3-glucoside | 704.3378 | 15.228021 |  |  |  |
| 260.0457@11.21801 | 260.0457 | 11.21801 |  |  |  |
| 262.0331@9.19 | 262.0331 | 9.19 |  |  |  |
| 268.22@15.386987 | 268.22 | 15.386987 |  |  |  |
| 287.2099@9.844982 | 287.2099 | 9.844982 |  |  |  |
| 3-(3,4-Methylenedioxyphenyl)propenal | 176.0467 | 11.2159815 |  |  |  |
| 3-Acetyldihydro-2(3H)-furanone | 128.0476 | 13.604986 |  |  |  |
| 3-methyl-thiopropionic acid | 120.0209 | 11.214996 |  |  |  |
| 3-Oxochola-1,4,6-trien-24-oic Acid | 368.2335 | 14.600983 |  |  |  |
| 3,4-Dimethylmethcathinone | 190.1571 | 11.081016 |  |  |  |
| 3,5,8-Trimethoxy-3',4'-methylenedioxy-7-prenyloxyflavone | 440.1415 | 11.9100065 |  |  |  |
| 3,5,8-Trimethoxy-3',4'-methylenedioxy-7-prenyloxyflavone Esi+12.286009 | 440.1434 | 12.286009 |  |  |  |
| 3''-Chloro-3''-deoxytriphasiol | 366.1249 | 10.201009 |  |  |  |
| 3''-Chloro-3''-deoxytriphasiol Esi+10.202015 | 366.1252 | 10.202015 |  |  |  |
| 315.2417@11.432006 | 315.2417 | 11.432006 |  |  |  |
| 317.1842@11.899013 | 317.1842 | 11.899013 |  |  |  |
| 330.1044@0.639 | 330.1044 | 0.639 |  |  |  |
| 342.2636@15.725983 | 342.2636 | 15.725983 |  |  |  |
| 356.0802@8.758005 | 356.0802 | 8.758005 |  |  |  |
| 356.0815@8.758005 | 356.0815 | 8.758005 |  |  |  |
| 367.4186@15.560031 | 367.4186 | 15.560031 |  |  |  |
| 372.053@10.349986 | 372.053 | 10.349986 |  |  |  |
| 376.0153@6.8540034 | 376.0153 | 6.8540034 |  |  |  |
| 376.0362@9.015011 | 376.0362 | 9.015011 |  |  |  |
| 386.1869@13.094979 | 386.1869 | 13.094979 |  |  |  |
| 391.9939@9.1900015 | 391.9939 | 9.1900015 |  |  |  |
| 397.32@13.607022 | 397.32 | 13.607022 |  |  |  |
| 399.3361@13.989992 | 399.3361 | 13.989992 |  |  |  |
| 4-[[5-(acetylamino)-1-methyl-1H-indol-3-yl]methyl]-3-methoxy-N-[(2-methylphenyl)sulfonyl]-Benzamide | 505.1646 | 11.439981 |  |  |  |
| 4-Hydroxy-5-phenyltetrahydro-1,3-oxazin-2-one | 193.0747 | 6.6929946 |  |  |  |
| 4-Vinylcyclohexene | 108.0933 | 8.653002 |  |  |  |
| 4-Vinylcyclohexene Esi+7.74799 | 108.0933 | 7.74799 |  |  |  |
| 4,5-Dimethyl-4-hexen-3-one | 126.1039 | 7.7479897 |  |  |  |
| 402.0742@10.350996 | 402.0742 | 10.350996 |  |  |  |
| 403.1997@11.985991 | 403.1997 | 11.985991 |  |  |  |
| 407.3616@15.477023 | 407.3616 | 15.477023 |  |  |  |
| 407.9685@9.189993 | 407.9685 | 9.189993 |  |  |  |
| 408.0456@8.294004 | 408.0456 | 8.294004 |  |  |  |
| 408.046@8.282996 | 408.046 | 8.282996 |  |  |  |
| 409.1137@6.3530097 | 409.1137 | 6.3530097 |  |  |  |
| 411.0692@8.63599 | 411.0692 | 8.63599 |  |  |  |
| 411.2873@15.281004 | 411.2873 | 15.281004 |  |  |  |
| 415.0875@8.606007 | 415.0875 | 8.606007 |  |  |  |
| 420.2762@16.744009 | 420.2762 | 16.744009 |  |  |  |
| 423.3359@13.833989 | 423.3359 | 13.833989 |  |  |  |
| 424.3552@16.823 | 424.3552 | 16.823 |  |  |  |
| 429.3085@10.697999 | 429.3085 | 10.697999 |  |  |  |
| 433.0519@8.632997 | 433.0519 | 8.632997 |  |  |  |
| 435.1787@1.7689983 | 435.1787 | 1.7689983 |  |  |  |
| 444.2265@11.985991 | 444.2265 | 11.985991 |  |  |  |
| 453.1035@6.8170033 | 453.1035 | 6.8170033 |  |  |  |
| 468.0988@2.6170018 | 468.0988 | 2.6170018 |  |  |  |
| 479.3385@15.275016 | 479.3385 | 15.275016 |  |  |  |
| 481.3539@15.337012 | 481.3539 | 15.337012 |  |  |  |
| 483.1822@11.447013 | 483.1822 | 11.447013 |  |  |  |
| 485.1295@9.505002 | 485.1295 | 9.505002 |  |  |  |
| 486.3612@15.232022 | 486.3612 | 15.232022 |  |  |  |
| 5-Aminoimidazole-4-carboxamide-1-β-D-ribofuranosyl 5'-monophosphate | 338.0603 | 6.856005 |  |  |  |
| 5,7,3',4',5'-Pentahydroxy-3,6,8-trimethoxyflavone | 392.0719 | 8.236007 |  |  |  |
| 5,7,3',4',5'-Pentahydroxy-3,6,8-trimethoxyflavone Esi+8.182997 | 392.0724 | 8.182997 |  |  |  |
| 5,7,3',4',5'-Pentahydroxy-3,6,8-trimethoxyflavone Esi+8.294004 | 392.0716 | 8.294004 |  |  |  |
| 5,8,10,14-Eicosatetraenoic acid, 12-hydroxy-, [S-(E,Z,Z,Z)]-; 12(S)-5,8,10,14-Hydroxyeicosatetraenoic acid; 12(S)-HETE; 12-HETE; 12-Hydroxy-5,8,10,14-eicosatetraenoic acid; 12-Hydroxyeicosatetraenoic acid; 12-L-Hydroxy-5,8,10,14-eicosatetraenoic acid; 12S | 320.2336 | 14.68799 |  |  |  |
| 5′-Deoxy-5′-(methylthio)adenosine | 297.0895 | 5.9990044 |  |  |  |
| 501.3202@15.280022 | 501.3202 | 15.280022 |  |  |  |
| 507.3671@15.458006 | 507.3671 | 15.458006 |  |  |  |
| 512.528@17.273998 | 512.528 | 17.273998 |  |  |  |
| 521.3508@15.182006 | 521.3508 | 15.182006 |  |  |  |
| 530.3779@15.238014 | 530.3779 | 15.238014 |  |  |  |
| 532.0417@10.352006 | 532.0417 | 10.352006 |  |  |  |
| 533.2909@15.03501 | 533.2909 | 15.03501 |  |  |  |
| 539.4403@15.489012 | 539.4403 | 15.489012 |  |  |  |
| 548.0107@10.35499 | 548.0107 | 10.35499 |  |  |  |
| 549.155@13.668 | 549.155 | 13.668 |  |  |  |
| 549.3818@15.621979 | 549.3818 | 15.621979 |  |  |  |
| 549.8687@18.839 | 549.8687 | 18.839 |  |  |  |
| 552.5198@17.276 | 552.5198 | 17.276 |  |  |  |
| 571.3638@15.277984 | 571.3638 | 15.277984 |  |  |  |
| 572.1967@8.821001 | 572.1967 | 8.821001 |  |  |  |
| 573.3783@15.609991 | 573.3783 | 15.609991 |  |  |  |
| 583.4654@15.555018 | 583.4654 | 15.555018 |  |  |  |
| 592.5624@15.833006 | 592.5624 | 15.833006 |  |  |  |
| 596.1037@9.066004 | 596.1037 | 9.066004 |  |  |  |
| 596.1046@8.611004 | 596.1046 | 8.611004 |  |  |  |
| 6-HydroxyKetanserin | 411.1558 | 5.2069926 |  |  |  |
| 6,8-Dihydroxy-1,7-diprenylxanthone-2-carboxylic acid | 408.1548 | 11.987024 |  |  |  |
| 604.4166@15.066015 | 604.4166 | 15.066015 |  |  |  |
| 611.2136@8.041993 | 611.2136 | 8.041993 |  |  |  |
| 612.0706@8.635004 | 612.0706 | 8.635004 |  |  |  |
| 615.1701@12.401988 | 615.1701 | 12.401988 |  |  |  |
| 627.4925@15.567003 | 627.4925 | 15.567003 |  |  |  |
| 653.3262@15.2220125 | 653.3262 | 15.2220125 |  |  |  |
| 657.5036@15.253018 | 657.5036 | 15.253018 |  |  |  |
| 658.2836@15.22301 | 658.2836 | 15.22301 |  |  |  |
| 671.5187@15.568984 | 671.5187 | 15.568984 |  |  |  |
| 674.2557@15.227023 | 674.2557 | 15.227023 |  |  |  |
| 690.1793@10.357012 | 690.1793 | 10.357012 |  |  |  |
| 7-Methylrosmanol | 360.1934 | 10.434007 |  |  |  |
| 706.1448@10.357003 | 706.1448 | 10.357003 |  |  |  |
| 720.4954@15.990988 | 720.4954 | 15.990988 |  |  |  |
| 723.9726@10.352001 | 723.9726 | 10.352001 |  |  |  |
| 725.4367@10.772998 | 725.4367 | 10.772998 |  |  |  |
| 726.2692@15.223014 | 726.2692 | 15.223014 |  |  |  |
| 730.4878@15.989992 | 730.4878 | 15.989992 |  |  |  |
| 7alpha,25-dihydroxycholesterol | 418.3462 | 17.53701 |  |  |  |
| 8-Propyloxycaffeine | 252.1323 | 10.356006 |  |  |  |
| 828.4441@10.351 | 828.4441 | 10.351 |  |  |  |
| 90.0468@6.693987 | 90.0468 | 6.693987 |  |  |  |
| 92.0253@11.218995 | 92.0253 | 11.218995 |  |  |  |
| 947.5443@10.776989 | 947.5443 | 10.776989 |  |  |  |
| 968.022@9.132007 | 968.022 | 9.132007 |  |  |  |
| 969.53@10.784993 | 969.53 | 10.784993 |  |  |  |
| 994.687@15.036993 | 994.687 | 15.036993 |  |  |  |
| 9Z,12E-Octadecadienoic acid | 280.2407 | 14.68601 |  |  |  |
| 9Z,12E-Octadecadienoic acid Esi+15.558987 | 280.2402 | 15.558987 |  |  |  |
| Acetylenic acids; 17-Octadecen-9-ynoic acid | 278.2241 | 14.27298 |  |  |  |
| Acetylenic acids; 17-Octadecen-9-ynoic acid Esi+15.286008 | 278.2249 | 15.286008 |  |  |  |
| Adenine | 135.0547 | 5.998995 |  |  |  |
| Alfuzosin | 389.2042 | 9.41499 |  |  |  |
| Aliskiren | 551.3961 | 15.619999 |  |  |  |
| Amastatin | 474.2691 | 8.463008 |  |  |  |
| Arachidonoyl Serotonin | 462.3231 | 17.670004 |  |  |  |
| Arg Asp Lys | 417.2358 | 9.986008 |  |  |  |
| Arg Tyr Gln | 465.2365 | 11.020997 |  |  |  |
| Azafrin | 426.2776 | 15.381984 |  |  |  |
| Benzyl b-L-arabinopyranoside | 238.1171 | 11.272986 |  |  |  |
| beta-Funaltrexamine | 454.2116 | 11.989011 |  |  |  |
| Blasticidin S | 422.1996 | 7.6950064 |  |  |  |
| Butoctamide hydrogen succinate | 315.2042 | 6.3590055 |  |  |  |
| Cappariloside B | 496.1705 | 7.4990067 |  |  |  |
| CAY10580 | 341.2571 | 12.057004 |  |  |  |
| Cephamycin C | 446.1118 | 10.522996 |  |  |  |
| CerP(d18:1/8:0) | 505.3545 | 15.395012 |  |  |  |
| CHAETOCHROMIN | 546.1572 | 13.750989 |  |  |  |
| Chlorogenoquinone | 352.0804 | 8.140997 |  |  |  |
| Choline chloride | 103.0995 | 0.60100013 |  |  |  |
| Cinnamyl propionate | 190.0981 | 10.350995 |  |  |  |
| Citiolone | 159.035 | 10.340985 |  |  |  |
| Citiolone Esi+8.73901 | 159.0352 | 8.73901 |  |  |  |
| Citiolone Esi+8.741007 | 159.0344 | 8.741007 |  |  |  |
| Citreoviridinol A1 | 420.1768 | 11.382998 |  |  |  |
| Colupulone | 400.2597 | 15.248017 |  |  |  |
| CUDA | 340.2609 | 15.009009 |  |  |  |
| Cycasin | 252.0974 | 9.395997 |  |  |  |
| Cyromazine | 166.0974 | 7.747993 |  |  |  |
| Daidzein 7-O-glucuronide | 430.088 | 11.561005 |  |  |  |
| Delphinidin 3-(acetylglucoside) | 507.1119 | 9.536998 |  |  |  |
| Diethyl decanedioate | 258.1842 | 13.604986 |  |  |  |
| Dihydro Isorescinnamine | 636.3013 | 15.2220125 |  |  |  |
| Dihydro-3-methyl-2(3H)-furanone | 100.0525 | 13.604986 |  |  |  |
| Dimefuron | 338.113 | 11.895011 |  |  |  |
| Eicosapentaenoic Acid methyl ester | 316.2407 | 16.05502 |  |  |  |
| epi-Tulipinolide diepoxide | 322.1409 | 11.892981 |  |  |  |
| Eprosartan | 424.1424 | 13.096994 |  |  |  |
| Ethyl 4-(acetylthio)butyrate | 190.069 | 0.5159995 |  |  |  |
| exo-Dehydrochalepin | 296.1382 | 13.60302 |  |  |  |
| Ferulic acid | 194.0577 | 9.190998 |  |  |  |
| Fluconazole | 306.1041 | 10.350997 |  |  |  |
| Geranyl acetoacetate | 238.1547 | 12.659022 |  |  |  |
| Gibberellin A87 | 362.1354 | 11.478006 |  |  |  |
| Gingerenone B | 386.1733 | 11.985991 |  |  |  |
| Helinorbisabone | 250.1189 | 12.864008 |  |  |  |
| Hercynine | 198.1235 | 8.812992 |  |  |  |
| Himbacine | 345.2677 | 15.510008 |  |  |  |
| His Met Gly | 343.1338 | 6.9229913 |  |  |  |
| His Phe Ile | 415.2197 | 9.439008 |  |  |  |
| Kamahine C | 268.1301 | 10.062009 |  |  |  |
| L-Menthyl acetoacetate | 240.1708 | 11.319984 |  |  |  |
| L-Xylulose | 150.0541 | 4.4650016 |  |  |  |
| Lauroyl diethanolamide | 287.2468 | 13.661009 |  |  |  |
| Leu Glu Gly | 317.1541 | 6.9249926 |  |  |  |
| Leucyl-Glutamate | 260.1331 | 5.778996 |  |  |  |
| Levoamine (Chloramphenicol D base) | 212.0816 | 10.345991 |  |  |  |
| Lys Val Trp | 431.2519 | 10.512994 |  |  |  |
| Lys Val Trp Esi+10.640995 | 431.2519 | 10.640995 |  |  |  |
| Lys-Asn-OH | 368.1319 | 11.891999 |  |  |  |
| Melosatin A | 353.1613 | 0.6539995 |  |  |  |
| Met-Trp-OH | 443.1187 | 7.5590086 |  |  |  |
| Methyl (3b,11x)-3-Hydroxy-8-oxo-6-eremophilen-12-oate | 280.1654 | 13.604003 |  |  |  |
| Methyl 3b,24-dihydroxy-11,13(18)-oleanadien-30-oate | 484.3526 | 5.7769933 |  |  |  |
| MG(22:6(4Z,7Z,10Z,13Z,16Z,19Z)/0:0/0:0) | 402.275 | 15.443979 |  |  |  |
| Mulberrofuran P | 574.1235 | 8.619008 |  |  |  |
| N-(1-Deoxy-1-fructosyl)tryptophan | 366.1425 | 5.4580064 |  |  |  |
| N-Acetyl-DL-serine | 147.0526 | 0.65699995 |  |  |  |
| N-Acetyldjenkolic acid | 296.0495 | 1.5559999 |  |  |  |
| N-cis-octadec-9Z-enoyl-L-Homoserine lactone | 365.2903 | 15.139014 |  |  |  |
| N-cis-tetradec-9Z-enoyl-L-Homoserine lactone | 309.2289 | 13.660009 |  |  |  |
| N-Formylmethionyl-Leucylphenylalanine | 437.1995 | 8.166009 |  |  |  |
| N-Oleoyl-L-Serine | 369.2888 | 13.005004 |  |  |  |
| N-palmitoyl serine | 343.273 | 12.556021 |  |  |  |
| N-stearoyl serine | 371.3049 | 13.37902 |  |  |  |
| N-stearoyl serine Esi+13.376985 | 371.305 | 13.376985 |  |  |  |
| N2-Fructopyranosylarginine | 336.1656 | 0.63999957 |  |  |  |
| N6-Carbamoyl-L-threonyladenosine | 412.134 | 6.851009 |  |  |  |
| Nicarbazin | 302.064 | 4.1910024 |  |  |  |
| Nicarbazin Esi+3.0130002 | 302.064 | 3.0130002 |  |  |  |
| Nicotine imine | 161.1053 | 0.649 |  |  |  |
| PA(17:0/0:0) | 424.2619 | 15.233008 |  |  |  |
| PAF C-16-d4 | 528.3911 | 15.171 |  |  |  |
| Palmyrolide A | 337.26 | 14.505004 |  |  |  |
| PC(18:3(6Z,9Z,12Z)/0:0) | 517.3172 | 15.036005 |  |  |  |
| PE(19:0/0:0) | 495.3361 | 15.036006 |  |  |  |
| PE(20:0/0:0) | 509.3491 | 15.296009 |  |  |  |
| PE(21:0/0:0) | 523.3668 | 15.5279875 |  |  |  |
| PE(22:1(11Z)/0:0) | 535.3639 | 15.411991 |  |  |  |
| PE(P-18:0/0:0) | 465.3223 | 14.941015 |  |  |  |
| PE(P-18:0/0:0) Esi+14.941015 | 930.6452 | 14.941015 |  |  |  |
| PE(P-18:0/0:0) Esi+15.724002 | 465.3225 | 15.724002 |  |  |  |
| Persenone A | 378.2763 | 15.557 |  |  |  |
| Phe4Cl-His-OH | 444.0884 | 2.1679995 |  |  |  |
| Phloretin 2'-O-glucuronide | 450.1158 | 8.421009 |  |  |  |
| Phthalocyanine | 514.1602 | 9.164998 |  |  |  |
| Pifithrin-μ | 181.0168 | 10.355993 |  |  |  |
| pitavastatin | 421.1669 | 8.046009 |  |  |  |
| pitavastatin Esi+7.5789924 | 421.1669 | 7.5789924 |  |  |  |
| pitavastatin Esi+8.056016 | 421.1673 | 8.056016 |  |  |  |
| Polyoxin B | 507.1432 | 7.843004 |  |  |  |
| Polystachin (flavone) | 466.1603 | 11.215984 |  |  |  |
| Pranlukast | 480.168 | 8.698995 |  |  |  |
| PS(17:2(9Z,12Z)/12:0) | 689.4249 | 15.205997 |  |  |  |
| Pyraclofos | 360.042 | 6.8590055 |  |  |  |
| Pyridoxamine-5'-Phosphate | 248.0532 | 9.192005 |  |  |  |
| Quinacridone | 312.0915 | 10.359006 |  |  |  |
| Quinoline | 129.0563 | 6.8710017 |  |  |  |
| Sertaconazole | 436.0009 | 11.213009 |  |  |  |
| Shoyuflavone C | 418.051 | 10.35499 |  |  |  |
| sn-glycero-3-phosphoethanolamine | 215.056 | 6.6929946 |  |  |  |
| Styrene | 104.0621 | 11.985991 |  |  |  |
| Sulfabenzamide | 276.0551 | 0.63999957 |  |  |  |
| Tegaserod | 301.1891 | 5.3640084 |  |  |  |
| Thr-Asp-OH | 356.0816 | 10.349 |  |  |  |
| Triacanthine | 203.117 | 0.7249991 |  |  |  |
| Triamcinolone acetonide sulfate | 514.1609 | 9.026989 |  |  |  |
| Trilobinol | 300.2055 | 15.287988 |  |  |  |
| Trp Ala Thr | 376.1754 | 6.5310044 |  |  |  |
| Tylosin | 915.5175 | 10.781005 |  |  |  |
| Tyr His Lys | 446.2261 | 9.267014 |  |  |  |
| Tyr Leu His | 431.2152 | 9.302992 |  |  |  |
| Umbelliprenin | 366.2185 | 14.423981 |  |  |  |
| Vaccenyl carnitine | 425.3518 | 14.134009 |  |  |  |
| Val Asp Glu | 361.1443 | 6.9240065 |  |  |  |
| YM-53601 | 336.1596 | 15.56201 |  |  |  |
| δ-12-PGD2 | 352.2228 | 12.994996 |  |  |  |

**Table S3.** Metabolites measured in epithelial effluent on D0 and D6, related to PCA in Figure 2E; HILIC modes.

| **HILIC Negative** | | | **HILIC Positive** | | |
| --- | --- | --- | --- | --- | --- |
| Compound (ID) | Mass | RT | Compound (ID) | Mass | RT |
| 92.0361@3.4479983 | 92.0361 | 3.4479983 | Gamma-glutamyl-Alanine | 217.1085 | 6.783006 |
| 141.0393@6.1510005 | 141.0393 | 6.1510005 | Ribavirin 5'-phosphate | 324.0502 | 6.7119913 |
| 155.1048@6.8090067 | 155.1048 | 6.8090067 | Dihydroaceanthrylene | 204.0942 | 4.4359994 |
| 193.0553@1.2699999 | 193.0553 | 1.2699999 | 346.0316@6.7080064 | 346.0316 | 6.7080064 |
| 193.056@0.7649995 | 193.056 | 0.7649995 | N-Ornithyl-L-taurine | 239.0922 | 6.783007 |
| 218.0194@3.4490032 | 218.0194 | 3.4490032 | Allopurinol | 136.042 | 1.4819987 |
| 230.0869@6.163007 | 230.0869 | 6.163007 | 128.0956@7.200008 | 128.0956 | 7.200008 |
| 260.055@4.299994 | 260.055 | 4.299994 | (3beta,8beta)-3-Hydroxy-7(11)-eremophilen-12,8-olide | 250.1577 | 7.8430057 |
| 280.0455@1.326 | 280.0455 | 1.326 | 8-Acetylegelolide | 308.1286 | 6.911007 |
| 295.9963@7.854992 | 295.9963 | 7.854992 | 188.1286@8.88599 | 188.1286 | 8.88599 |
| 308.1175@6.900008 | 308.1175 | 6.900008 | Alanyl-Glycine | 146.0734 | 6.591003 |
| 308.1189@5.4010005 | 308.1189 | 5.4010005 | Fenson | 267.9968 | 6.286007 |
| 323.9945@1.2699999 | 323.9945 | 1.2699999 | 379.1632@6.783006 | 379.1632 | 6.783006 |
| 325.9925@1.2699999 | 325.9925 | 1.2699999 | N-Acetyl-DL-serine | 147.0563 | 7.205005 |
| 354.0143@7.5169992 | 354.0143 | 7.5169992 | Ismine | 257.105 | 7.1519904 |
| 384.1518@4.3980045 | 384.1518 | 4.3980045 | Margrapine A | 401.1447 | 6.781996 |
| 399.1727@4.9330015 | 399.1727 | 4.9330015 | Tylosin | 915.526 | 0.62099993 |
| 426.0334@7.4130096 | 426.0334 | 7.4130096 | 4-formyl Indole | 145.0547 | 4.434998 |
| 438.125@1.3049989 | 438.125 | 1.3049989 | 5-Nitro-ortho-anisidine | 168.0541 | 6.5919943 |
| 453.0955@7.4860077 | 453.0955 | 7.4860077 | 159.1638@1.0559999 | 159.1638 | 1.0559999 |
| 504.1844@6.7990055 | 504.1844 | 6.7990055 | coumarin-SAHA | 346.1528 | 7.4400015 |
| 513.1299@6.1209903 | 513.1299 | 6.1209903 | 1-nitroheptane | 145.1126 | 6.9160028 |
| 527.1692@4.840003 | 527.1692 | 4.840003 | 662.4576@0.5889997 | 662.4576 | 0.5889997 |
| 555.3508@0.81099933 | 555.3508 | 0.81099933 | (Z)-3-(1-Butenyl)-1H-2-benzopyran-1-one | 200.0818 | 6.781004 |
| 574.1175@6.7260084 | 574.1175 | 6.7260084 | Hirsutidin 3-glucoside | 507.1485 | 4.3320003 |
| 616.1276@6.737 | 616.1276 | 6.737 | Levofloxacin | 361.1513 | 0.8109994 |
| (2R,2'R)-3,3'-disulfanediylbis(2-acetamidopropanoic acid) | 324.043 | 7.1959963 | L-Homoserine lactone | 101.05 | 7.205005 |
| (R)-(+)-2-Pyrrolidone-5-carboxylic acid | 129.0419 | 6.5659933 | 254.0962@6.778995 | 254.0962 | 6.778995 |
| (R)-(+)-2-Pyrrolidone-5-carboxylic acid Esi-7.2009926 | 129.0414 | 7.2009926 | Cytosine | 111.0436 | 2.806 |
| 12R-HOME(13Z) | 298.2493 | 0.72000074 | quinaldine | 143.0744 | 4.4359994 |
| 15-oxo-hexadecanoic acid | 270.2192 | 0.73000103 | 1-[(5-Amino-5-carboxypentyl)amino]-1-deoxyfructose | 308.1646 | 8.021987 |
| 2-(Arabinosylamino)-3-(glucosylamino)propanenitrile | 379.134 | 6.1719975 | 330.1114@6.90999 | 330.1114 | 6.90999 |
| 2-Formyloxymethylclavam | 171.0501 | 6.138994 | 361.2137@1.7189987 | 361.2137 | 1.7189987 |
| 3-(N-Nitrosomethylamino)propionitrile | 113.0588 | 1.6069983 | HMMF | 89.0503 | 6.1909933 |
| 3-Ethenyl-4-hydroxy-2,5-dimethylhex-5-en-2-yl acetate | 212.1402 | 0.5670007 | 362.0066@6.7100077 | 362.0066 | 6.7100077 |
| 3-Hydroxy-3-methyl-glutaric acid | 162.0508 | 7.0209966 | 364.1568@0.93299955 | 364.1568 | 0.93299955 |
| 3-Hydroxymethylantipyrine | 204.0891 | 4.4219975 | 308.1253@1.0589995 | 308.1253 | 1.0589995 |
| 3alpha,4,7,7alpha-Tetrahydro-1H-isoindole-1,3(2H)-dione | 151.0632 | 1.059999 | 190.9784@5.5 | 190.9784 | 5.5 |
| 4,5-Dihydroxyphthalate | 198.0133 | 1.768002 | 223.1101@4.4099994 | 223.1101 | 4.4099994 |
| 4-hydroxy-2-oxo-Heptanedioic acid | 190.0476 | 3.7749958 | N-(1-Deoxy-1-fructosyl)histidine | 317.1292 | 7.18599 |
| 5-Aminoimidazole-4-carboxamide-1-β-D-ribofuranosyl 5'-monophosphate | 338.056 | 6.591003 | 541.1563@6.7629995 | 541.1563 | 6.7629995 |
| 6,8-Dihydroxypurine | 152.0335 | 1.6940001 | 5-Amino-6-(4-hydroxy-2-butenoyl)-2,2-dimethyl-4-chromanone | 275.1163 | 7.513992 |
| 6-Acetyl-2,3-dihydro-2-(hydroxymethyl)-4(1H)-pyridinone | 169.074 | 1.059999 | 1-(1-Pyrrolidinyl)-2-propanone | 127.1012 | 2.125 |
| 6-HpOME(7E) | 314.2444 | 0.7330011 | Phe Arg Ala | 392.2213 | 3.6979983 |
| 8-hydroxymianserin glucuronide | 456.1909 | 6.808997 | 444.0958@7.252994 | 444.0958 | 7.252994 |
| Acetoxyacetone | 116.0471 | 1.1610001 | 383.1973@1.6369982 | 383.1973 | 1.6369982 |
| Ala Ala His | 297.1419 | 5.17 | 218.1837@0.59700006 | 218.1837 | 0.59700006 |
| Bismuth subsalicylate | 361.9984 | 6.710999 | Tylosin Esi+0.71499884 | 915.5228 | 0.71499884 |
| D-Aspartic acid | 133.0363 | 7.1669936 | 368.0129@6.7020097 | 368.0129 | 6.7020097 |
| De-O-methylsimmondsin | 361.1365 | 5.2789955 | 504.2013@6.7839975 | 504.2013 | 6.7839975 |
| Decyl isobutyrate | 228.2084 | 0.6930006 | P-Fluorophenylalanine | 183.068 | 8.633992 |
| Dehydroxymethylflazine | 278.064 | 0.7939992 | 361.2137@1.5899976 | 361.2137 | 1.5899976 |
| Dihydro-2-methyl-3(2H)-furanthione | 116.0278 | 3.4519975 | 148.0185@0.55500025 | 148.0185 | 0.55500025 |
| E3040 | 299.1072 | 6.806001 | 383.1958@1.7299995 | 383.1958 | 1.7299995 |
| Ethyl aconitate | 202.0438 | 3.4479983 | Triamcinolone acetonide sulfate | 514.17 | 3.795004 |
| F-Honaucin A | 188.0495 | 3.4509957 | 616.1436@6.8280025 | 616.1436 | 6.8280025 |
| Fenothiocarb sulfoxide | 269.1092 | 6.1760073 | N-Desmethyltolmetin | 243.085 | 2.8080037 |
| Fentrazamide | 349.1257 | 6.1510005 | 161.017@6.711992 | 161.017 | 6.711992 |
| Ferulic acid | 194.057 | 1.2699999 | Butenafine | 317.2105 | 10.567008 |
| Gamma-glutamyl-Alanine | 217.1046 | 6.806014 | 278.0467@6.711992 | 278.0467 | 6.711992 |
| Glucosereductone | 88.0152 | 7.869007 | 308.1257@7.2080064 | 308.1257 | 7.2080064 |
| Glucosereductone Esi-1.8220024 | 88.0162 | 1.8220024 | 221.0532@0.972 | 221.0532 | 0.972 |
| Glucosereductone Esi-6.668012 | 88.0155 | 6.668012 | 453.1072@7.4639926 | 453.1072 | 7.4639926 |
| Glutamyl-Cysteine | 250.0615 | 2.7640045 | 616.1395@6.818987 | 616.1395 | 6.818987 |
| Glutarimide | 113.0463 | 7.2000084 | Methyl acrylate | 86.0382 | 6.9219975 |
| HMMF | 89.0476 | 6.1569963 | 385.0868@6.1859922 | 385.0868 | 6.1859922 |
| Haplopappin | 434.134 | 7.1909924 | Metyrapone | 226.1096 | 7.7670135 |
| His Cys Ala | 329.1138 | 4.690002 | 2-Protocatechoylphloroglucinolcarboxylate | 306.0411 | 6.710999 |
| His Gly Ser | 299.1197 | 6.177993 | Kamahine C | 268.1312 | 0.5849993 |
| Histidinyl-Aspartate | 270.094 | 4.0169997 | famprofazone | 377.2471 | 1.0329992 |
| Imiquimod | 240.1345 | 0.6569997 | Acetylagmatine | 172.1364 | 7.8060155 |
| Isoferuloyl C1-glucuronide | 370.0896 | 4.194 | Thr Cys Gly | 279.0898 | 7.11401 |
| Methyl acrylate | 86.0363 | 7.3820066 | 113.1216@1.0520015 | 113.1216 | 1.0520015 |
| Methyl hydrogen fumarate | 130.0257 | 7.3820066 | (R)-(+)-2-Pyrrolidone-5-carboxylic acid | 129.045 | 7.205005 |
| Methyl levulinate | 130.0629 | 0.97699904 | Panaquinquecol 5 | 202.1345 | 6.5729985 |
| Metsulfovax | 232.0649 | 6.492994 | Phosphodimethylethanolamine | 169.0548 | 4.4359994 |
| N-(1-Deoxy-1-fructosyl)alanine | 251.0972 | 6.6740065 | Scutellarein 7-glucuronosyl-(1-2)-glucuronide | 638.1253 | 6.8239913 |
| N-(1-Deoxy-1-fructosyl)threonine | 281.1068 | 6.625 | 5-Methoxy-3-hydroxyanthranilate | 183.0551 | 6.781004 |
| N-Acetyl-5-hydroxysulfapyridine | 307.0641 | 1.327 | CGP 52608 | 244.0454 | 6.1810117 |
| N-Acetyl-DL-serine | 147.0521 | 7.1950064 | 83.0392@7.2039924 | 83.0392 | 7.2039924 |
| N-Acetyldjenkolic acid | 296.0458 | 6.8909945 | 157.1485@2.2090018 | 157.1485 | 2.2090018 |
| N-Acetylneuraminic Acid | 309.1057 | 5.422994 | 382.1032@6.718004 | 382.1032 | 6.718004 |
| N-Acetylsulfadiazine | 292.0628 | 3.434998 | 5,5-Dimethyl-2(5H)-furanone | 112.0534 | 0.5830001 |
| N-Nitroso-N-methylurethane | 132.0527 | 6.572007 | 4-(Trimethylammonio)but-2-enoate | 144.103 | 5.164001 |
| Norfuraneol | 114.0317 | 4.334005 | 276.2338@0.59199935 | 276.2338 | 0.59199935 |
| O-Acetylethanolamine | 103.0626 | 7.1950064 | Tiopronin | 163.0326 | 6.7130084 |
| Pseudouridine | 244.0661 | 2.2359958 | N-(1-Deoxy-1-fructosyl)alanine | 251.1064 | 6.6939874 |
| Pseudouridine Esi-1.3209993 | 244.0687 | 1.3209993 | Mammea B/AC | 358.1777 | 4.807005 |
| Pterolactam | 115.0628 | 5.375996 | 2-Hydroxy-3-carboxybenzalpyruvate | 236.0343 | 6.7130084 |
| SB 228357 | 431.1275 | 6.130996 | 356.0872@0.78099895 | 356.0872 | 0.78099895 |
| Saxitoxin | 299.1356 | 1.0309991 | O-Demethylfonsecin | 276.0623 | 1.25 |
| Sulfamethazine | 278.0795 | 3.3169947 | Clausarinol | 414.2039 | 3.752004 |
| Theobromine | 180.0652 | 3.444 | N-Acetyldjenkolic acid | 296.056 | 6.8959966 |
| Undecyl isobutyrate | 242.2247 | 0.677 | Decanoyl m-Nitroaniline | 292.1812 | 7.8060155 |
| Val Met Phe | 395.1872 | 4.3989935 | 2094.0605@5.8179946 | 2094.0605 | 5.8179946 |
| Velcorin | 134.0201 | 7.409989 | 2090.9773@8.09001 | 2090.9773 | 8.09001 |
| Zidovudine glucuronide | 443.1277 | 4.638 | 1-Methyl-4-phenyl-1,2,3,6-tetrahydropyridine N-oxide | 189.1138 | 7.6599884 |
| apo-[3-methylcrotonoyl-CoA:carbon-dioxide ligase (ADP-forming)] | 173.1153 | 6.8070045 | 504.1336@6.1829977 | 504.1336 | 6.1829977 |
| beta-Alaninamide | 88.0626 | 6.804995 | 424.1367@4.851995 | 424.1367 | 4.851995 |
| formyl 14-methyl-8E-hexadecenoate | 282.2539 | 0.6589993 | 623.2596@7.802001 | 623.2596 | 7.802001 |
| methoxyacetic acid | 90.0325 | 3.442995 | 294.9475@5.5109925 | 294.9475 | 5.5109925 |
| trans-Aconitate | 174.0152 | 7.3820066 | 386.9348@5.521993 | 386.9348 | 5.521993 |
| xi-3-Hydroxy-2-oxobutanoic acid | 118.0261 | 7.3150015 | Nα-Acetyl-L-arginine | 216.1273 | 7.806015 |
| γ-Glutamyl-β-aminopropiononitrile | 199.0943 | 6.8049946 | 686.2177@6.280011 | 686.2177 | 6.280011 |
|  |  |  | CMP-N-trimethyl-2-aminoethylphosphonate | 473.1225 | 7.117993 |
|  |  |  | NS-102 | 261.0764 | 6.791994 |
|  |  |  | 1602.0385@5.8179946 | 1602.0385 | 5.8179946 |
|  |  |  | Methyl 3b,24-dihydroxy-11,13(18)-oleanadien-30-oate | 484.3564 | 0.6330006 |
|  |  |  | Mulberrofuran P | 574.1328 | 6.7310047 |
|  |  |  | Capecitabine | 359.146 | 0.761001 |
|  |  |  | (3R,7R)-1,3,7-Octanetriol | 162.1266 | 0.6139992 |
|  |  |  | 364.0042@6.7100077 | 364.0042 | 6.7100077 |
|  |  |  | Gly-Tyr-OH | 346.0828 | 6.914992 |
|  |  |  | 476.2072@7.661004 | 476.2072 | 7.661004 |
|  |  |  | 329.2248@0.573 | 329.2248 | 0.573 |
|  |  |  | 1-Phenanthrol | 194.0753 | 6.8380084 |
|  |  |  | 1-[(5-Amino-5-carboxypentyl)amino]-1-deoxyfructose Esi+7.6489906 | 308.1629 | 7.6489906 |
|  |  |  | 507.1307@4.3320003 | 507.1307 | 4.3320003 |
|  |  |  | 644.2031@7.8020015 | 644.2031 | 7.8020015 |
|  |  |  | 350.0012@6.276993 | 350.0012 | 6.276993 |
|  |  |  | Daunorubicin | 527.1804 | 3.5520017 |
|  |  |  | Patuletin 3-rhamnoside-7-(3''',4'''-diacetylrhamnoside) | 708.1975 | 6.2830048 |
|  |  |  | ε-Rhodomycinone | 428.1175 | 4.6790004 |
|  |  |  | 182.1431@1.9010009 | 182.1431 | 1.9010009 |
|  |  |  | Daunorubicin Esi+4.8460035 | 527.1775 | 4.8460035 |
|  |  |  | Ascariadole epoxide | 184.1088 | 0.61299956 |
|  |  |  | 272.1069@6.911007 | 272.1069 | 6.911007 |
|  |  |  | Methyl 6-O-galloyl-beta-D-glucopyranoside | 346.0915 | 6.5689883 |
|  |  |  | (S)-a-Amino-2,5-dihydro-5-oxo-4-isoxazolepropanoic acid N2-glucoside | 334.1006 | 1.1259996 |
|  |  |  | 615.1593@6.1840057 | 615.1593 | 6.1840057 |
|  |  |  | 8-Acetylegelolide Esi+7.2620087 | 308.1253 | 7.2620087 |
|  |  |  | 582.8898@5.533997 | 582.8898 | 5.533997 |
|  |  |  | 526.1847@6.7919946 | 526.1847 | 6.7919946 |
|  |  |  | isoflurophate | 184.0684 | 7.445996 |
|  |  |  | Penicilloic G acid | 352.1138 | 4.840003 |
|  |  |  | 680.8677@5.5419936 | 680.8677 | 5.5419936 |
|  |  |  | 1734.7382@8.062008 | 1734.7382 | 8.062008 |
|  |  |  | 633.2422@7.4429946 | 633.2422 | 7.4429946 |
|  |  |  | 655.1038@6.710999 | 655.1038 | 6.710999 |
|  |  |  | Glutamyl-Arginine | 303.1602 | 8.526 |
|  |  |  | 549.1629@4.841999 | 549.1629 | 4.841999 |
|  |  |  | 520.33@0.6330006 | 520.33 | 0.6330006 |
|  |  |  | Salivaricin A | 688.2382 | 6.7850027 |
|  |  |  | N-Succinyl-L-diaminopimelic acid | 290.1155 | 6.9119916 |
|  |  |  | 697.1618@6.177994 | 697.1618 | 6.177994 |
|  |  |  | 882.3378@8.0230055 | 882.3378 | 8.0230055 |
|  |  |  | 1438.0304@5.820006 | 1438.0304 | 5.820006 |
|  |  |  | 585.1282@6.744004 | 585.1282 | 6.744004 |
|  |  |  | Arg Trp Gln | 488.2559 | 7.667 |
|  |  |  | 645.171@6.1929946 | 645.171 | 6.1929946 |
|  |  |  | 3-Isoxazolidinone | 87.0343 | 7.161991 |
|  |  |  | 808.191@6.179994 | 808.191 | 6.179994 |
|  |  |  | 686.2202@6.280011 | 686.2202 | 6.280011 |
|  |  |  | 264.0278@6.7130084 | 264.0278 | 6.7130084 |
|  |  |  | 727.1746@6.1920023 | 727.1746 | 6.1920023 |
|  |  |  | Ethyl aconitate | 202.0486 | 3.184998 |
|  |  |  | 838.2012@6.190002 | 838.2012 | 6.190002 |
|  |  |  | 779.1646@6.164001 | 779.1646 | 6.164001 |
|  |  |  | 728.2341@7.664993 | 728.2341 | 7.664993 |
|  |  |  | Asn Met Arg | 419.2065 | 0.93699896 |
|  |  |  | 415.2231@1.0039998 | 415.2231 | 1.0039998 |
|  |  |  | Alfuzosin | 389.2111 | 1.0650008 |
|  |  |  | Ser Ser Ser | 279.1081 | 0.97200006 |
|  |  |  | Desmethylglymidine | 295.0627 | 7.116011 |
|  |  |  | 336.1682@8.093005 | 336.1682 | 8.093005 |
|  |  |  | Apocynin A | 468.1035 | 7.682006 |
|  |  |  | Glaudine | 399.1717 | 1.6579999 |
|  |  |  | 419.2672@1.7330012 | 419.2672 | 1.7330012 |
|  |  |  | Dilazep | 604.3028 | 10.567994 |
|  |  |  | Ganoderic acid theta | 530.2879 | 5.359993 |
|  |  |  | 529.1317@4.3330007 | 529.1317 | 4.3330007 |
|  |  |  | 694.192@6.577008 | 694.192 | 6.577008 |
|  |  |  | 623.2575@8.098008 | 623.2575 | 8.098008 |
|  |  |  | D2PM | 253.1461 | 4.6050014 |

**Table S4**. GSEA of Hallmark gene sets on gene expression differences between stretched and not stretched organoids on-chip. N=5 independent doners.

|  | pval | padj | log2err | ES | NES | size |
| --- | --- | --- | --- | --- | --- | --- |
| Hm Epithelial Mesenchymal Transition | 0.00014694 | 0.00367349 | 0.51884808 | 0.44083093 | 1.65230454 | 182 |
| Hm Cholesterol Homeostasis | 0.01951873 | 0.19518728 | 0.35248786 | 0.44827367 | 1.47073698 | 70 |
| Hm Uv Resp Dn | 0.10596027 | 0.6164966 | 0.15524197 | 0.3411643 | 1.24506405 | 137 |
| Hm Kras Sig Dn | 0.11096939 | 0.6164966 | 0.1482615 | 0.31875637 | 1.20029038 | 192 |
| Hm Coagulation | 0.16059379 | 0.69444444 | 0.12443417 | 0.32767973 | 1.18500168 | 133 |
| Hm Myogenesis | 0.2962963 | 0.92592593 | 0.08312913 | 0.28885995 | 1.0838531 | 184 |
| Hm Spermatogenesis | 0.3480589 | 0.9635832 | 0.07707367 | 0.29341845 | 1.06031105 | 129 |
| Hm Il6 Jak Stat3 Sig | 0.35901163 | 0.9635832 | 0.07977059 | 0.3145295 | 1.06359863 | 82 |
| Hm Complement | 0.36616162 | 0.9635832 | 0.07147863 | 0.27726631 | 1.04311746 | 189 |
| Hm Angiogenesis | 0.72267537 | 1 | 0.05121844 | 0.29206767 | 0.83246052 | 31 |
| Hm Hypoxia | 0.7394636 | 1 | 0.03895565 | 0.23617768 | 0.88617999 | 184 |
| Hm Bile Acid Met | 0.74858757 | 1 | 0.04301732 | 0.24317263 | 0.8568326 | 107 |
| Hm Tnfa Sig Via Nfkb | 0.89016603 | 1 | 0.03013298 | 0.21500662 | 0.80992048 | 193 |
| Hm Pancreas Beta Cells | 0.9222395 | 1 | 0.03784877 | 0.22787297 | 0.67754614 | 38 |
| Hm Kras Sig Up | 0.98847631 | 1 | 0.02484939 | 0.18989792 | 0.71168821 | 183 |
| Hm Inflammatory Resp | 0.98863636 | 1 | 0.02407421 | 0.18906798 | 0.7113021 | 189 |
| Hm Estrogen Resp Late | 0.99489144 | 1 | 0.0243564 | 0.17758591 | 0.66895832 | 193 |
| Hm Heme Met | 0.99489144 | 1 | 0.0243564 | 0.18459042 | 0.695344 | 193 |
| Hm Androgen Resp | 1 | 1 | 0.0312053 | 0.14614203 | 0.5015908 | 95 |
| Hm Apoptosis | 1 | 1 | 0.02470863 | 0.1601592 | 0.59461707 | 154 |
| Hm Estrogen Resp Early | 1 | 1 | 0.02421535 | 0.15632959 | 0.58983657 | 196 |
| Hm G2m Checkpoint | 1 | 1 | 0.08653997 | -0.1363952 | -0.5928953 | 198 |
| Hm Glycolysis | 1 | 1 | 0.08628656 | -0.154953 | -0.6718947 | 193 |
| Hm Mitotic Spindle | 1 | 1 | 0.08653997 | -0.1615358 | -0.7021786 | 198 |
| Hm Mtorc1 Sig | 1 | 1 | 0.02407421 | 0.09424711 | 0.3550247 | 193 |
| Hm Tgf Beta Sig | 1 | 1 | 0.06238615 | -0.145661 | -0.5090601 | 53 |
| Hm Pi3k Akt Mtor Sig | 0.99344262 | 1 | 0.06928365 | -0.1782324 | -0.7023942 | 102 |
| Hm Interferon Gamma Resp | 0.99082569 | 1 | 0.08705159 | -0.1805945 | -0.7817574 | 192 |
| Hm Reactive Oxygen Species Path | 0.97727273 | 1 | 0.06307904 | -0.194347 | -0.6717266 | 49 |
| Hm Interferon Alpha Resp | 0.975 | 1 | 0.06783383 | -0.1836181 | -0.720754 | 95 |
| Hm E2f Targets | 0.96380091 | 1 | 0.08783126 | -0.1872578 | -0.8160926 | 199 |
| Hm Unfolded Protein Resp | 0.9566787 | 1 | 0.07608372 | -0.1929561 | -0.7691752 | 113 |
| Hm Dna Repair | 0.94650206 | 1 | 0.08359906 | -0.1929485 | -0.8067684 | 149 |
| Hm Protein Secretion | 0.91875 | 1 | 0.07078991 | -0.2031986 | -0.797613 | 95 |
| Hm Il2 Stat5 Sig | 0.85321101 | 1 | 0.09560315 | -0.203541 | -0.8810884 | 192 |
| Hm Myc Targets V1 | 0.77777778 | 1 | 0.1017139 | -0.2075188 | -0.903373 | 200 |
| Hm Peroxisome | 0.72727273 | 1 | 0.08504275 | -0.2274954 | -0.898746 | 103 |
| Hm Uv Resp Up | 0.68859649 | 1 | 0.10592029 | -0.2193626 | -0.9157781 | 154 |
| Hm Notch Sig | 0.61298701 | 1 | 0.08220549 | -0.2964426 | -0.906637 | 29 |
| Hm Apical Junction | 0.44495413 | 1 | 0.13959967 | -0.2326307 | -1.0070115 | 192 |
| Hm Myc Targets V2 | 0.29360465 | 0.92592593 | 0.13649044 | -0.3105021 | -1.0966444 | 58 |
| Hm Allograft Rejection | 0.29357798 | 0.92592593 | 0.17520405 | -0.2407796 | -1.0407987 | 191 |
| Hm Apical Surface | 0.18181818 | 0.6993007 | 0.17520405 | -0.3518062 | -1.1824404 | 44 |
| Hm P53 Path | 0.16666667 | 0.69444444 | 0.23439265 | -0.2561994 | -1.113248 | 195 |
| Hm Hedgehog Sig | 0.16442049 | 0.69444444 | 0.17978232 | -0.379539 | -1.2309591 | 36 |
| Hm Wnt Beta Catenin Sig | 0.09722222 | 0.6164966 | 0.24133998 | -0.3988376 | -1.3101533 | 39 |
| Hm Xenobiotic Met | 0.0696565 | 0.58047081 | 0.28780513 | -0.2953058 | -1.2764947 | 191 |
| Hm Adipogenesis | 0.00997106 | 0.12463829 | 0.3807304 | -0.3127518 | -1.3574977 | 196 |
| Hm Fatty Acid Met | 0.00235234 | 0.0392056 | 0.4317077 | -0.3560367 | -1.4863545 | 154 |
| Hm Oxidative Phosphorylation | 1.02E-07 | 5.10E-06 | 0.70497572 | -0.4416094 | -1.919628 | 198 |

**Table S5**. Summary Table of The Fold Change in Papp Values of Stretched/Not Stretched Chips.

|  | US | | UP | |
| --- | --- | --- | --- | --- |
|  | **Fold Change** | **P-Value** | **Fold Change** | **P-Value** |
| **D-2** | 1.16 | 0.394 | 1.06 | 0.863 |
| **D-1** | 1.06 | 0.863 | 1.01 | 0.965 |
| **D0** | 1.23 | 0.382 | 1.14 | 0.78 |
| **D1** | 0.77 | 0.724 | 0.49 | 0.535 |
| **D2** | 0.97 | 0.966 | 1.13 | 0.866 |
| **D3** | 1.77 | 0.397 | 1.32 | 0.813 |

**Table S6.** GSEA of Hallmark gene sets on fold changes of stretched vs not stretched gene expression between KRAS WT vs MUT organoid-chips. N=3 independent doners for KRAS WT organoid-chips; N=2 independent doners for KRAS MUT organoid-chips.

|  | pval | padj | log2err | ES | NES | size |
| --- | --- | --- | --- | --- | --- | --- |
| Hm Oxidative Phosphorylation | 1.37E-07 | 6.85E-06 | 0.69013246 | 0.38636022 | 1.9944946 | 199 |
| Hm Adipogenesis | 9.34E-07 | 2.34E-05 | 0.6594444 | 0.36720791 | 1.89840568 | 200 |
| Hm Fatty Acid Met | 1.35E-05 | 0.00016819 | 0.59332548 | 0.37263374 | 1.86189704 | 158 |
| Hm Androgen Resp | 9.68E-05 | 0.00067639 | 0.5384341 | 0.40511351 | 1.85448678 | 100 |
| Hm Uv Resp Up | 0.00027569 | 0.00153164 | 0.49849311 | 0.33383047 | 1.6680131 | 158 |
| Hm Heme Met | 0.00031694 | 0.00158468 | 0.49849311 | 0.31878267 | 1.64564127 | 199 |
| Hm Mtorc1 Sig | 0.00035527 | 0.00161484 | 0.49849311 | 0.31152291 | 1.61052323 | 200 |
| Hm Glycolysis | 0.00119634 | 0.00498475 | 0.45505987 | 0.30131372 | 1.55546189 | 199 |
| Hm Bile Acid Met | 0.00175914 | 0.0066005 | 0.45505987 | 0.34877053 | 1.63302374 | 111 |
| Hm Cholesterol Homeostasis | 0.00326202 | 0.0101938 | 0.4317077 | 0.38355183 | 1.66424669 | 74 |
| Hm Hypoxia | 0.0095996 | 0.02666557 | 0.3807304 | 0.27566971 | 1.42308063 | 199 |
| Hm Peroxisome | 0.01233697 | 0.03246572 | 0.3807304 | 0.31927989 | 1.47526763 | 104 |
| Hm Interferon Alpha Resp | 0.02795091 | 0.06135568 | 0.35248786 | 0.30642934 | 1.40562752 | 97 |
| Hm Complement | 0.02822361 | 0.06135568 | 0.35248786 | 0.25701051 | 1.32675686 | 199 |
| Hm Kras Sig Up | 0.07609544 | 0.12273459 | 0.28780513 | 0.24363567 | 1.25510293 | 198 |
| Hm Protein Secretion | 0.08 | 0.125 | 0.26166352 | 0.27338877 | 1.24872076 | 95 |
| Hm Interferon Gamma Resp | 0.0881459 | 0.13355439 | 0.26635066 | 0.23470644 | 1.21339443 | 200 |
| Hm Kras Sig Dn | 0.11343284 | 0.16393443 | 0.23112671 | 0.22606555 | 1.16443121 | 196 |
| Hm Spermatogenesis | 0.1147541 | 0.16393443 | 0.21925035 | 0.25081693 | 1.22007297 | 133 |
| Hm Xenobiotic Met | 0.11854103 | 0.16464032 | 0.2279872 | 0.22408861 | 1.158502 | 200 |
| Hm Notch Sig | 0.16666667 | 0.21367521 | 0.16693385 | 0.34998939 | 1.24756427 | 32 |
| Hm Coagulation | 0.18539326 | 0.22070626 | 0.17232434 | 0.23371004 | 1.13934269 | 138 |
| Hm Il6 Jak Stat3 Sig | 0.20779221 | 0.23807916 | 0.15524197 | 0.26126523 | 1.15923412 | 83 |
| Hm Apoptosis | 0.28366762 | 0.31518625 | 0.13802224 | 0.21493465 | 1.07996463 | 161 |
| Hm Reactive Oxygen Species Path | 0.48218527 | 0.51296306 | 0.09054289 | 0.24314413 | 0.96867297 | 49 |
| Hm Inflammatory Resp | 0.79447853 | 0.81069238 | 0.0772747 | 0.17111706 | 0.88335198 | 199 |
| Hm Pancreas Beta Cells | 0.92592593 | 0.92592593 | 0.05896945 | 0.18160072 | 0.68441352 | 39 |
| Hm Dna Repair | 0.53250774 | 0.55469556 | 0.06307904 | -0.2111185 | -0.9659116 | 149 |
| Hm Estrogen Resp Late | 0.36849926 | 0.40054267 | 0.07955647 | -0.2182038 | -1.0486114 | 200 |
| Hm Tnfa Sig Via Nfkb | 0.20950966 | 0.23807916 | 0.11284336 | -0.2349781 | -1.1292229 | 200 |
| Hm Angiogenesis | 0.17918089 | 0.21851328 | 0.13355495 | -0.3480009 | -1.2256458 | 36 |
| Hm Apical Surface | 0.17346939 | 0.21683673 | 0.13574094 | -0.3314941 | -1.2306001 | 44 |
| Hm Allograft Rejection | 0.12573964 | 0.1654469 | 0.1501698 | -0.2449855 | -1.1771629 | 198 |
| Hm Wnt Beta Catenin Sig | 0.12265758 | 0.1654469 | 0.16440576 | -0.3536946 | -1.3034872 | 42 |
| Hm Pi3k Akt Mtor Sig | 0.074883 | 0.12273459 | 0.20429476 | -0.295733 | -1.2922686 | 105 |
| Hm E2f Targets | 0.07429421 | 0.12273459 | 0.19991523 | -0.2579621 | -1.239676 | 200 |
| Hm Uv Resp Dn | 0.05417957 | 0.09674923 | 0.24133998 | -0.2870101 | -1.3114715 | 144 |
| Hm Myogenesis | 0.05082212 | 0.09411504 | 0.24504179 | -0.2675438 | -1.2815705 | 197 |
| Hm Il2 Stat5 Sig | 0.05029586 | 0.09411504 | 0.24504179 | -0.2658609 | -1.2774696 | 198 |
| Hm P53 Path | 0.04457652 | 0.08915305 | 0.26166352 | -0.2688333 | -1.2919191 | 200 |
| Hm Tgf Beta Sig | 0.04139073 | 0.08623068 | 0.28785712 | -0.3740299 | -1.4494734 | 54 |
| Hm Hedgehog Sig | 0.02591317 | 0.06135568 | 0.35248786 | -0.4521773 | -1.5925507 | 36 |
| Hm Apical Junction | 0.02359714 | 0.05899285 | 0.35248786 | -0.2868159 | -1.3738864 | 197 |
| Hm Unfolded Protein Resp | 0.00371487 | 0.0109261 | 0.4317077 | -0.3500308 | -1.5550663 | 113 |
| Hm Myc Targets V2 | 0.00265208 | 0.00884026 | 0.4317077 | -0.4428755 | -1.7374037 | 58 |
| Hm G2m Checkpoint | 0.00184814 | 0.0066005 | 0.45505987 | -0.3224741 | -1.549698 | 200 |
| Hm Myc Targets V1 | 0.00010822 | 0.00067639 | 0.5384341 | -0.3557797 | -1.7097532 | 200 |
| Hm Epithelial Mesenchymal Transition | 6.97E-05 | 0.00058088 | 0.5384341 | -0.3611741 | -1.7356767 | 200 |
| Hm Estrogen Resp Early | 4.13E-05 | 0.00041327 | 0.55733224 | -0.3678154 | -1.7675926 | 200 |
| Hm Mitotic Spindle | 4.35E-06 | 7.26E-05 | 0.61052688 | -0.387394 | -1.8606975 | 199 |

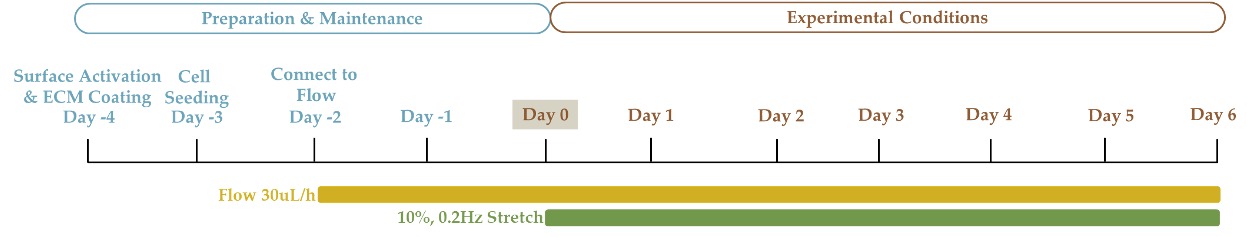

**Supplemental Figure 1.** **Experimental Timeline.** Detailed overview of key steps on critical days of organoid-on-chip preparation and experimental conditions.

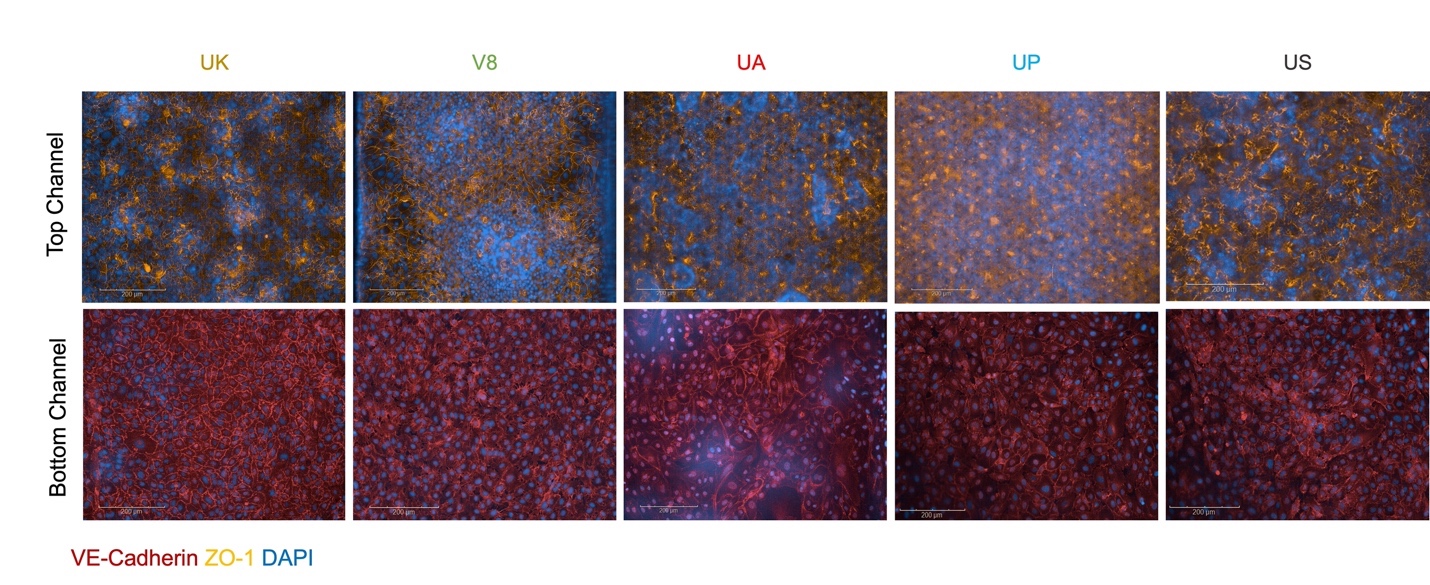

**Supplemental Figure 2**. **Tight junction formation in the organoid-on-chip.** Large scale confocal fluorescent images of the epithelial (top panel) and endothelial (bottom panel) channels of the organoid-chips on day 6 stained for tight junction protein ZO-1 (gold) and VE-Cadherin (red). Cell nuclei are labeled with DAPI (blue). Scale bars represent 200 μm.

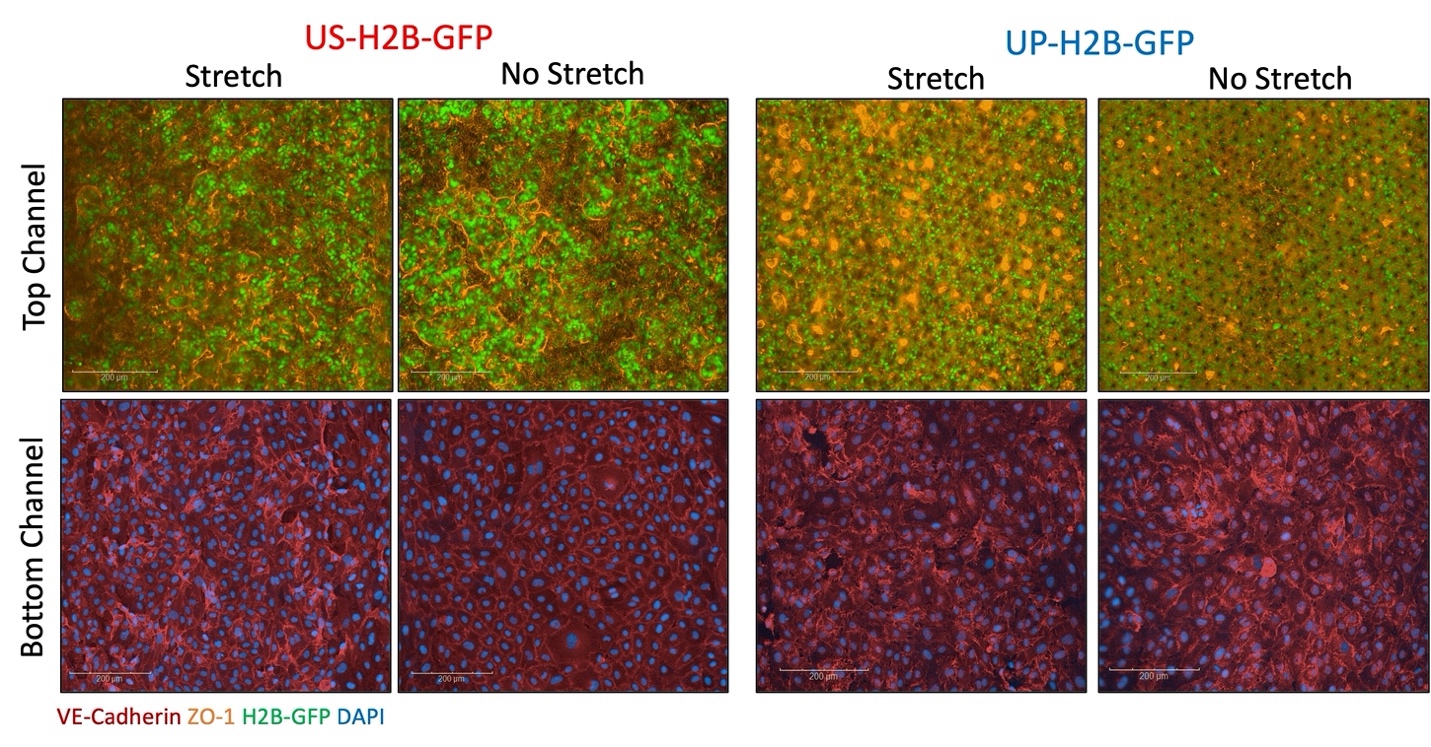

**Supplemental Figure 3. On-chip characterization of barrier function in response to peristalsis.** Representative confocal immunofluorescent images of the epithelial (top) and endothelial (bottom) channels of the US-H2B-GFP (left) and UP-H2B-GFP (right) organoids on-chip in the presence or absence of peristalsis stained for ZO-1 (gold) or VE-Cadherin (red) on day 6. Scale bars represent 200 μm.

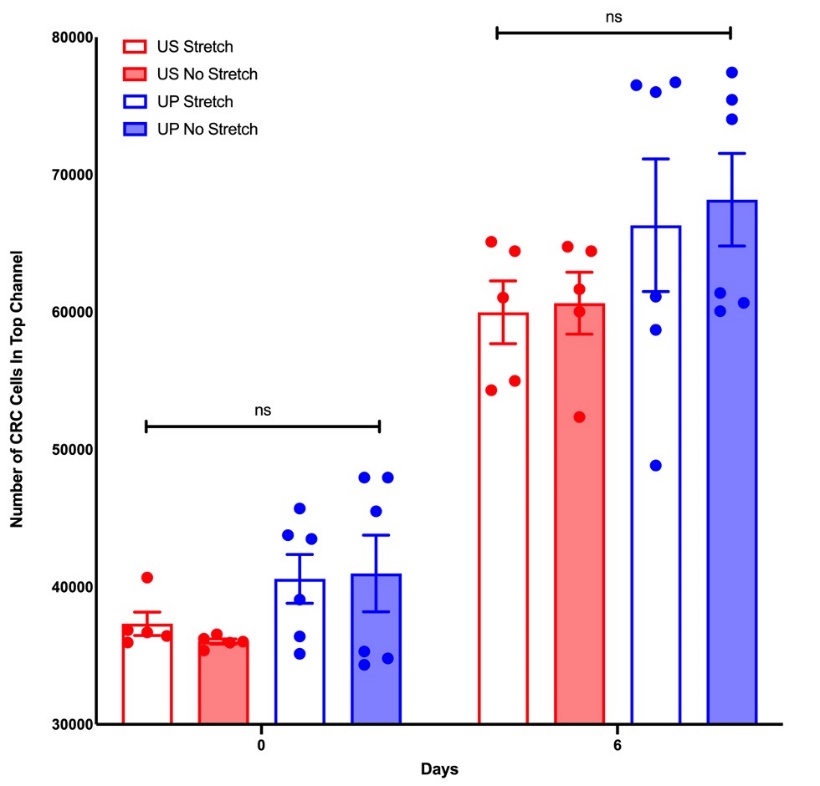

**Supplemental Figure 4. CRC Organoid growth in top channel in response to peristalsis.** Chips were imaged via fluorescent microscopy on days 0 and 6 and the number of GFP+ cells in the top channel for the US-H2B-GFP (red) and UP-H2B-GFP (blue) organoids was quantified. N=5-6 chips. Individual data are shown, with mean ± SEM represented and analyzed using a two-way ANOVA; p>0.05 for all comparisons on day 0 or day 6.

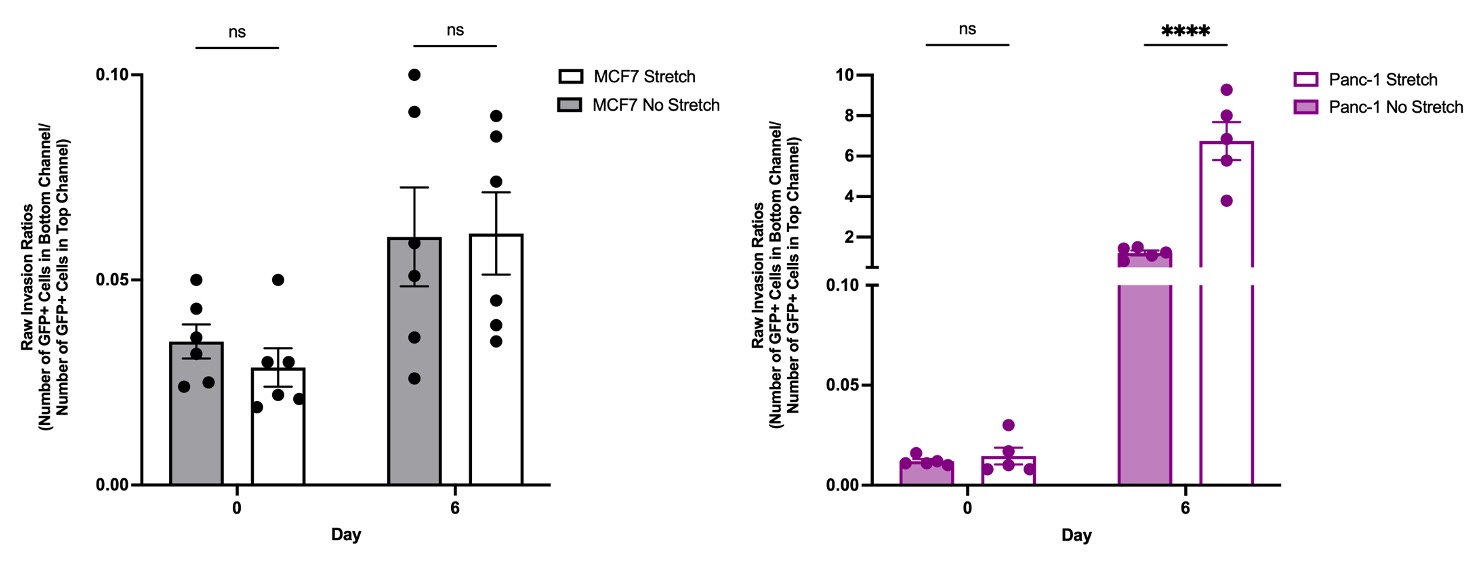

**Supplemental Figure 5.** **Tumor type invasion in response to peristalsis.** GFP+ MCF7 (left; grey) or Panc-1 (right; plum) tumor cell invasion was monitored over time by imaging the same chip regions at days 0 and 6. Invasion ratios were calculated for each imaged region as the number of GFP+ tumor cells in the bottom channel divided by the number of GFP+ tumor cells in the top channel at each timepoint. N=5-6 chips. Individual data are shown, with mean ± SEM represented and analyzed using a two-way ANOVA; ****p<0.0001.

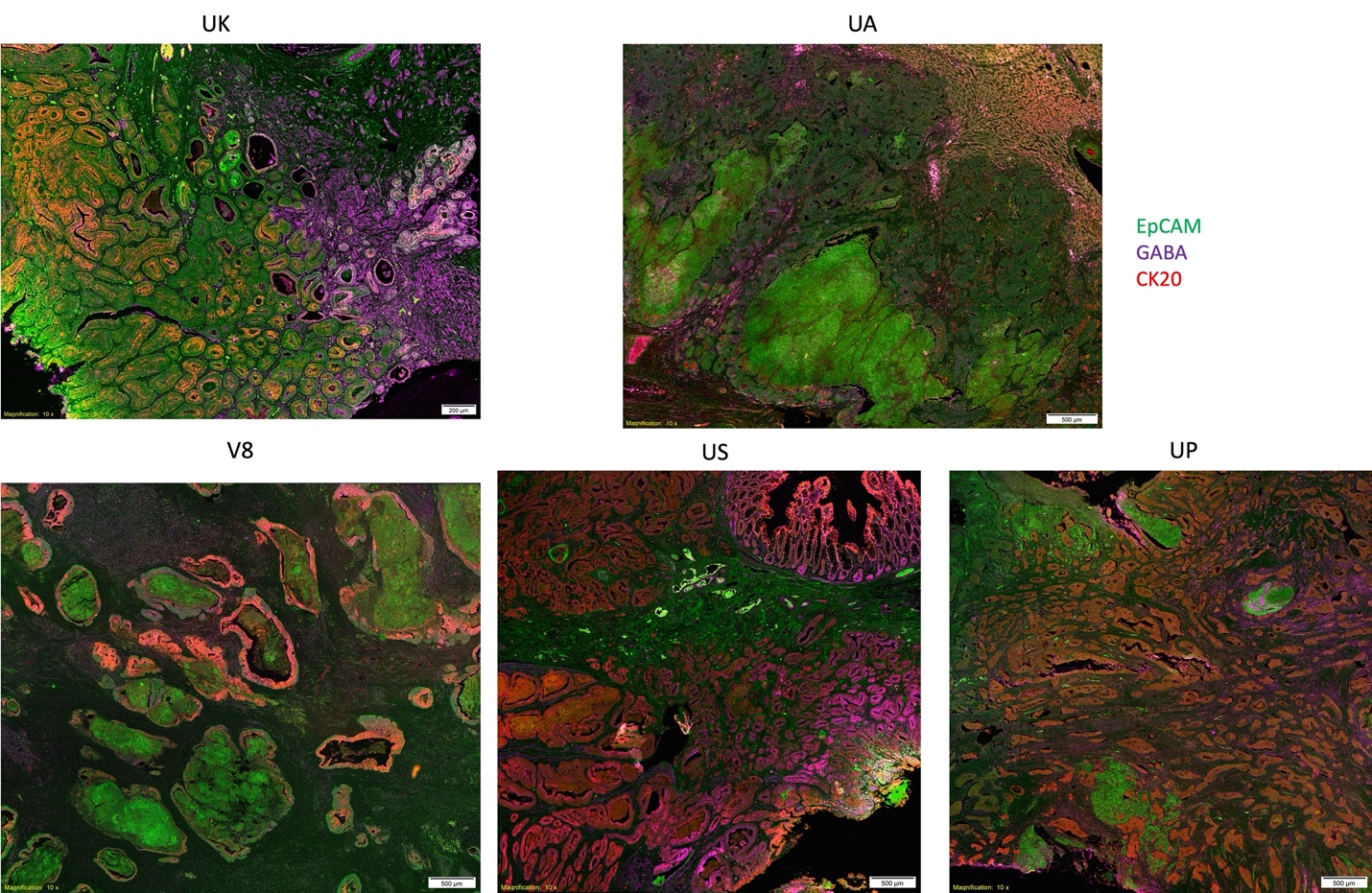

**Supplemental Figure 6. GABA in CRC tumors.** Un-cropped representative images of 10x immunofluorescence images of the 5 tumors stained for EpCam (green), CK20 (red), and GABA (purple). Scale bars represent 500 μm and 200 μm for UK.

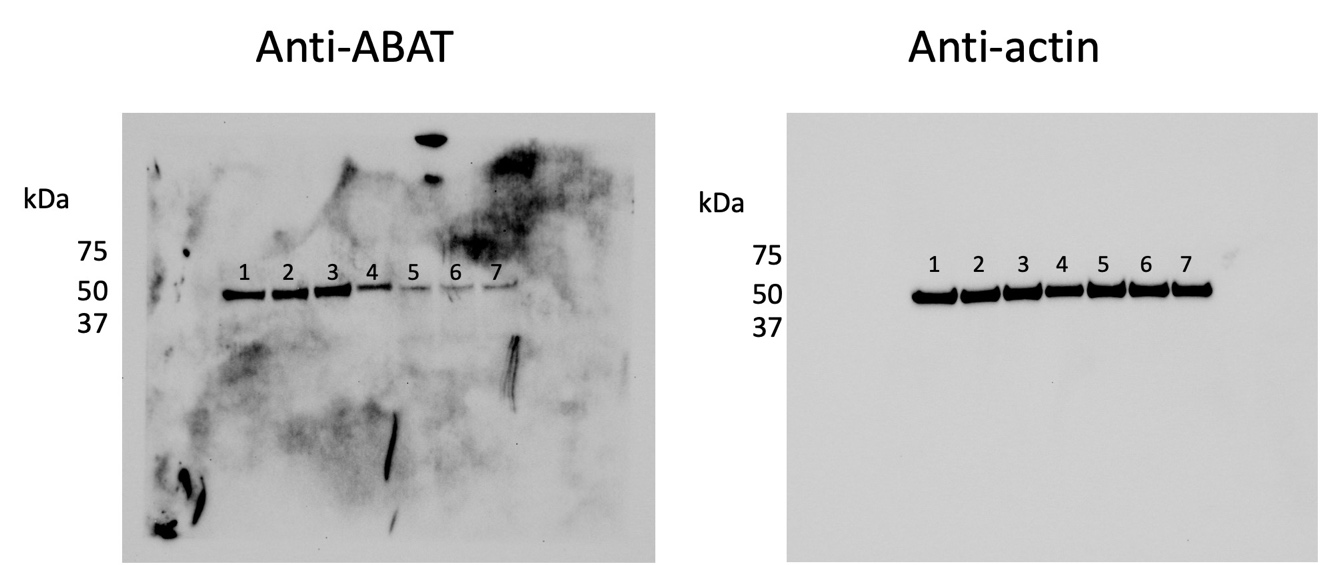

**Supplemental Figure 7. Confirmation of ABAT knockdown in HCT116 tumor cells.** Uncropped originals of the Western blots depicted in the manuscript. Protein expression for ABAT (left) and actin (right) was measured in HCT116 shRNA control MOI5 (lane 1), HCT116 shRNA control MOI2 (lane 2), HCT116 shRNA control MOI1 (lane 3), HCT116 no shRNA control (lane 4), HCT116 ABAT shRNA MOI5 (lane 5), HCT116 ABAT shRNA MOI2 (lane 6), or HCT116 ABAT shRNA MOI1 (lane 7). HCT116 shRNA control MOI5 tumor cells were chosen for further studies in the manuscript.
